# Nano-particles carried by multiple dynein motors: A Self-Regulating Nano-Machine

**DOI:** 10.1101/2020.07.09.194720

**Authors:** I. Fayer, G. Halbi, D. Aranovich, S. Gat, S. Bar, V. Erukhimovitch, Rony Granek, Anne Bernheim-Groswasser

## Abstract

Native cargos demonstrate efficient intra-cellular active transport. Here we investigate the motion of spherical nano-particles (NPs) grafted with flexible polymers, each ending with a nuclear localization signal peptide, thereby allowing recruitment of mammalian cytoplasmic dynein. Bead-motility assays show several unique motility features, depending on the number of NP-bound motors. NPs perform angular motion, in which the plus-end directed and right-handed motions are correlated. To simulate the system, we formulate a theoretical model that builds on single mammalian dynein properties, generalized to include motor-motor elastic and excluded-volume interactions. We find that long time trajectories exhibit both left- and right-handed helical motion, consistent with the measured angular velocity. The number of participating motors is self-regulated, thus allowing the NP to benefit from alternations between single and multiple transporting motors. Native cargos could use a similar approach to achieve both obstacle bypassing and persistent motion in the crowded cellular environment.

**Significance Statement:** The mechanism of active transport of native cargos, such as some viruses, is a long-standing conundrum. Their need for persistence motion towards the nucleus, while bypassing obstacles in the super-crowded intracellular milieu, requires sophisticated natural design. To fathom this machinery, we study a smartly designed nano-particle that recruits *several* dynein motor-proteins from the cytoplasm. Motility assays and model simulations reveal long run-times, long run-lengths, and helical motion around the microtubule symmetry axis. Moreover, the nano-particles self-regulate the number of dyneins participating in the motion, which optimizes its motility properties. We suggest that alternating between single motor motility, which we believe is beneficial for obstacle bypassing, and multiple motor states, which engender persistent motion towards the nucleus, the NP achieves optimal transport efficiency.

## Introduction

Active, motor-protein mediated transport is crucial for the intracellular conveying of a large variety of cargos in eukaryotes. Notably, microtubule-associated (MT-associated) motor proteins, dynein and kinesin, play here a cardinal role, hence, fueling a variety of vital biological processes^1–3^. While dynein is responsible for transport towards the cell center, members of the kinesin family are mostly responsible for transport towards the cell periphery^4–7^ The dynamic interplay between these two classes of motion orchestrates over the subcellular arrangement of organelles, e.g., mitochondria^8^ and Golgi-complexes^1^. In addition, different types of viruses have evolved to harness the dynein machinery for efficient targeting of the nucleus, where the infection process of the host cell takes place. HIV and herpes-simplex virus, for instance, express nuclear localization signal (NLS) peptides^7^ that recruit dynein from the cytoplasm^9–13^. Likewise, adenoviruses use other ligands (e.g., hexon) for engaging the active transport mechanisms^14,15^. Recently, preliminary theoretical work has been performed to mimic these viruses in rationally designed cargos that could be used for drug delivery applications^16^.

While knowledge of single dynein motility has been extensively increasing in the last decade^17–23^, much less is known about the collective motility of multiple motors as they carry a single cargo. Several observations lead to the prevailing belief that native cargos are carried by more than one motor^24–28^. Studies of supercoiled DNA plasmids enriched by NFκB receptors showed robust nuclear temporal localization^29^, supporting multiple dynein recruitment mediated by NLS binding. Complexes of fluorescently labeled single-stranded DNA, such as the VirE2 protein of Agrobacterium and the VirE2 itself, contain putative NLS regions that can recruit dynein^30^.

The collective behavior of motor proteins carrying a single cargo is often described in oversimplified terms such as cooperative or agonistic behavior (where motors walk at unison) *vs*. antagonistic (where motors interfere with each other walk)^31–35^. Nevertheless, this distinction is far from being clear cut. Moreover, recently, complex modes of motion have also been observed, in particular, a remarkable helical motion of dynein coated beads, demonstrating both right- and left-handed helices^36–38^. Notably, a theoretical description for the sideway motion of a *single yeast* dynein has been recently put forward^39^, in qualitative accord with these experiments.

Previous theoretical works modeled certain features of multi-motor motion^16,31,32,35,40–46^. Yet, these theoretical studies do not account for essential features for dynein stepping, e.g., they do not include the detailed locations of the dynein binding sites over the MT surface. In this spirit, as mentioned above, the *single motor* stepping was recently revisited to describe this two dimensional (2D) stepping of a single *yeast* dynein^39^. The 2D stepping model correctly accounts for the measured longitudinal step-size distribution and predicts a broad angular distribution of steps with small right-handed bias.

Motivated by a previous theoretical work^16^, we present here combined experimental-theoretical research on a rationally designed particle that, on the one hand, can serve as a model particle for native cargos and, on the other hand, allows quantitative examination of several transport characteristics. Our strategy is based on grafting a spherical NP with prescribed grafting density of Biotin-Polyethylene-glycol-thiol (Biotin-PEG-thiol) molecules and end-linking a controlled fraction of them with a single NLS, see Fig. 1A. It is expected, therefore, that under exposure to cell extract (CE), an NLS peptide will first recruit an α-importin protein, followed by binding of a β-importin protein, which subsequently will recruit dynein. Thus, the Biotin-PEG-thiol, which connects between the dynein and the NP, serves as a spacer polymer with variable flexibility (depending on its contour length, i.e., molecular weight). These spacers allow, by stretching, for motors originating from the NP sides to readily reach the MT surface (Fig. 2A-B), thereby increasing the number of motors participating in the transport. It follows that the NP characteristics, such as Biotin-PEG-thiol grafting density, linked NLS fraction, Biotin-PEG-thiol contour length, particle size, and CE concentration, all together govern over the number of dynein motors that bind to the MT simultaneously – which brings a great advantage for the use of this model cargo.

**Figure 1.**
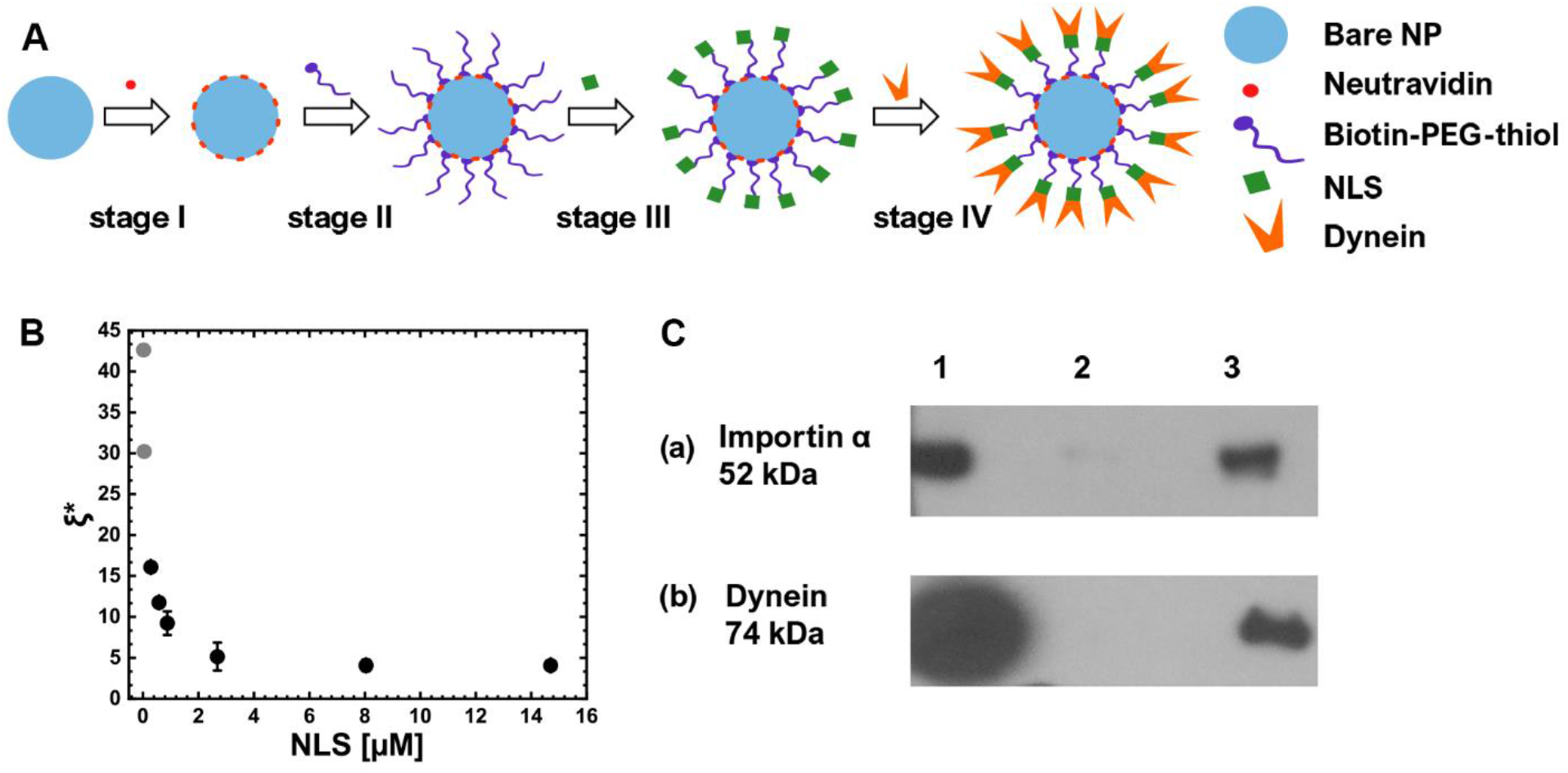
NPs synthesis and characterization. **(A)** NP synthesis process entails several consecutive steps, where at each step, a single component is added. **(B)** Mean anchoring distance between neighboring PEG- NLSs, ξ^*^, against the concentration of TAMRA-NLS (marked as NLS (x-axis) in the figure). The grey dots correspond to extrapolated values which were calculated from the fit of 〈*N*〉 *vs* [NLS] (see Table 1, SI Sec. 1 and Fig. S1.2). Error bars indicate the standard deviations for 3 experiments. **(C)** Western blot (WB) analysis results demonstrating the recruitment of importin-α (a) and *mammalian* Dynein motors (b) to the NPs after incubation in Hela cells extract. Group 1 (in both (a) and (b)) refers to Hela cells extract without NPs, group 2 refers to NPs coated with Biotin-PEG-thiol, and group 3 refers to PEG-NLS coated NPs. The concentration of cells extract used in WB is 3.4 mg/mL.

**Figure 2.**
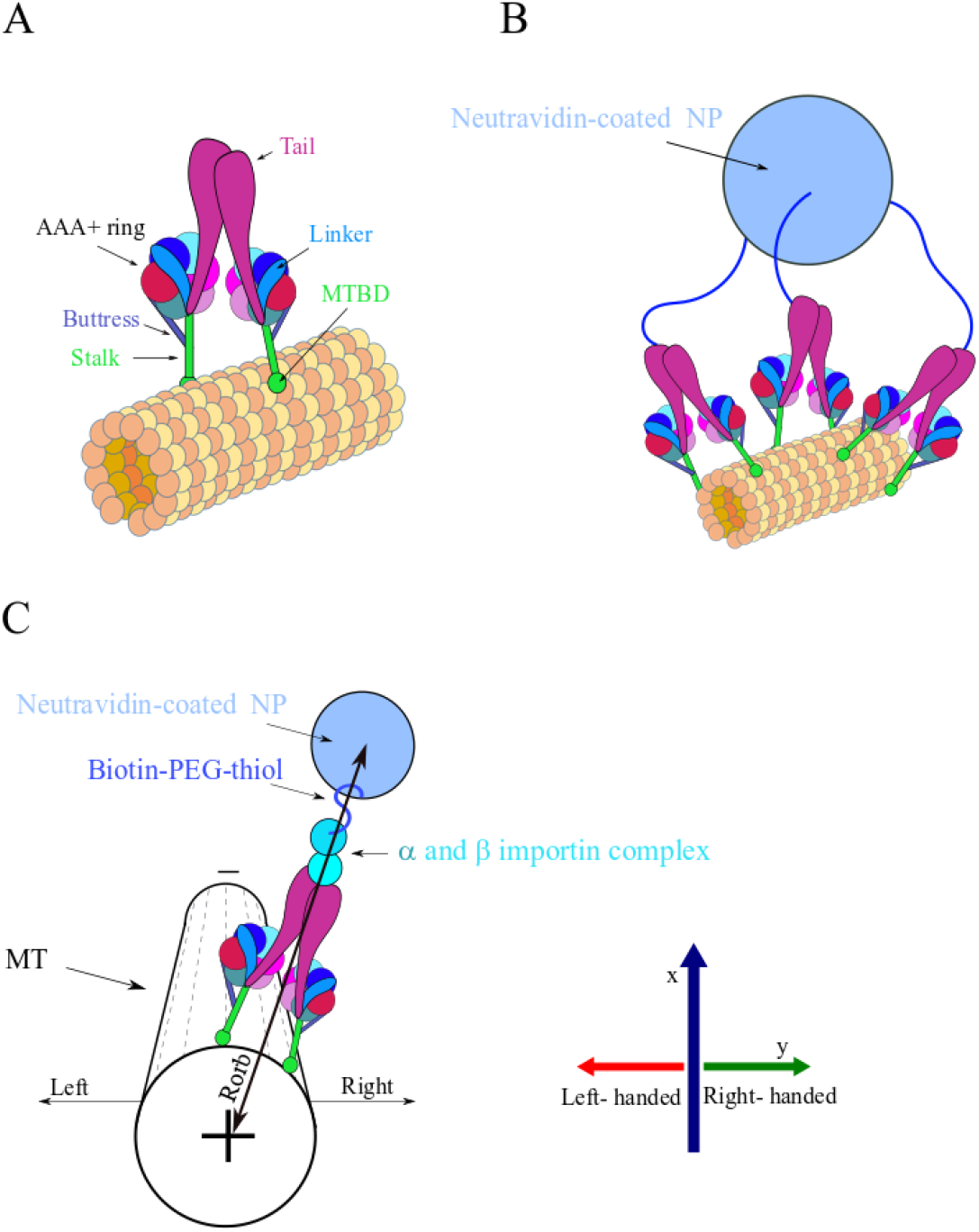
Illustration of single dynein and multi-motor NP (not to scale). Some biological factors, such as dynactin and adaptors, are not shown even though necessary for the motor activity. **(A)** *Left* – Single dynein motor-protein bound to the MT surface. The MT protofilaments and the corresponding *α* and *ß* tubulin subunits are represented by the collection of orange and yellow spheres. **(B)** A simplified illustration of a NP having three anchored PEG polymers over its surface, each of which connected to a single dynein. The Neutravidin-coated NP is represented by light-blue sphere and the PEG polymers are represented by the blue curved lines. **(C)** Illustration of the spatial orientations used throughout the article to characterize NP motion, i.e., longitudinal motion towards the MT plus- or minus-end and left or right transverse motion with respect to the MT long axis. The Neutravidin-coated NP color code is as in (B); the *α* and *ß* importin complex is also shown. The MT is presented with less details, where the dashed lines represent the MT protofilaments. Note that the longitudinal axis is denoted by *x* throughout the text and the transverse axis is denoted by *y*; positive *y* is the right direction and negative *y* is the left direction.

Our NPs are quite different from the cargos reported in Refs.^36–38^; the latter lack critical characteristics if they are to serve as model native cargos. First, they do not involve the native protein assembly for the recruitment of dynein from the cytoplasm. Second, their size is one order of magnitude larger than typical native cargo size, in the radius range 20 – 100 *nm*. Importantly, in this size range the drag force on the NP is entirely negligible^1^, implying that the single motor will move with its free-load velocity, and no load-sharing effect will appear^47^; such an effect will occur only for micron size NPs for which the drag force becomes significant^37,55^. Third, the cargos used in Refs^36,37,55^ lack control of mechanical coupling between motors. As such, they are less suitable for a fundamental experimental and theoretical study (of a model multi-motor cargo) that could lead to a profound understanding of the motion of native cargos.

In this work, we performed bead-motility assays using total internal reflection fluorescence (TIRF) microscopy. Single-particle tracking algorithms were used to extract the NPs’ trajectories. We measured different properties of the trajectories: (i) modes of motion, such as directional motion and hops between crossing MTs, (ii) run lengths, (iii) run times, and (iv) off-axis steps. On the theoretical side, the NP active transport is modeled by including a few competing processes, such as binding/unbinding to/from the MT surface and the stepping kinetics of individual dynein motor proteins. These are performed on the 2D curved microtubule surface, and are influenced by both the elastic coupling between motors (*via* the spacer polymers) and the excluded-volume interaction between motors. We used Monte-Carlo (MC) simulations to describe these dynamics. We then highlight the unique features of the multi-motor carried NP, based on a close comparison between experiment and theory (simulations), which gives mutual support to one another. Moreover, this comparison provides insight into the mechanism of NP motion, in particular, the role of multiple motor action during transport, and it suggests a possible mechanism for NP obstacle bypassing^51^.

## Results

### 1.1 NP preparation and analysis (experiment)

The NP synthesis process entails several consecutive steps, where at each step, a single component is added (Fig. 1A). Bare NPs saturated with Neutravidin were conjugated with Biotin-PEG-thiol spacers, then incubated for a short time in a solution containing SV40 T large antigen NLS peptide at variable concentrations.

The resulting NPs are being characterized (SI Sec.1) using cryo-transmission electron microscopy (cryo-TEM) (Fig. S1.1), dynamic light scattering (DLS), zeta potential (ζ-potential), and ultraviolet-visible (UVVis) adsorption isotherm, yielding the surface density and mean number of grafted Biotin-PEG-thiols, 〈*N*〉, that are end-conjugated by NLS (PEG-NLS) (Fig. S1.2). The latter can be transformed into a mean *anchoring* distance between neighboring PEG-NLS molecules, *ξ*^*^. The resulting anchoring distance, *ξ*^*^, is depicted in Fig. 1B against [NLS], showing that incubation of NPs in higher [NLS] solutions lead to a shorter distance between adjacent PEG-NLSs until it reaches saturation above [NLS] = 4 μM. Note that the theoretical value of the (free polymer) gyration radius is *R*_g_ = 2.27 nm (for a Biotin-PEG-thiol of *M*_w_ = 5 kDa; see SI Sec. 2). Thus, since Neutravidin diameter is about 5 *nm*^52^, and since the Neutravidins are closely packed on the NP surface, we may conclude that the anchored Biotin-PEG-thiol molecules are effectively in the so-called “mushroom regime”^53^, *ξ*^*^ > *R*_g_.

Next, NPs are incubated in (Hela) cell extract (CE), allowing the recruitments of α- and β- importins, followed by *mammalian* dynein association; the recruitments of the importins and dynein are verified *via* Western Blot (WB). Fig. 1C(a) shows the WB image after exposure to cell extract, demonstrating antibody specific binding to importin-α. Fig. 1C(b) shows the WB image demonstrating dynein recruitment. Group 1 (in both (a) and (b)) refers to CE without NPs; group 2 refers to NPs coated with Biotin-PEG-thiol incubated in CE; and group 3 refers to PEG-NLS coated NPs, which were also incubated in CE. Note that although importin-β binding was not directly verified, dynein recruitment requires importin-β binding, suggesting that, in the absence of NLS, the dynein machinery is not recruited to the NP surface.

An important issue regarding our NP construct is whether it allows non-specific binding of kinesin motors, which will influence the motility characteristics. Since the bare NP is covered with a first layer of Neutravidin, a second layer of BSA, and a third layer of Biotin-PEG-thiol (henceforth, PEG), all of which together are expected to efficiently passivate the NP surface, we can exclude the possibility of direct binding of kinesin to the NP.

### 1.2 Model and simulations (theory)

To explain the above experimental results and gain profound knowledge regarding the NP motion and behavior, we constructed a model for the active transport of the NP. Previous work on multi-motor complexes builds on a single motor motion on a 1D microtubule track^31,32,35,41,54^, and as such, cannot describe the motion of the NP around the 2D MT surface. The single motor stepping was recently revisited to describe 2D stepping of a single *yeast* dynein on the microtubule surface; this stepping model correctly accounts for the measured longitudinal step-size distribution and, moreover, predicts a broad angular distribution of steps (of a single motor) with a small right-handed bias^39^. More recently, Elshenawy *et al*.^37,55^ studied the single *mammalian* dynein stepping kinetics, providing the longitudinal- and transverse- step-size distributions. Using Yildiz co-workers raw data^37,55^, we adjusted our *yeast* dynein model^39^ to describe single *mammalian* dynein (see Materials and Methods, Model and simulation algorithm section).

The details of our model are described in the Materials and Methods (Model and simulation algorithm section). The model entails a NP on which polymers of equal contour length and fixed density – corresponding to a prescribed mean spacing *ξ^*^* – are grafted at random positions. Each free polymer-end is assigned with a dynein motor, such that *ξ* = *ξ^*^* by definition. Thus, 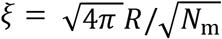, where *R* is the NP radius and *N*_m_ is the number of motors that are anchored to the NP surface. Dynein is assumed to perform binding/unbinding kinetics to/from the MT surface, competing with its stepping kinetics (see Methods section). Note that by the term “unbinding” we refer to the full detachment of the motor from the MT. This implies that its two microtubule binding domains (MTBDs) get disconnected from the MT, such that the inverse of the single motor unbinding rate defines its processivity time. Likewise, “binding” refers to the attachment of a motor to the MT surface from the bulk solution. The motors motion on the 2D curved microtubule surface is influenced by the elastic coupling between them (*via* the spacer polymers) – a coupling that influences both the motors step vector distribution and the binding-unbinding kinetics. In addition, we account for the excluded-volume interaction between motors that prevents them from stepping over each other, however, we do not account for the excluded volume interaction between different spacer polymers which we believe is negligible (see Methods section). We note again, consistent with the nm scale of the NP, that the drag force on the NP is negligible. We used Monte-Carlo (MC) simulations to describe the different competing processes, while readjusting the NP center-of-mass and rotational angle after each MC step to achieve mechanical force balance – processes that are orders of magnitude faster than the binding/unbinding and stepping processes (SI Sec. 7). The model allows investigation of the different microscopic internal states between which the NP fluctuates in time, in particular, the number of MT bound motors, *M*_B_, that participate in the motion at a given time.

We note that our model does not include a few experimental setup constraints and boundary conditions. First, it permits free motion of the NP around the MT surface (similar to the bridge-like setup of Yildiz and co-workers^36^), unlike our experimental setup in which the motion might be impaired due to the presence of the impenetrable glass surface. Second, it does not include MT ends (i.e., MTs are considered infinite) and MT junctions (i.e., only isolated MT tracks are considered).

### 1.3 NP motility assays (experiment)

Nano particles were washed from the excess cell extract, incubated in an ATP solution, and then were injected into a flow cell in which MTs are adsorbed and immobilized on a glass surface. We investigated the different modes of motion of the NPs, under different [NLS] and [CE], as defined in Table 1 (henceforth referred to as systems I, II, III, IV). Identical CE (i.e. extracts from an identical batch) was used in systems I and II, and another CE was used in systems III and IV. Since, in general, protein concentrations are likely to be different in different cell extracts we shall limit the comparison of motility assays results only within each pair, namely I *vs* II and III *vs* IV. (An additional system – system V – with high [CE] was examined. However, it is not shown or discussed here since its motility is strongly reduced presumably due to molecular crowding effects, see SI Sec.3, Table S3.1). We used total internal reflection fluorescence (TIRF) microscopy to follow the NPs motion (see Movie 1). First, we used marked minus-end MTs to confirm the expected correlation between the MT polarity and the NP motional direction – as dictated by dynein bias to move towards the minus-end (Fig. 3A). Next, a standard particle tracking algorithm was used to extract the NPs center-of-mass position with nm-scale precision^56,57^ (SI Sec. 4), resulting in time-dependent individual trajectories.

**Figure 3.**
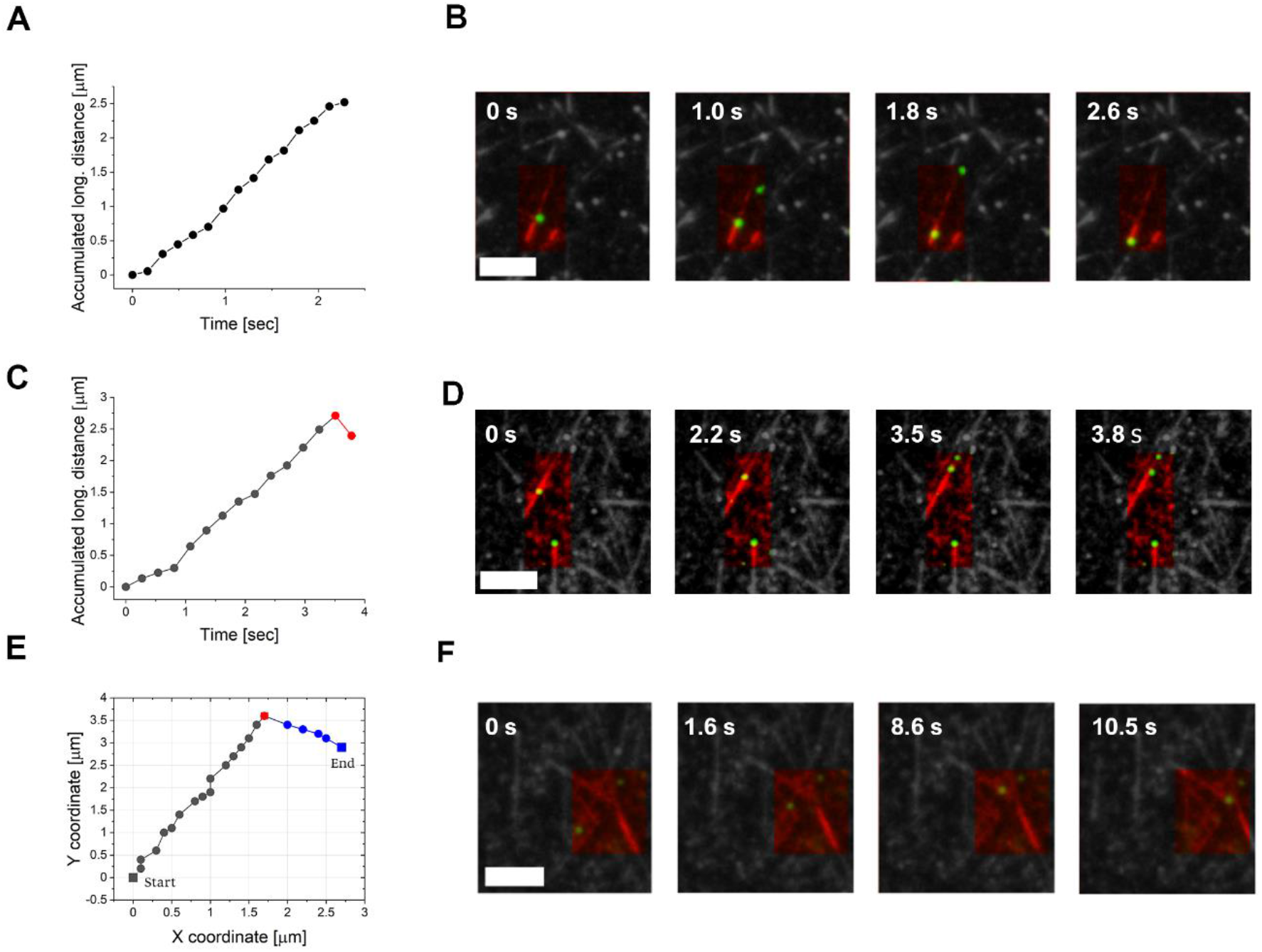
Examples of the NPs motion modes. Two first rows show the accumulated longitudinal distance of a NP and the corresponding snapshots for different motion modes. **(A-B)** Directed motion of a NP moving towards MT minus-end (marked-end MTs are shown). **(C-D)** Minus-end directed motion with a backward (plus-end) step at the end of the NP trajectory. **(E-F)** Trajectory of a NP hopping between crossing MT tracks. The MT tracks are colored in red and the NPs in green. Conditions: [*NLS*] = 0.05 μM and that of cells extract is 3.4 mg/mL (system III). The time interval between frames is 0.27 sec. The bare NPs mean diameter is 40 nm. (Bars are 5 μm).

**Table 1.** Summary of the systems studied and their basic properties: NLS and CE concentrations, estimated mean number 〈*A*〉 of PEG-NLS per NP, estimated mean anchoring distance *ξ*^*^, and fractions of NPs performing distinct modes of motion. An identical cell extract was used in systems I and II, and another extract was used in systems III and IV. The bare NPs mean diameter is 40 nm.

| System |  | I | II | III | IV |
| --- | --- | --- | --- | --- | --- |
| NLS conc. [$\mu\text{M}$] | | 0.025 | 0.05 | 0.05 | 0.3 |
| CE conc. [$\mu\text{M}$] | | 3.4 | 3.4 | 3.4 | 3.4 |
| $\langle N \rangle$ | | 5.2 | 10.4 | 10.4 | 36.6 |
| $\xi^*$ [nm] | | 42.6 | 30.2 | 30.2 | 16.1 |
| Motion mode |  | Fraction |  |  |  |
|  |  | I | II | III | IV |
| 1.1 | Minus-end directed motion – detachment before arrival to the end. | 0 | 0 | 0.74 | 0.72 |
| 1.2 | Minus-end directed motion, NP stuck when MT-end is reached. | 0.11 | 0.24 | 0.10 | 0.20 |
| 1.3 | Minus-end directed motion + a single backward (plus-end) directed step, when MT-end is reached or during the motion. | 0.87 | 0.52 | 0.10 | 0 |
| 1.4 | MT track crossing | 0.02 | 0.24 | 0.06 | 0.08 |

To analyze the NP motion, we differentiated the NPs trajectories into plus-end directed and minus-end directed motion, with respect to the movement along the MT long-axis (henceforth ‘longitudinal motion’) (Fig. 2C). We have detected three major modes of motion (Fig. 3 and Table 1): (i) NPs moving continuously towards the MT minus-end (Fig. 3A,B); (ii) NPs that have reached the MT-end after exhibiting continuous minus-end directed motion, followed by a single backward step (i.e., towards the MT plus-end) and detachment (Fig. 3C,D); sometimes the NP is remaining immobile for some time before detaching (not shown); and (iii) NPs traversing between crossing MTs (Fig. 3E-F, Movie 2). Noteworthy, the trajectories involving shifting between crossing MT tracks (mode (iii) above) are very unlikely to occur if the NP is carried by a single motor. Since such traversing events do occur, it is safe to assume that the NP is carried by more than one motor, even if only instantaneously.

We tested the influence of the [NLS] and [CE] on the NPs motion. In Table 1, we summarize the estimated values of the mean anchoring distance *ξ*\* of PEG-NLS for the four systems (deduced from the data presented in Fig. 1B and SI Sec. 1). Note that *ξ^*^* serves as an estimated lower bound for the mean anchoring distance ξ of the PEG-NLS-*αβ*-dynein (*αβ* refers to the *α* and *β* importin complex), such that *ξ* should approach *ξ^*^* (from above) as [CE] increases (*ξ* has not been measured). Moreover, given the expected sub-mM concentrations of dynein^58^, we assume the system is much below the saturation of dynein binding “isotherm,” such that increase of [CE] (hence dynein bulk concentration) is likely to increase dynein surface concentration, i.e., decrease of *ξ*. Thus, systems I and II, belonging to the same batch and having the same [CE], are expected to have the same value of *ξ/ξ^*^* or, equivalently, the same value 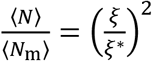; similarly for systems III and IV. Note that the value of *ξ^*^* determines the mean number of PEG-NLS per NP *via* 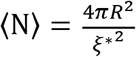. This implies that 〈*N*〉 varies between 5 and 37 for [NLS] varying between 0.025 and 0.3 μM (Table 1 and SI Sec.1).

#### Longitudinal velocity

The mean properties of the NP trajectories are shown in Fig. 4 and summarized in Table 2 as per system. Comparison within each of the pairs I-II and II-IV, we observe a significant decrease of the NP longitudinal velocities – from the single motor value – with increasing number of NP attached motors, i.e. *N*_m_ in system II being larger than in system I, and *N*_m_ in IV being larger than in III. If one assumes an increase in the mean number of motors participating in the transport 〈*M*_B_〉 with increasing *N*_m_ (as shown by our theoretical predictions in section 1.4), this suggests a decrease of the NP velocities on the increase in 〈*M*_B_〉, presumably due to inter-motor interactions. We emphasize that the regime of transport corresponds to *vanishing drag* due the nm size of the NPs, implying that no load-sharing effect whatsoever should be present; such a reduction of velocity with increasing number of participating motors is commonly observed in any standard motility assay of MTs moving on kinesin/dynein decorated glass surface^59,60^. Moreover, we have examined the average NP run-time, 〈*τ*_p_〉, and run length (along the MT symmetry axis), 〈*λ*〉, for the different systems (Fig. 4B and Table 2). We observe an increase of 〈*τ*_p_〉 with increasing system [NLS] and [CE] concentrations (i.e. decrease of *ξ* or increase of *N*_m_), consistent with the anticipated increase mean number of participating motors, 〈*M*_B_〉 (*c.f*. Sec. 1.4). Note, however, that since 〈*λ*〉 = 〈*ν*_x_ × *τ*_p_〉, the run length dependence on system concentrations (influencing ξ) is non-monotonous and shows a maximum for system IV (Tables 2 and S3.1 to include system V in the comparison).

**Figure 4.**
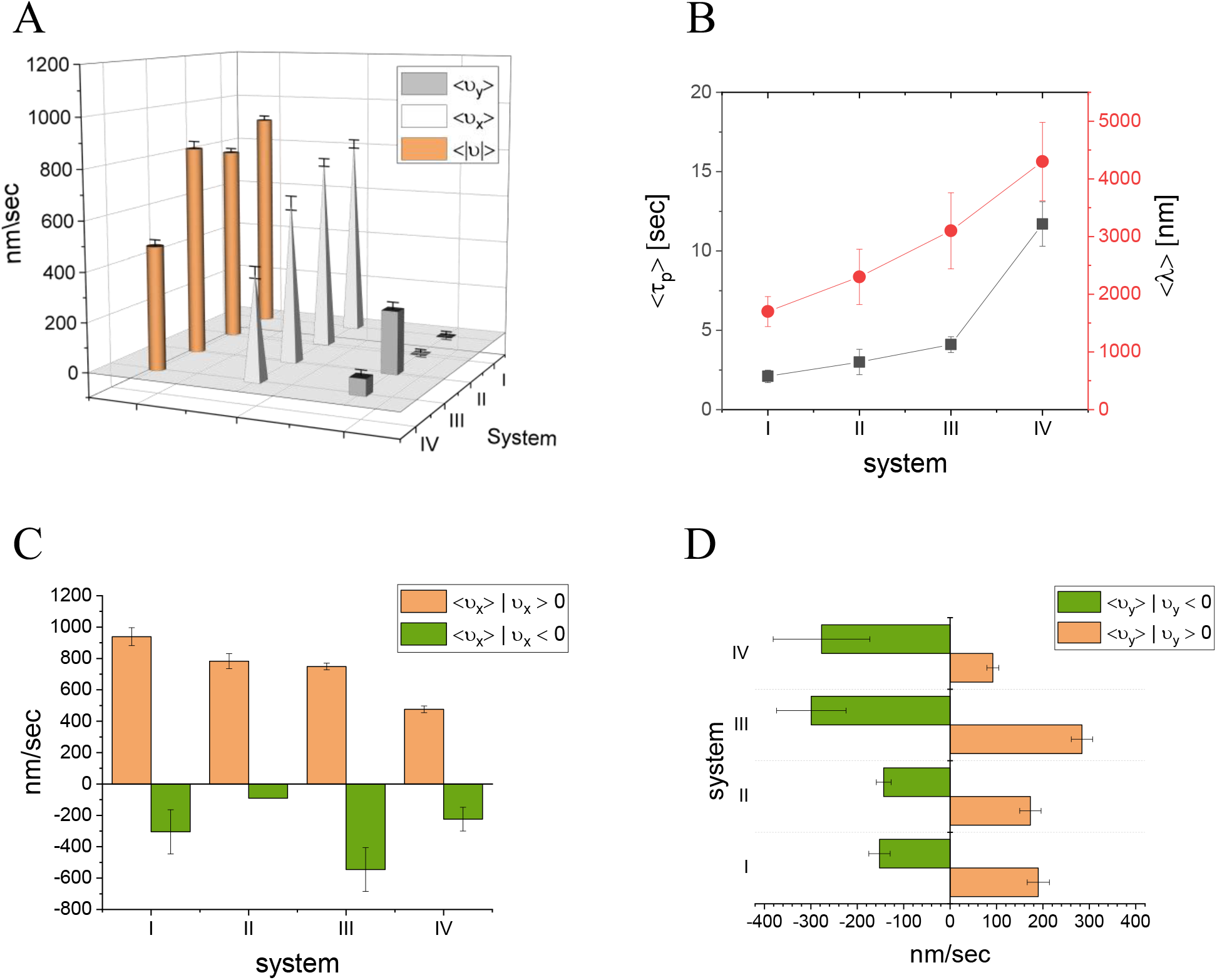
NPs experimental mean velocities, run-times, and run-lengths for systems I to IV. **(A)** Mean values of longitudinal, 〈*ν*_x_〉, transverse, 〈*ν*_y_〉, and absolute, 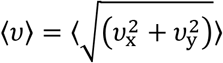, velocities. A minus-end directed motion corresponds to *ν*_x_ > 0, and a right-handed motion corresponds to *ν*_y_ > 0 (see Fig. 2C for spatial orientation). **(B)** Run-times and run-lengths; the run-time is measured from the time the NP binds to the MT until the time it unbinds from it; the run-length is the accumulated distance traveled along the MT symmetry axis; *ν*_x_ > 0 defines minus-end directed motion, and *ν*_x_ < 0 defines plus-end directed motion. **(C)** Mean longitudinal velocity evaluated separately for minus-end directed and plus-end directed motions. **(D)** Mean transverse velocity evaluated separately for right-handed and left-handed motions. Values correspond to mean ± SEM. The bare NPs mean diameter is 40 nm.

**Table 2.** NPs experimental mean velocities, run-times, and run-lengths for the NPs motion in systems I to IV. *ν* is the absolute velocity, *ν*_x_ is the longitudinal velocity, *ν*_y_ is the transverse velocity, *τ*_p_ is the processivity time, *λ*^*^ is the absolute accumulated distance that the NP covered regardless of the motion direction, and *λ* is the total longitudinal run length in the direction of the MT minus-end. Values correspond to mean ± SEM. The bare NPs mean diameter is 40 nm.

|  | I | II | III | IV |
| --- | --- | --- | --- | --- |
| [NLS] [$\mu\text{M}$] | 0.025 | 0.05 | 0.05 | 0.3 |
| $\langle N \rangle$ | 5.2 | 10.4 | 10.4 | 36.6 |
| [CE] [$\mu\text{M}$] | 3.4 | 3.4 | 3.4 | 3.4 |
| #steps | 80 | 62 | 362 | 134 |
| $\langle \vert v\vert \rangle$ [nm/sec] | 905±18 | 793 ± 16 | 842 ± 28 | 493 ± 23 |
| $\langle v_x \rangle$ [nm/sec] | 815±68 | 769 ± 49 | 642 ± 29 | 422 ± 26 |
| $\langle v_x \rangle \vert v_x > 0$ [nm/sec] | 939±57 | 783 ± 48 | 749 ± 21 | 476 ± 22 |
| $\langle v_x \rangle \vert v_x < 0$ [nm/sec] | -305±141 | -91 ± 0 | -545 ± 140 | -224 ± 76 |
| $\langle v_y \rangle$ [nm/sec] | 1.7 ± 25 | -0.07 ± 24 | 252 ± 23 | 67 ± 16 |
| $\langle v_y \rangle \vert v_y > 0$ [nm/sec] | 190±24 | 173 ± 23 | 284 ± 23 | 92 ± 13 |
| $\langle v_y \rangle \vert v_y < 0$ [nm/sec] | -152±23 | -143 ± 16 | -299 ± 75 | -277 ± 104 |
| $\langle \tau_p \rangle$ [sec] | 2.1±0.4 | 3.0 ± 0.8 | 4.1 ± 0.5 | 11.7 ± 1.4 |
| $\langle \lambda^* \rangle$ [nm] | 2200±300 | 2600 ± 540 | 4100 ± 820 | 4900 ± 750 |
| $\langle \lambda \rangle$ [nm] | 1700±260 | 2300 ± 480 | 3100 ± 660 | 4300 ± 680 |
| Fractions of forward (f) & backward (b) steps | 0.9 f \| 0.1 b | 0.98 f \| 0.02 b | 0.92 f \| 0.08 b | 0.92 f \| 0.08 b |
| Fraction of Right (r)- & Left (l)-handed steps | 0.45 r \| 0.55 l | 0.45 r \| 0.55 l | 0.94 r \| 0.06 l | 0.93 r \| 0.07 l |

Notably, about 90 *%* of the steps are minus-end directed (*ν*_x_ > 0); see Table 2, Table S3.2 and Fig. S3.1 for the distributions of the *temporal absolute* velocity, ν, longitudinal velocity, *ν*_x_, and transverse velocity, *ν*_y_. To further characterize the NP motions – as also shown in Fig. 4 and Table 2 – we have extracted the mean values of the longitudinal velocity (minus-end directed, *ν*_x_ > 0, and plus-end directed, *ν*_x_ < 0, motions) and of the transverse velocity (left, *ν*_y_ < 0, and right, *ν*_y_ > 0). As [NLS] and [CE] increase – implying a decrease of ξ and increase of the number of NP bound motors, 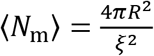 – the corresponding absolute and longitudinal velocities decrease. Conversely, the probability of plus-end directed and minus-end directed motions do not vary greatly (Table 2). Notably, in the diluted systems (i.e. system I, for the pair of systems I-II, and III for pair III-IV), the longitudinal minus-end directed velocity, 〈*ν*_x_〉 | *ν*_x_ > 0, is significantly higher – in absolute values – than the plus-end directed velocity 〈*ν*_x_〉 | *ν*_x_ < 0, suggesting that NPs move mostly in the direction of the MT minus-end. Moreover, in these NLS diluted systems, the transverse motion is significantly smaller than the longitudinal velocity.

#### Angular / Transverse motion

As discussed above, while most dynein motor studies assume purely longitudinal motion, more recent studies discovered rich transverse dynamics^36,37,47,61,55^. Accordingly, we analyzed the NP trajectories for transverse motion. First, we calculated (see Materials and methods and SI Sec. 4) the estimated transverse motion, *ν*_y_, of the NPs for the different systems (Fig. 4D). To obtain further insight, as shown in Fig. 5 and Table S5.1 (see also Materials and methods), we map the transverse velocity into an angular velocity, 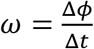 (where Δ*ϕ* is the angular increment of a single time step, Δ*t*), assuming an ideal angular motion with constant orbital radius, *R*_orb_ = 153 nm (Fig. 2C); the latter is estimated by considering molecular dimensions (Materials and methods and SI Sec. 4). The angular motion, together with the longitudinal motion, implies that if the MT had been elevated above the surface, the NP would have performed a helical motion around the MT symmetry axis. Ignoring the fluctuations in angular motion, we could consider the mean angular velocity associated with Fig. 5A as a representative number and estimate the mean pitch size of the *assumed* helix using 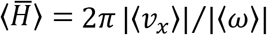 (see Materials and methods and SI Sec.4). This leads to the results shown in Fig. 5C and Table S5.1.

**Figure 5.**
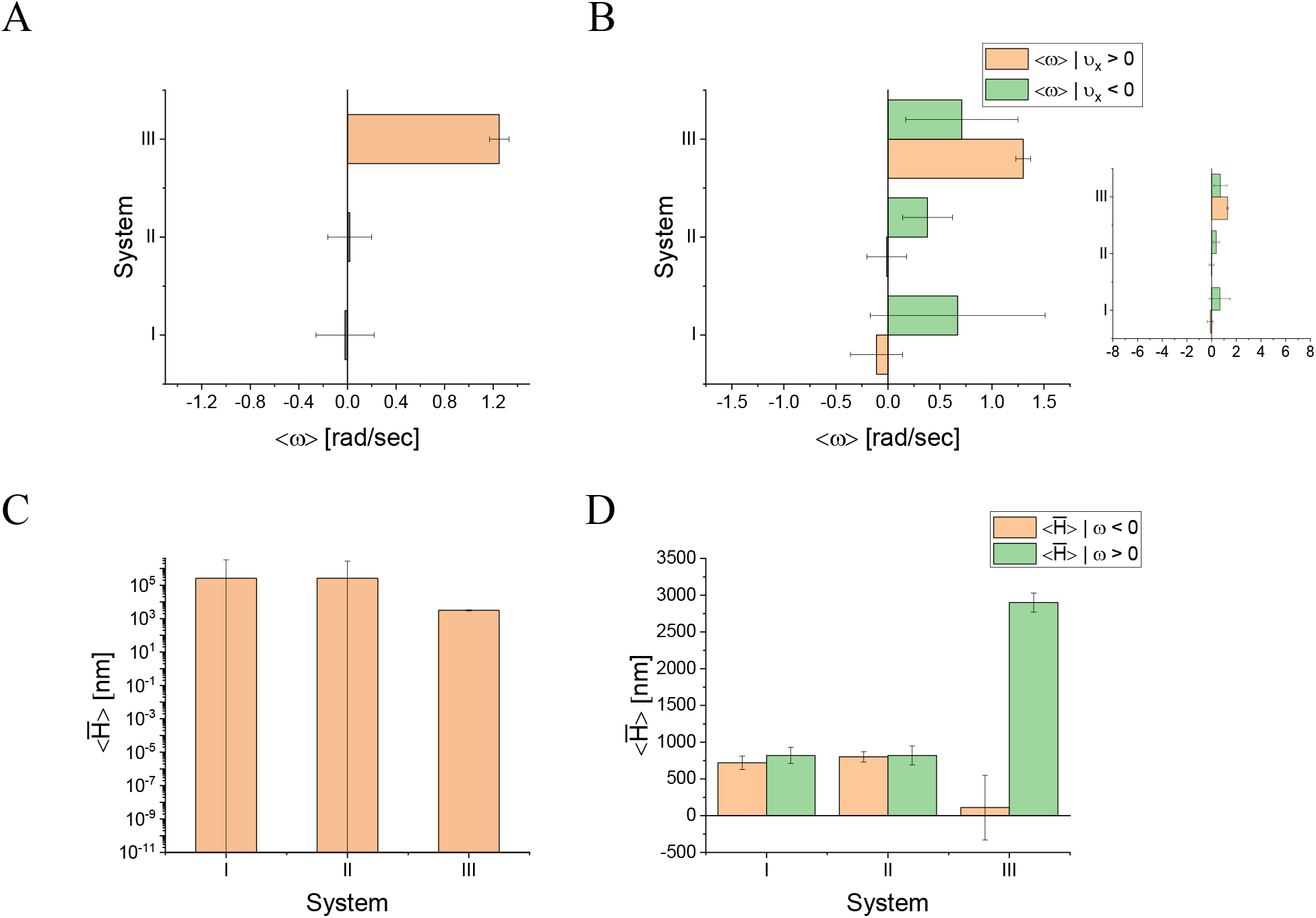
Mean angular velocity, 〈*ω*〉, and mean helical pitch size, 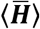, for systems I, II, and III. **(A)** Mean angular velocity, 〈*ω*〉. **(B)** Mean angular velocity evaluated separately for minus-end directed (*ν*_x_ > 0) and plus-end directed (*ν*_x_ < 0) longitudinal motions. Inset: same as main figure, but on a larger x-axis scale, similar to the scale of Fig. 9C (theory). **(C)** Mean helical pitch size, 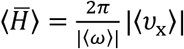, and **(D)** mean helical pitch size, 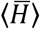, evaluated separately for right-handed (*ω* > 0) and left-handed (*ω* < 0) angular motions. The bare NPs mean diameter is 40 nm. For further details see SI Sec. 4 and Table S5.1.

However, using the theoretical model simulations presented in the next section, we demonstrate in Movies 3-6 and Fig. 6 selected trajectories of a single NP, showing angular motion “fluctuations” of different magnitudes, such that the NP effectively performs both right- and left-handed helical motion in a single trajectory. We define a helical motion as a single and complete turn (*ϕ* = ±2*π*) around the MT. Therefore, we now split the angular motion into right- and left-handed motion (Table S5.1; see Materials and methods for details). For systems I and II, the resulting mean angular velocities shows that indeed the right- (〈*ω*〉|*ω* > 0) and left- (〈*ω*〉|*ω* < 0) handed motions occur at similar angular velocities, which almost cancel each other when we consider the overall mean value, 〈*ω*〉, as suspected. The resulting mean helical pitches for the anticipated right- 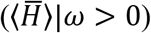 and left- 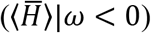 handed helices (in a hypothetical surface free experiment) are comparable to each other (Fig. 5D and Table S5.1; see also Materials and methods).

**Figure 6.**
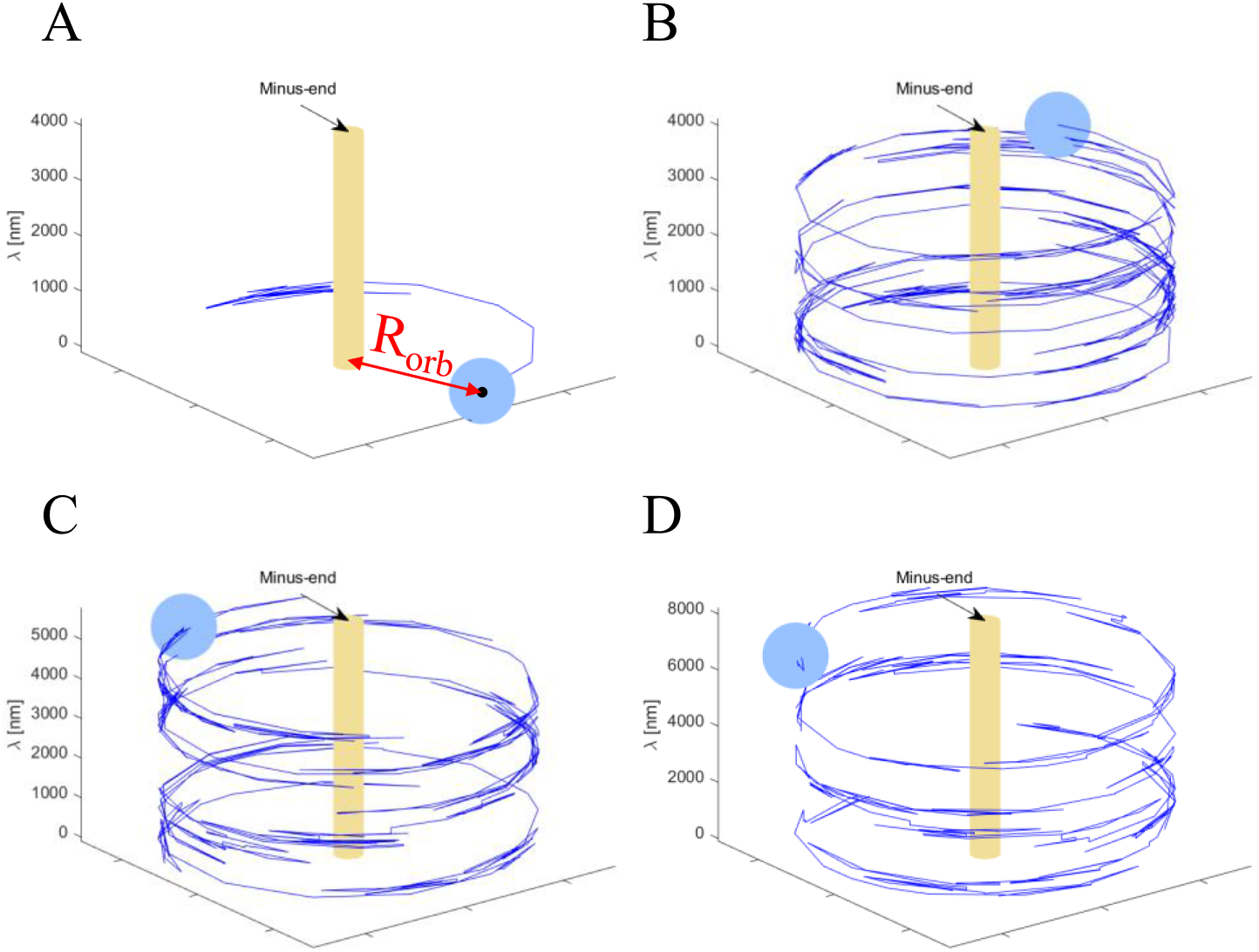
Traces of selected simulated trajectories. The light-blue sphere represents the NP (*R* = 20 nm), the blue trailing lines represent the trajectory, and the yellow cylinder in the middle represents the MT. Note that while the MT and NP radii are drawn to scale, the MT length is not. **(A)** Initial trace of a single motor trajectory. The red arrow marks the distance between the MT and NP centers, *R*_orb_. **(B-D)** Traces of selected trajectories of a NP having *N*_m_ = 1,3, and 13 motors, respectively.

Furthermore, we analyze the correlations between longitudinal direction *and* angular directions. In Fig. 5B (see also Table S5.1), we present the mean angular velocity for separated minus-end directed (〈*ω*〉|*ν*_x_ > 0) and plus-end directed (〈*ω*〉|*ν*_x_ < 0) motions for systems I-III (the inset shows the same results on a large *x*-axis scale). We observe that, for *all systems, ν*_x_ < 0 (green-colored columns) is associated with *ω* > 0, and – except for system III – *ν*_x_ > 0 (orange-colored columns) is associated with *ω* < 0. Thus, we demonstrate that backward (plus-end directed) steps – although being relatively rare – are highly correlated with right-handed motion. Comparing systems, I and II, we see that in system I forward steps are highly correlated with left-handed motion, while in system II the azimuthal motion associated with *ν*_x_ > 0 essentially vanishes. This suggests that the increase of NP bound motors (larger in II vs I) is suppressing the latter correlations (i.e. *ν*_x_ > 0 correlated with *ω* < 0, and they are likely a single motor property, as supported by our theoretical results described below. The implications of plus-end directed steps being correlated with right-handed azimuthal steps will be addressed in the Discussion section.

### 1.4 Theoretical predictions

Results were obtained for various NP configurations, which are characterized by two parameters: *R* and *N*_m_. Below we discuss NP of radius *R* = 20 nm and number of motors ranging between *N*_m_ = 1 to *N*_m_ = 13. For each (*R* = 20 nm, *N*_m_), we ran the simulations for a few hundreds of (identical) particles to obtain high statistical accuracy. Note that, due to the very small NP drag coefficient (SI Sec.7), *N*_m_ = 1 is used as a representative of the single motor case. The motility characteristics (on a time scale of 0.27 sec) of each NP configuration are described in Figs. 7, 9 and in SI Sec. 8. In Figs. 8–10 and SI Sec.9, we complement these with analyses on the timescale of a single MC step (which is 1-2 orders of magnitude smaller). Selected trajectories of single-motor, three-motor, and seven-motor NPs are shown in Fig. 6 and Movies 3-5, demonstrating rich, fluctuating, helical motions. Below we investigate the influence of the number of motors on such behavior.

**Figure 7.**
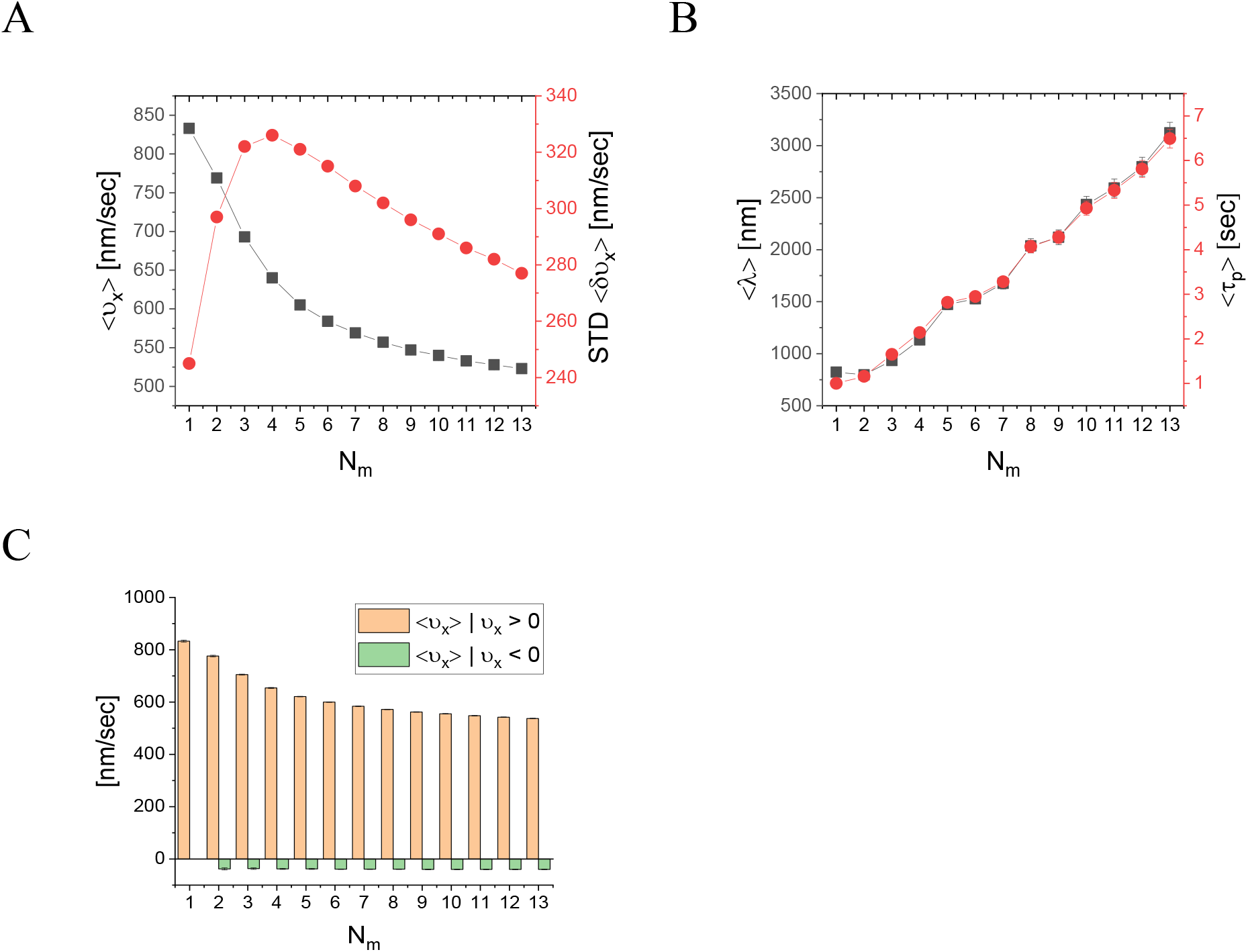
Simulation results for the mean longitudinal velocity, 〈*ν*_x_〉, run-length, 〈*λ*〉, and run-time, 〈*τ*_p_〉 for different 〈*R* = 20 nm, *N*_m_) configurations, for time interval Δ*t* = 0. 27 sec. **(A)** Mean longitudinal velocity (mean ± SEM) and STD (raw data is provided in Table S8.1; see Fig. S9.1 and Table S9.1 for Δ*t* = MC step time). **(B)** Mean run-length and run-time (mean ± SEM) (raw data is provided in Table S9.2). **(C)** Mean instantaneous longitudinal velocity, 〈*ν*_x_〉, separated for minus-end directed, *ν*_x_ > 0, and plus-end directed, *ν*_x_ < 0, directions (mean ± SEM) (raw data is provided in Table S8.2).

**Figure 8.**
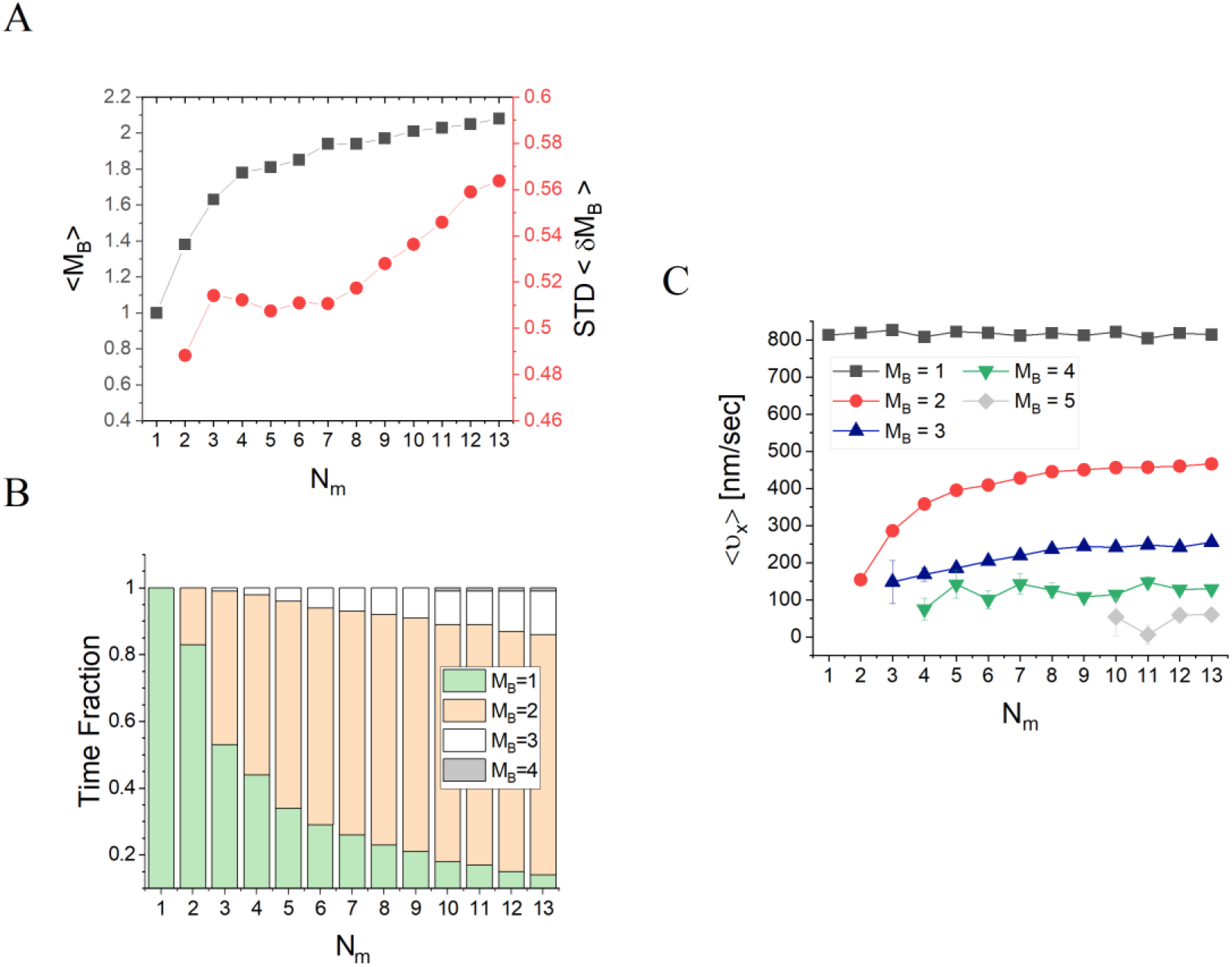
Participating (MT-bound) motors, *M*_B_, for different (*R* = 20 nm, *N*_m_) configurations, for time interval Δ*t* = MC-step time. **(A)** Mean number of participating motors 〈M_B_〉 and STD against the number NP-bound motors N_m_ (mean ± SEM) (raw data appears in Table 3). **(B)** Fraction of time the NPs spend in each of the MB-states, for different values of *N*_m_ (raw data appears in Table 4). (**C**) Mean longitudinal velocities of the different MB-states for different values of *N*_m_ (mean ± SEM) (raw data appears in Table 5). Note that states of *M*_B_ ≥ 5 are extremely rare for *N*_m_ ≤ 13; therefore in (B) the state *M*_B_ = 5 is not shown at all and is appears only for *N*_m_ ≥ 10 in (C).

**Figure 9.**
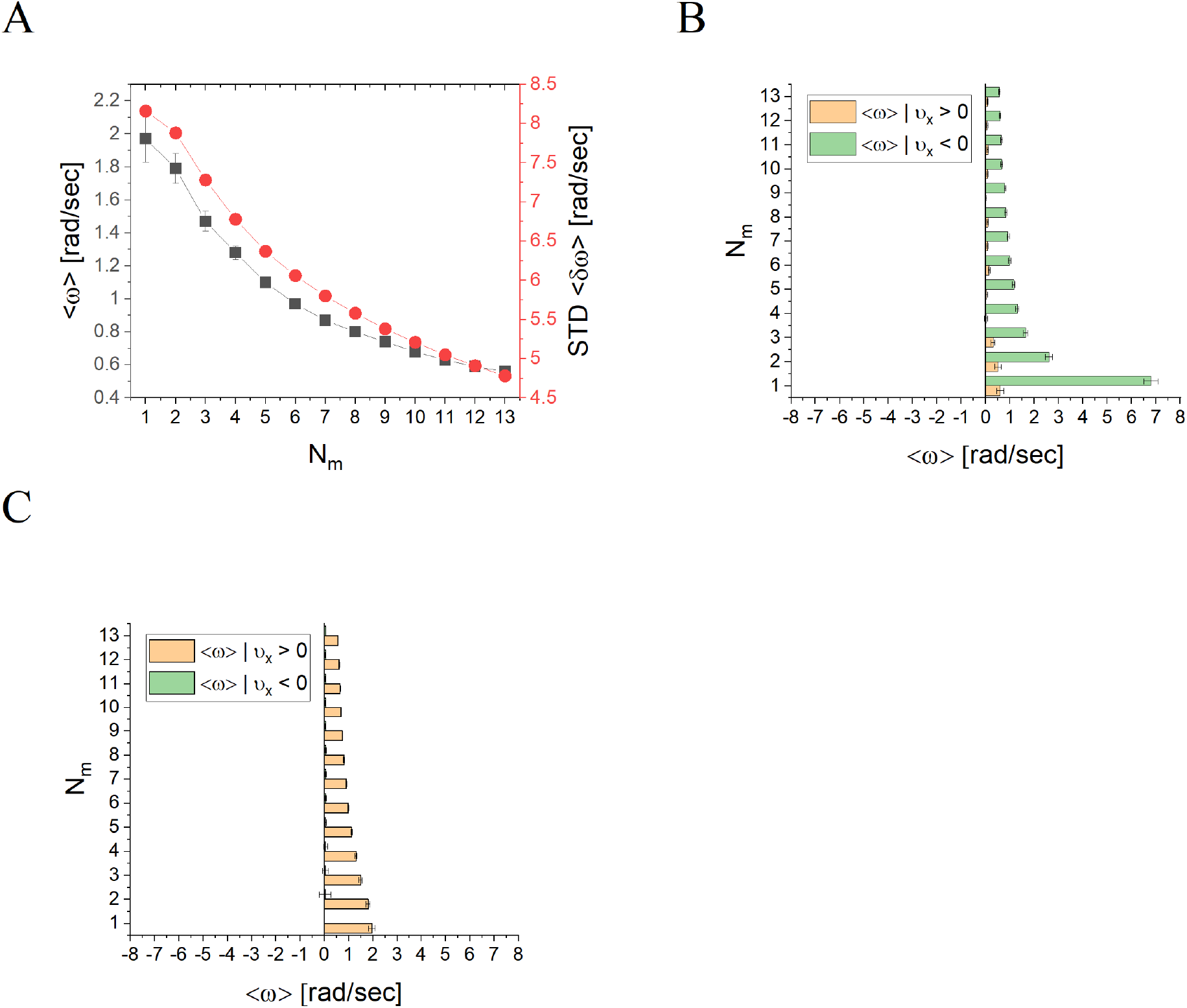
Mean angular velocity, 〈*ω*〉, as a function NP-bound motors *N*_m_, for NPs of radius *R* = 20 nm. **(A)** Mean angular velocity, 〈*ω*〉, and the corresponding STD, against *N*_m_, for time interval Δ*t* = 0.27 sec (raw data provided in Table S8.3; see Table S9.6 and Fig. S9.4 for Δ*t* = MC step time). **(B and C)** Mean angular velocity calculated separately for minus-end directed motion, 〈*ω*〉 | *ν*_x_ > 0, (orange bars) and plus-end directed motion, 〈*ω*〉 | *ν*_x_ < 0, (green bars), against the number of NP-bound motors *N*_m_, for the time intervals Δ*t* equal to (B) MC-step time (raw data provided in Table S9.8) and (C) 0.27 sec (raw data provided in Table S8.4). **(B and C)** Values correspond to mean ± SEM.

**Figure 10.**
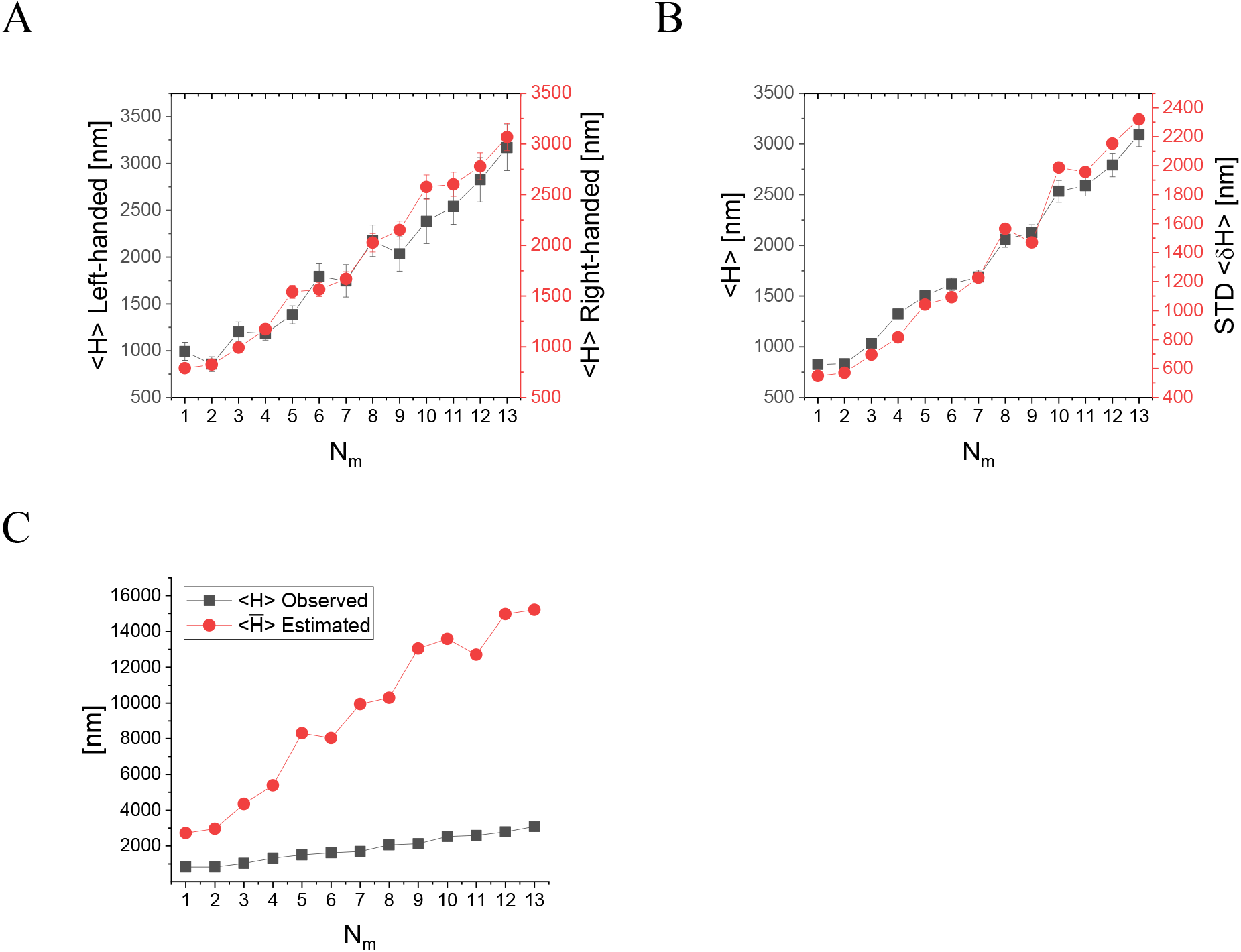
Analysis of the actual NP helical motion from observed trajectories. **(A)** Mean helical pitch size, 〈*H*〉, evaluated separately for left- and right-handed helical vorticity (raw data provided in raw data is provided in Table S9.9). **(B)** Mean and STD of the helical pitch size, regardless of the helical vorticity (raw data provided in Table S9.10). **(C)** Comparison between the actual helical pitch size (from the observed simulated trajectories) and the helical pitch size estimated using the mean values of angular and longitudinal velocities, 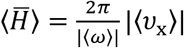 (raw data provided in Table S9.11).

#### Longitudinal velocity

Fig. 7A shows the mean longitudinal velocity, 〈*ν*_x_〉, against *N*_m_. We find suppression of 〈*ν*_x_〉 from the single motor velocity (833 ± 4 nm/s) as *N*_m_ increases, which is a clear signature of motor-motor coupling in the absence of any load-sharing effect (due to the vanishing drag). The corresponding standard deviation about the mean also varies with *N*_m_; a maximum appears at *N*_m_ = 4 (corresponding to *ξ* = 35.4 nm), whereas the minimal standard deviation corresponds to the single motor case. In addition, as seen in Fig. 7B, the characteristic run-time, 〈*τ*_p_〉, and longitudinal run-length, 〈*λ*〉, are also affected by the value of *N*_m_. We observe an effectively monotonous increase of 〈*τ*_p_〉 and 〈*λ*〉 with an increase of *N*_m_ up to *N*_m_ = 13. Following the experimental analysis shown in Fig. 4C, we compute separately the mean longitudinal velocity of minus-end directed steps and plus-end directed steps, see Fig. 7C. We observe a monotonous decrease of the mean minus-end directed velocity with increasing *N*_m_, effectively saturating above *N*_m_ ≈ 7, similar to the trend seen in the experimental results; the velocity of the plus-end directed steps is not sensitive to *N*_m_.

To better understand the connection between these results and the actual number of motors participating in the motion, we depict in Fig. 8A the mean number (per MC unit time) – over all runs – of transient MT-bound motors, 〈*M*_B_〉, for each *N*_m_; see also Table 3. As expected, as *N*_m_ increases the value of 〈*M*_B_〉 increases, yet, surprisingly, it effectively saturates at around 〈*M*_B_〉 = 2. Furthermore, we extracted the fractions of time for different NP “states”; we define an NP state by the number of MT-bound (transporting) motors *M*_B_ (Fig. 8B); see also Table 4. Note that although 〈*M*_B_〉 is roughly ranging between one and two, the motor number distributions are wide and show that there is significant contribution of the one-, two-, and three-motor states (while *M*_B_--states of four motors and above have negligible contribution). Fig. 8C shows the corresponding mean velocity for each state (see also Table 5), avoiding NP temporal velocities associated with transitions between these states *via* binding-unbinding events. Surprisingly, the mean velocity of a particular *M*_B_-state increases with elevation of *N*_m_, rather than remaining a constant. This can be rationalized by noting that an MB-state still corresponds to several microscopic configurational states. At larger *N*_m_, i.e. smaller NP anchoring distances, most of such microscopic states are associated with MT-bound motors whose NP anchoring distance *ξ* is smaller. This implies to a reduction in polymer tension, i.e. suppression of the elastic coupling between MT-bound motors. Indeed, such an effect has been observed in previous theoretical studies^31,32,35,41^, and is associated with the fact that a forward pulling force on a lagging motor, while somewhat increasing its temporal velocity, has a much smaller effect than the reduction in temporal velocity due to a backward pulling force on a leading motor. This analysis confirms that the reduction with increasing *N*_m_ of the longitudinal velocity – both experimentally and in the simulations – results from the increased number of motors participating in motion, MB (i.e. MT-bound motors).

**Table 3.** Raw data for Fig. 8A. Mean number and STD of transient MT-bound motors, *M*_B_, for different (*R* = 20 nm, *N*_m_) configurations, for time intervals equal to MC step times.

| $N_m$ | 1 | 2 | 3 | 4 | 5 | 6 | 7 |
| --- | --- | --- | --- | --- | --- | --- | --- |
| $\langle M_B \rangle$ | 1 | 1.38 | 1.63 | 1.73 | 1.81 | 1.85 | 1.94 |
| $\langle \delta M_B \rangle$ | 0 | 0.4883 | 0.5142 | 0.5123 | 0.5075 | 0.5110 | 0.5107 |
| $N_m$ | 8 | 9 | 10 | 11 | 12 | 13 | |
| $\langle M_B \rangle$ | 1.94 | 1.97 | 2.01 | 2.04 | 2.07 | 2.09 | |
| $\langle \delta M_B \rangle$ | 0.5174 | 0.5280 | 0.5363 | 0.5459 | 0.5590 | 0.5638 | |

**Table 4.**
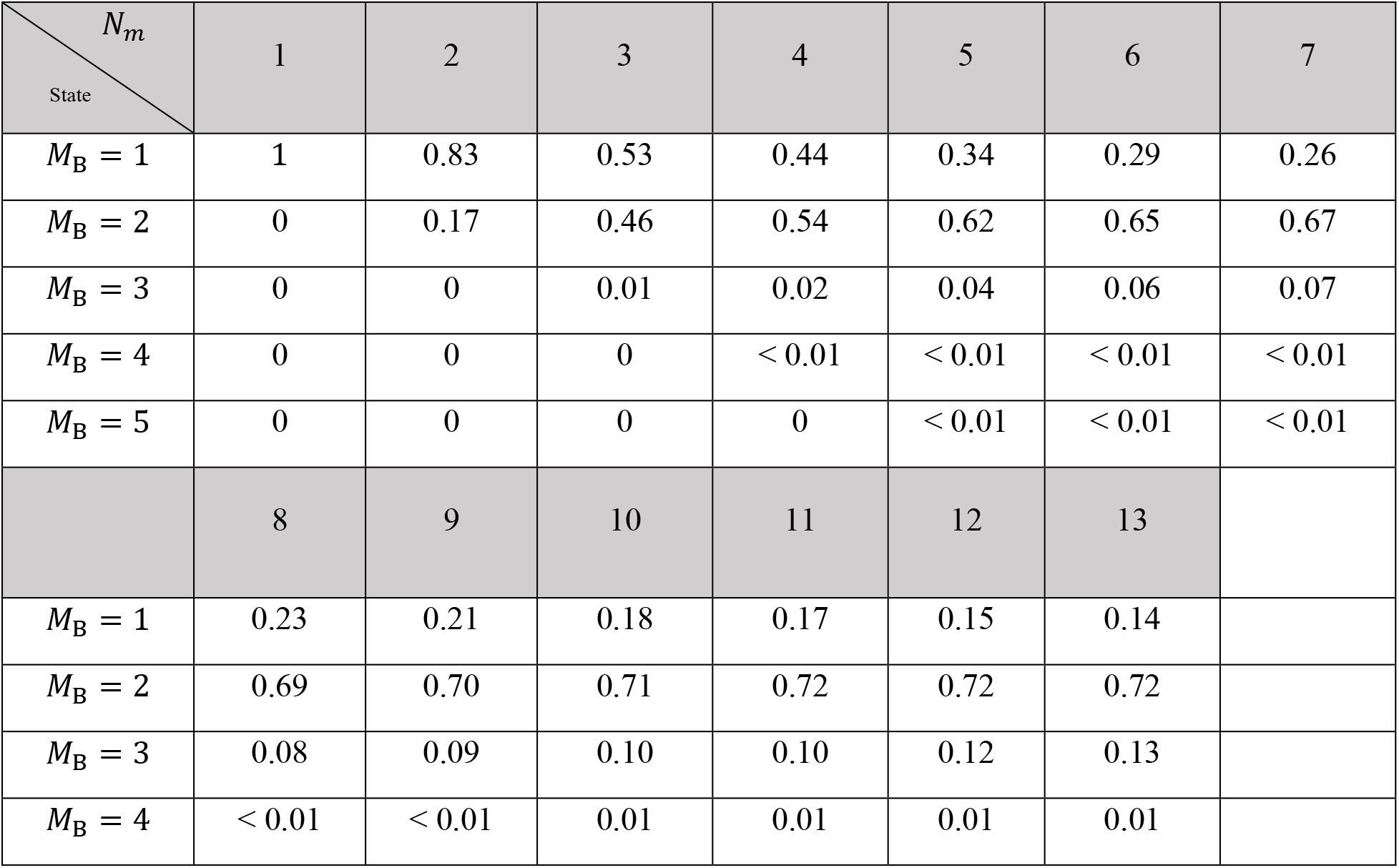
Raw data for Fig. 8B. Time fraction of the different NP states, where a state is defined according to the number of MT bound motors, *M*_B_, for different (*R* = 20 nm, *N*_m_) configurations, and for time intervals equal to MC step times.

**Table 5.**
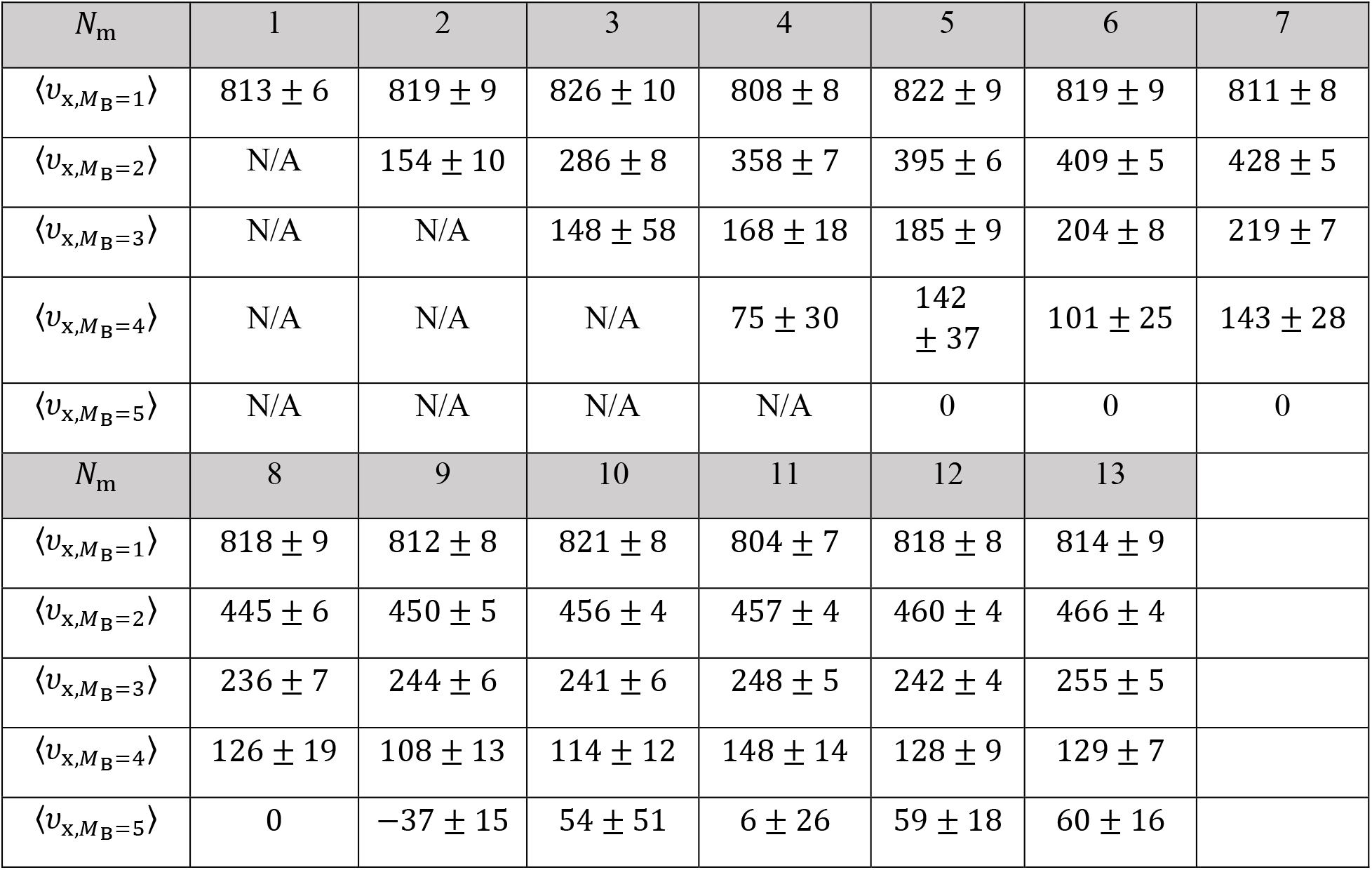
Raw data for Fig. 8C. Mean longitudinal velocity of the different NP states, where a state is defined according to the number of MT bound motors, *M*_B_, for different (*R* = 20 nm, *N*_m_) configurations, for time intervals equal to MC step times (mean ± SEM).

As manifested by the STD shown in Figs. 7A and S9.1 (and by the corresponding histograms in Figs. S8.1 and S9.2, respectively), the width of the NP longitudinal velocity distribution varies with *N*_m_. We associate this width *both* with the width of the velocity distribution at each MB-state (not shown here), and with the probability (i.e. time fractions) distribution of these states (Fig. 8B), which is sensitive to the value of *N*_m_. For *N*_m_ ≥ 2, we unexpectedly observed a second peak around *ν*_x_ = 0, which becomes more pronounced as *N*_m_ increases (Fig. S8.1). This is more strongly demonstrated by the equivalent histograms corresponding to a single MC time-step (Fig. S9.2). We associate this new peak to the growing abundance – with increasing *N*_m_ – of states *M*_B_ ≥ 2 (Fig. 8C). These states have a greater chance for “jamming configurations,” i.e., configurations where the “jamming events” (controlled by motor-motor excluded-volume interactions, see Methods section) are dominating.

From this analysis, we can conclude that the increase of *N*_m_ leads to a non-linear increase of 〈*M*_B_〉. This increase of 〈*M*_B_〉 leads, in turn, to longer run-times, 〈*τ*_p_〉, and diminishing 〈*ν*_x_〉, yet the effect on the run-time is more pronounced than on 〈*ν*_x_〉 such that, in most cases, the resulting run-length, 〈*λ*〉, is also enhanced. Moreover, the increase of 〈*M*_B_〉 leads to a slightly narrower NP longitudinal velocity distribution for *N*_m_ > 4. While an increase of run-time is expected even without the inclusion of motor-motor coupling^16^, the decrease of velocity is a sole consequence of the (elastic and excluded volume) motor-motor interactions. Similarly, the *non-trivial* change of width of the ν_x_ distribution with increasing *N*_m_ (STD in Fig. 7A) reflects the competition between: (i) the increase of the MB fluctuations (i.e., the corresponding STD, 〈*δM*_B_〉; Table 3), acting to increase the velocity fluctuations, (ii) the variability of the velocity fluctuations within each *M*_B_-state (i.e. (*δv*_x,M_B__); Table S9.4), which can be attributed to motor-motor coupling, and (iii) the contributions of the unbinding events, since after each unbinding event an immediate jump of the NP position occurs to balance elastic forces; unbinding events increase with increasing *N*_m_, see Fig. S9.3 and Table S9.5.

#### Angular / Transverse motion

Consider now the angular motion around the MT symmetry axis that leads to a transverse motion on the projected base plane. As can be seen in Fig. 6 and Movies 3-6, the motion is composed of apparent both left- and right-handed helices combined with large fluctuations. However, on a very long trajectory, the net helical motion (in which the right- and left-handed helices cancel each other) might appear minor, and will not reflect the true nature of the motion. Therefore, we require here very delicate analyses that will reflect both characteristics.

The variation of the mean angular velocity 〈*ω*〉 and its STD with *N*_m_ is shown in Fig. 9A (and Fig S9.4), both demonstrating a monotonous decrease with an increase of *N*_m_. As all values are positive, this suggests a net right-handed helical motion of the NP for all motor numbers. Combined with a variation of the mean longitudinal velocity with *N*_m_ (Fig. 7A), and the relation 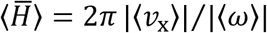 describing the mean helical pitch size 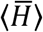, this leads to the dependence of 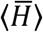 on *N*_m_ shown in Fig. 10C, exhibiting an increase of 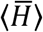 for growing *N*_m_.

However, as discussed in the experimental section, this pitch size represents the *net* helical motion, i.e., it includes cancelations of left- and right-handed helices (as seen in Fig. 6 and Movies 3-6). To refine the helical motion analyses, we define a helix (be it left- or right-handed) whenever the angular motion completes a full round (i.e., *ϕ* = ±2*π*). Note that, even within a single full round, the motion consists of large and frequent right- and left-handed fluctuations.

In Fig. 10A, we show the resulting mean pitch size for left- (*ϕ* = −2*π*) and right- (*ϕ* = +2*π*) handed helices separately, and the mean pitch size regardless of the helix vorticity. Notably, the pitch sizes obtained for all these three definitions are all comparable to each other and are much shorter than those deduced using the mean angular velocity – all increase with increasing *N*_m_. To complement these results, we also present, in Fig S9.5, the mean angular velocity separately for left- and right-handed motion.

#### Longitudinal and angular/transverse motion are correlated

To gain further insight into the complex motion of the NP, we wish to verify whether the longitudinal and angular motions are correlated. In Fig. 9B, we *dissect* the angular velocity for forward and backward steps associated with MC time-step (i.e., positive and negative longitudinal velocities). As seen, backward steps have a greater (positive) mean angular velocity, with the single motor NP showing the largest value. This implies that when a backward step is being performed, the NP is likely to move to the right. Both angular velocity types, associated with either forward or backward steps, show a decrease with increasing motor number *N*_m_. For comparison, we also plot in Fig. 9C the same distributions on the experimental time interval (0.27 sec), showing that these correlations are suppressed. This signifies the importance of the value of the time interval for making the basis for comparison between different results, be it experimental or theoretical.

## Discussion

Comparison between experiment and theory requires consideration of two issues: (i) Knowledge of the relative NP motor coverage 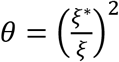. The latter corresponds to the mean number of NP-bound motors *via* 〈*N*_m_〉 = 〈*N*〉 × 0, where 〈*N*〉 is the mean number of PEG-NLS polymers per NP. (ii) The expected (wide) binomial distribution of *N*_m_ between the different NPs. Regarding the first issue, we can reasonably assume that for a given [NLS], the increase of [CE] implies increase of 0, which allows qualitative comparison between different systems – as discussed below. Regarding the second issue, in SI Sec. 10, we have computed – using the binomial distribution – some of the reported motility characteristics for a varying 〈*N*_m_〉. For the mean longitudinal and transverse velocities calculated on the MC time step scale, this analysis shows a small increase of their values (relative to the values for a deterministic motor number, *N*_m_), especially for small motor numbers (Figs. S10.1 and S10.2); this is due to the contribution from NPs with very few motors (*N*_m_ = 1,2,3), whose velocities are always larger (Fig. 7A). Yet, the trends we have deduced theoretically for the deterministic number of motors (Figs. 7 and 9), are not altered. When we move to the experimental time scale (0.27 sec), the difference is almost non-discernable (Figs. S10.1 and S10.2). Moreover, the run-times and run-lengths are even less affected by the *N*_m_ fluctuations, probably due to the contributions from NPs with large *N*_m_ for which these values are much higher. Hence, we do not emphasize here anymore the difference between (simulation) results for deterministic *N*_m_ and those corresponding to 〈*N*_m_〉 using the binomial distribution.

Comparison between experiment and theory shows several similarities regarding the dependence of both longitudinal and angular motion on the number of NP-bound motors. Note again that (in the experiment) we have estimated the number of PEG-NLS (*N*) for the different [NLS] used for the motility assays. First, as *N* increases, while keeping [CE] fixed (or in other words, *θ* fixed) – implying an increase of *N*_m_ (or 〈*N*_m_〉), we observe both theoretically and experimentally a decrease of the longitudinal mean velocity (Figs. 4A, 7A, and S9.1). As for the angular velocity, *ω*, while the theoretical values show a decrease of *ω* with increasing *N*_m_, the estimated experimental values do not follow a clear trend. Regarding the actual values of *ω* and *ν*_x_, we can discern that both their values in the experimental system III match the theoretical values for *N*_m_ = 4 (or, 〈*N*_m_〉 = 4) (Tables 2, S5.1, S8.1, and S8.3). This could have been accidental. However, we find further evidence suggesting that the match is not accidental. Since *N* in system IV is 3.5 times larger than in system III (Table 1) while they both have the same [CE] and they are both prepared using the same extract, it follows that in system IV, 〈*N*_m_〉 should be 3.5 times larger as well, i.e., 〈*N*_m_〉 ≅ 14. Comparison of *ω* and *ν*_x_, between system IV (experimental: *ω* = 0.25 rad/sec and *ν*_x_ = 422 nm/sec) and extrapolated theoretical values (linear fit) for *N*_m_ = 14 (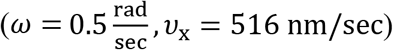 indicates a rough agreement between theory and experiment. As for these four systems run-times and run-lengths, both simulation and experimental values show an increase of their values with increasing *N*_m_ (due to increase of 〈*N*〉).

Further support of the above match appears in the width of the distributions of *ν*_x_ as exemplified by the corresponding STD. In the simulations, we observed first an increase of STD, followed by a decrease as *N*_m_ increases, with a maximum STD at *N*_m_ = 4 (Fig. 7A and Table S8.1). In the experimental results, there is an apparent maximum in the STD corresponding to system III (Table S3.2). Importantly, these two systems are just those found above to match in longitudinal and angular velocities.

Considering systems I and II, the most prominent effect seen in experimental studies is the strong correlation between backward (plus-end directed) and right-handed motions (Fig. 5B). Such strong correlations also appear in the theoretical studies when the analysis is performed on the MC time scale (Fig. 9B). Yet, these correlations are smeared when the analysis is performed on the experimental time interval (Fig. 9C). We are unable to pinpoint to the origin of this discrepancy. Despite this apparent discrepancy, both theoretical (MC time interval) and experimental studies show diminishing of these correlations with increasing *N_m_*. Thus, it is clear that these correlations emerge from the single motor property, and are somewhat suppressed by the action of multiple motors.

## Conclusions

In this paper, we reported on a combined experimental and theoretical study of NPs that are carried by a multiple number of *mammalian* dynein motor-proteins. We focus on the different motility characteristics of these multi-motor NPs in comparison with the known and well-studied case of the single dynein^6,17–20,37,38,61–65,55^. The multi-motor NPs differ from single motor behavior in several aspects. First, as the number of NP-bound motors (*N*_m_) increases, we observe decrease of the mean longitudinal velocity (*ν*_x_). Second, we observe increase of the run-times (*τ*_p_) and run-lengths (*λ*). Third, the width of the longitudinal velocity distribution depends peculiarly on the number of NP-bound motors, showing a maximum width at an intermediate value.

We also found that both single and multi-motor NPs perform an angular (transverse) motion that, together with the longitudinal motion, forms a helical trajectory around the MT symmetry axis. This is consistent with previous findings of Yildiz and co-workers^36,37,55^ in which large particles were studied (about 500 nm radius) and in which the number of particle-bound motors was not varied. Importantly, here we have demonstrated that the helical motion stems from single motor properties (consistent with previous claims^51^) and is somewhat suppressed (an increase of the helical pitch) as *N*_m_ increases.

One of the most peculiar features of the angular motion is its correlation to the direction of the longitudinal motion (i.e., minus-end or plus-end motion along the MT long-axis). We have unambiguously shown, both experimentally and theoretically, that plus-end directed motional intervals dictate large right-handed motion. Moreover, this characteristic also appears as a single motor property and it diminishes as more motors are available, as verified theoretically. In fact, our single *mammalian* dynein stepping model, which builds on the raw data of Yildiz and co-workers^37,55^, shows that these correlations do strongly appear in the single motor stepping. (Moreover, revisiting the results of our single *yeast* dynein stepping model, we found similar correlations^95^).

The good agreement between theory and experiment give mutual support to both. This has allowed us to use the simulations to unravel NP motional microscopic configurations that cannot be accessed in the experiment. In particular, we have deduced from the simulation the partitioning between the different number of motors, *M*_B_, that participate in the motion. These results have shown that most of the trajectories involve mainly 1-3 motors, with the mean value approaching two for large *N*_m_. In this way, the NP manages to maintain considerable longitudinal velocity, resulting in a longer run-length.

Our theoretical results for the NP motility features – supported by the experimental ones – suggest that the NPs are self-regulating nano-machines whose behavior is different from the single motor behavior. The self-regulation of the number of participating motors – regardless of the number of NP-bound motors – at a mean *M*_B_ value between 1 and 2 with dominating states *M*_B_ = 1, 2, 3, suggests that the NP is able to optimize between single motor and multi-motor motility properties. This allows the NP to achieve the following features: (i) longer run-time and run-length, (ii) substantial helical motion, (iii) significant longitudinal velocity, and (iv) plus-end directed motional intervals strongly correlated with large right-handed motion, especially during the temporal state of a single participating (MT-bound) motor. The latter plausibly implies on a mechanism for obstacle bypassing^51^. This behavior is strongly enhanced by temporally alternating to the single motor state (*M*_B_ = 1).

Some native cargos are rigid, e.g., the (cores of) HIV^13^, Herpes-Simplex virus^66^, and adenoviruses^14,15^, and super-coiled plasmids^67^, while others – such as cytoplasmic lipid granules (i.e. native liposomes) – are relatively flexible. Our current model simulations – as well as previous theoretical results^32,41,67^ – and the present motility assays of a rigid NP, demonstrate that, for rigid cargos, maximal transport efficiency can be achieved when flexible linkers mediate between the rigid NP body and the motor proteins, allowing the NP to optimally use the viable motors. We thus conjecture that rigid native cargos use a similar mechanism for their active intracellular transport, e.g., possibly the abundance of disordered loops in the NFκB structure gives it extra flexibility for enhancement of the super-coiled plasmid motility^67^. If used intracellularly, the ability of the NP to make long trajectories enhances its active transport towards the nucleus, which is relevant to both the rigid core viruses mentioned above and possibly to drug delivery applications^16,68,69^. In addition, by maintaining its helical motion, and in particular making large right-handed steps when stepping backward, we posit that the NP is capable of bypassing obstacles that reside on the MT tracks, or in their vicinity. Indeed, this suggestion – yet, without the mention of the backward-right-handed correlated motion – has already been put forward^36^, and further observed in (surface-free) bead motility assays^51^. We plan to further investigate these hypotheses in future work.

## Materials and Methods

### Experimental procedure

#### Materials

##### Hela cell extracts preparation

Hela cell extracts were prepared according to Fu et al.^70^. Briefly, Hela cells were grown in DMEM medium supplemented with 10 % of Fetal Bovine Serum (FBS) and 1 % of penicillin-streptomycin at 37 °C and 5 % CO_2_. The cells were detached by Trypsin-EDTA solution, washed with Phosphate Buffered Saline (PBS) (0.01 Phosphate buffer, 0.0027M KCL, 0.137 M NaCl, pH7.4), and pelleted for 5 min at RT and 500 g. The cells were then incubated on ice with lysis buffer (12 mM Pipes, 2 mM MgCl2, 1 mM EGTA, 0.4 % Triton X-100, 10 % protease inhibitor cocktail, pH 6.8) for 15 min. Finally, the lysates were cleared by centrifugation at 2,700 g and then at 100,000 g, at 4 °C for 10 min each. The cell extracts are diluted 10x with lysis buffer without Triton and protease inhibitors, and the total protein mass concentration is determined by Bradford assay. Sucrose was added to the extracts (10 % in mass), which were then flash-frozen in liquid nitrogen and stored at −80 °C. For the motility experiments, the extracts were used within four weeks.

##### Nuclear Localisation Signal (NLS)

The NLS sequence (PKKKRKVED) originates from SV40 T large antigen^71^. We use N-terminally bromine (Br-Ac) modified NLS. The bromine group is followed by a GGGG sequence (‘raft’). To generate a fluorescently labeled NLS peptide, Tetramethylrhodamine (TAMRA) was covalently attached to the NLS raft. Except for UVVis absorption experiments in which we use TAMRA-NLS, all experiments were performed using the non-labeled NLS.

##### NPs preparation

Green Fluorescent microspheres (Bangs Labs or Invitrogen) were used for NP preparation. The total surface area of the bare NPs was maintained constant through all experiments. NPs preparation starts with their incubation in 10 μM Neutravidin (31000, Thermo Fisher) for 20 min at 25 °C. Excess Neutravidin was separated via dialysis. (Note, that as a starting point, one can use Streptavidin-coated NPs (Bang Labs) instead of bare microspheres). Next, the NPs are incubated for 20 min at 25 °C with 10 mg/mL BSA to block remaining uncoated regions on the NPs surface. Excess BSA is separated by centrifugation (6800 g for 8 min at 25 °C). All subsequent separation steps are performed at the same centrifugation conditions. The NPs were then incubated with 1 mM Biotin-PEG-thiol-5kDa (PG2-BNTH-5, NANOCS) for 30 min at 25 °C. Excess Biotin-PEG-thiol was separated by two cycles of centrifugation. Then, NLS peptides were covalently bound to the NPs via their bromine group, which react with the thiol group on the PEG molecule. The reaction was performed at 25 °C for 20 min. Excess NLS was separated by two cycles of centrifugation. Finally, the NPs were incubated with Hela cell extracts at 30 °C for 20 min, allowing the recruitment of α- and β-importins, and *mammalian* dynein motors to the surface of the NPs. The NPs were separated from excess cell extracts by centrifugation, washed with BRB80 (80 mM PIPES, 1 mM MgCl2, 1 mM EGTA, pH 6.8) supplemented with 1 mM Mg-ATP, and centrifuged again. Finally, the NPs were suspended in 60 μL BRB80 supplemented with *1* mM Mg-ATP and stored on ice until used. In the motility assay experiments, we used two different batches of cell extracts. An identical cell extract was used in systems I and II, and another extract in systems III, IV, and V. To assure full activity of the motility protein machinery, the NPs were used within 3 h.

#### Methods

##### Cryo-electron microscopy (Cryo-TEM)

Samples for cryo-TEM were prepared according to a standard procedure^72^. Vitrified specimens of the NPs solution were prepared on a copper grid coated with a perforated lacy carbon 300 mesh (Ted Pella Inc.). *2.5* μL drop of that solution was applied on the grid and blotted with a filter paper to form a thin liquid film, few tens of nanometers thick. The blotted samples were immediately plunged into liquid ethane at its freezing point (−183 °C) using an automatic plunge freezer (Leica, EM GP). The vitrified specimens were then transferred into liquid nitrogen for storage. Samples were analyzed using an FEI Tecnai 12 G2 TEM, at 120 *kV* with a Gatan cryo-holder maintained at −180°*C*. Images were recorded on a slow scan cooled charge-coupled device CCD camera (Gatan manufacturer, Pleasanton, CA, USA) at low dose conditions, to minimize electron beam radiation damage. Recording is performed using Digital Micrograph software (Gatan).

##### Dynamics light scattering and ζ-potential experiments

DLS and *ζ*-potential measurements were used to conclude on the NP hydrodynamic diameter (*D*_h_) and charge at the various decoration steps. For these experiments, we use bare NPs of 0.196 μm in diameter. DLS and *ζ*-potential measurements were performed on a Malvern NanoZS instrument (ZN-NanoSizer, Malvern, England) operating with a 2 mW HeNe laser at a wavelength of 632.8 nm. The detection angle is 173 ° and 17 ° for DLS and ζ-potential measurements, respectively. All measurements were done in a temperature-controlled chamber at 25 °C (± 0.05 °C); for the analysis of *D*_h_ and ζ-potential, the viscosity is taken to be the same as that of water (0.8872 cP). The intensity size (hydrodynamic diameter) distribution was extracted from the intensity auto-correlation function calculated using an ALV/LSE 5003 correlator over a time window of 30 sec (10 runs of 3 sec) using the software CONTIN. Each measurement was repeated 3 times. The value of *D*_h_ was averaged over 3 independent experiments; error bars indicate the standard deviations for these 3 experiments. For ζ-potential measurements, the solution was transferred to a U-tube cuvette (DTS1070, Malvern, England); the instrument was operated in automatic mode. The electrophoretic mobility of the NPs was measured, from which the ζ-potential value is determined by applying the Henry equation^73^. The ζ-potential values were averaged over 3 independent experiments with 30 runs per experiments; error bars indicate the standard deviations for these *3* experiments.

##### UVVis absorption experiments

UVVis absorption experiments were used to determine the mean number of bound PEG-NLS molecules per NP, 〈*N*〉, and the mean anchoring distance between adjacent PEG-NLS molecules, *ξ^*^*. UVVis absorption experiments were carried out using fluorescently labeled NLS (TAMRA-NLS) and bare NPs of 40 nm in diameter. The concentration of TAMRA-NLS was determined using an extinction coefficient of 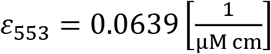. The value of 〈*N*〉 (and *ξ*^*^) were averaged over 3 independent experiments; error bars indicate the standard deviations for these 3 experiments.

##### Western Blot

Western-Blot (WB) was used to confirm the recruitment of importins, and dynein motors by the PEG-NLS coated NPs. In WB experiments, we use NPs of 0.22 μm in diameter, [NLS] is 14.8 μM, and [CE] is 3.4 mg/mL. The NPs were prepared as detailed above, then pelleted at 6800 g for 10 min at 25 °C, resuspended in 1x Laemmli Sample Buffer, and boiled for 5 min to promote detachment of the bound proteins. The proteins were separated by electrophoresis using a 12 % agarose gel, transferred to a nitrocellulose membrane. The membrane was incubated for 1 h in a blocking buffer of PBST (PBS supplemented with 0.1 v/v % Tween) and 10 % (v/w) dry skim milk (Sigma-Aldrich). The membrane was washed 3 times with PBST for 5 min, and then incubated for 1 h at 25 °C with anti-Dynein (Santa Cruz, sc-13524), anti-Karyoprotein α2 (Santa Cruz, sc-55538), or anti-Karyoprotein β1 antibody (Santa Cruz, sc-137016). The primary antibodies were diluted 1: 500 (v/v) in blocking buffer prior use. Then, the membrane was washed 3 times with PBST and incubated with an anti-Mouse HRP conjugated secondary antibody (Santa Cruz, sc-2005) diluted 1: 5000 (v/v) in PBST supplemented with 0.5 % (v/w) skim milk, for 1 h at 25 °C. To finalize the procedure, the membrane was washed 3 times with PBST and incubated with an ECL Western blot Reagent (1705060, Bio-Rad), for 5 min in the dark. Images were collected by chemiluminescence using Fusion FX imaging system (Vilber Lourmat, France).

#### Motility assay experiments

##### Chamber preparation

Flow cells were prepared using a glass slide and a glass coverslip (washed with deionized water, EtOH 70 %, and dried using nitrogen (gas)), and two stripes of warm Parafilm placed in between them, to form a few mm width channel. The chamber was incubated for 5 min with 20 μL (chamber volume) Biotinylated Casein 0.1 mg/ml and washed with 3 volumes of BRB80. Biotinylated Casein is prepared by biotinylation of k-Casein (Sigma-Aldrich) using EZ-Link (Thermo Fisher, 21336). Then, the chamber was incubated with 20 μL of 1 mg/mL Neutravidine (Thermo Fisher, #31000) for 5 min at 23 °C, and washed with 3 volumes of BRB80.

##### Microtubules preparation

Black tubulin from porcine brain, purified via three polymerization/depolymerization cycles^74^, was kindly provided by Uri Raviv (Hebrew University). Purified tubulin was flash-frozen in liquid nitrogen and kept at −80°C until used. Biotin-fluorescently labeled microtubules were prepared according to the protocol described in Christopher Gell et al. work^75^. A tubulin mix containing 51 μM black tubulin, 6 μM Biotin tubulin (T333P, Cytoskeleton), 3 μM Rhodamine tubulin (TL590M, Cytoskeleton), and 3 mM GMPCPP (NU-405S, Jena Bioscience) was mixed on ice, divided, flash-frozen in liquid nitrogen, and kept at −80 °C until used. Prior mix preparation, the black tubulin was thawed and kept on ice for 5 min, then centrifuged for 10 min at 126000 g and 4 °C. The supernatant was kept on ice, and its concentration was determined by absorbance at 280 nm 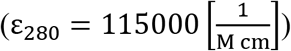. For microtubules (MTs) assembly, an aliquot of the tubulin mix was thawed and diluted with BRB80 to a final concentration of 4 μM, then incubated for 3 h at 37 °C to promote MTs assembly. For motility assay experiments, the MTs were diluted to 40 nM with warm (37 °C) Wash Buffer (WshB) (BRB80 containing 0.02 mM Paclitaxel, 15 mM Glucose, 0.1 mg/mL Glucose oxidase, 0.02 mg/mL Catalase, and 50 mM DTT) and used on the same day.

##### Formation of marked-end microtubules

Marked end microtubules were prepared using bright fluorescent MTs seeds that serve as nucleation sites for the polymerization of dim fluorescent, N-Ethylmaleimide (NEM) modified tubulin^76,77^. NEM modified tubulin was used to inhibit MT seeds minus-end assembly. NEM modified tubulin was prepared by mixing 100 μM black tubulin with 1 mM NEM (Sigma-Aldrich) and 0.5 mM GMPCPP in BRB80 and placing that solution on ice for 10 min. The reaction was quenched by the addition of 8 mM β-mercaptoethanol. The NEM-tubulin mix was incubated on ice for an additional 10 min.

For marked-end MTs preparation, a tubulin mix containing 3.2 μM black tubulin, 4 μM Rhodamine tubulin, 0.8 μM Biotin tubulin and 1mM GMPCPP in BRB80 was incubated for 15 min at 37 °C, to produce short ‘bright’ MT seeds. One volume of bright MT seeds mixed with seven volumes of a ‘dim’ tubulin mix containing 3.6 μM black tubulin, 0.4 μM Rhodamine tubulin, 0.8 μM Biotin tubulin, 3.2 μM NEM modified tubulin, and 1 mM GMPCPP in BRB80, was incubated at 37 °C for 1h to promote polymerization. The marked-end MTs were diluted to 40 nM with warm (37 °C) WshB and used on the same day.

Motility assay experiments. The mean diameter of bare NPs used in the motility assays is 40 nm. For the experiments, 40 nM of microtubules were introduced in the flow cell and incubated for 10 min at 23 °C. The chamber was washed with 3 volumes of WshB to remove unbound microtubules. Prior assay, the NPs were pelleted by centrifugation at 6800 g for 8 min at 4 °C, resuspended in 60 μL Motility Buffer (BRB80 containing 0.02 mM Paclitaxel, 15 mM Glucose, 0.1 mg/ml Glucose oxidase, 0.02mg/ml Catalase, and 50 mM DTT, 0.1 % Methyl cellulose 4000 cP, and 10 mM of Mg-ATP), and incubated on ice for 5 min. Prior insertion in the flow cell, the NPs are brought to 23°C. Samples were excited by total internal reflection illumination at 488 and 568 nm (LAS-AF-6000, Leica Microsystems). Images were captured using an Andor DU-897 EMCCD camera (Oxford Instruments).

#### Data analysis

Particle tracking and image analysis are detailed in the SI Sec. 4. In short, we use the IDL multi-particle tracking method^56^ implemented for MATLAB to automatically detect distinct NP trajectories. Using this method, we extract the 2D center-of-mass coordinates (*X*(*t*), *Y*(*t*)) per time point *t* of the individual NPs. The NP velocity *v* was calculated by taking the NP’s center-of-mass position at times *t* and *t* + Δ*t* and dividing it by Δ*t*. The longitudinal velocity was calculated by projecting the NP velocity on the MT long symmetry axis, 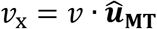, where 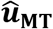 is the MT unit vector. Positive *v*_x_ corresponds to minus-end directed motion. Similarly, transverse motion, 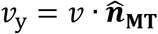, was determined by projecting the NP velocity on the MT normal unit vector, 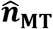. The corresponding transverse step is given by Δ*y* = *v*_y_ · Δ*t*. Right-handed transverse motion is defined as positive and left-handed transverse motion as negative. We use the mean value of transverse steps, 〈Δ*y*〉, to extract the mean angular velocity, 〈*ω*〉, and its error, *ω*_e_, as follows:

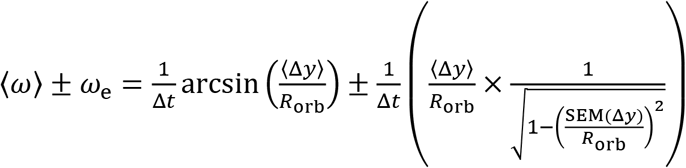

where SEM(Δ*y*) is the standard error of the mean (SEM) value of Δ*y*, and *R*_orb_ ≅ 153 nm is the distance from the MT center and the NP center-of-mass (see SI Sec. 3). Then, we estimate the mean helical pitch, 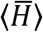, and its corresponding error, H_e_, as follows:

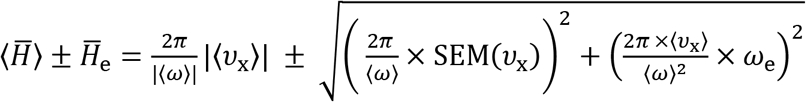

where SEM(*½*_x_) is the standard error of the mean value of *ν*_x_.

Similarly, we can estimate the mean angular velocities for right- (〈*ω*〉|*ω* > 0) and left- (〈*ω*〉|*ω* < 0) handed motions, and the resulting mean helical pitches for the anticipated right- 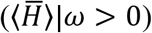 and left- 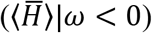 handed helices. Also, the mean angular velocity for separated minus-end directed (〈*ω*〉|*ν*_x_ > 0) and plus-end directed (〈*ω*〉|*ν*_x_ < 0) motions, and the resulting mean helical pitches for the anticipated minus-end directed 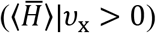 and plus-end directed 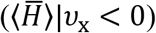 motions.

### Model and simulation algorithm

Our simulations are based on the well-known Monte-Carlo (MC) method (see Fig. S6.1). We simulate processes that occur on timescales much longer than the polymer Zimm relaxation time and the NP rotational and translational timescales. All these processes are associated with thermal diffusion, controlling fluctuations, and are also responsible for relaxation towards mechanical equilibrium. These relaxation times are estimated in SI Sec.7 for completeness.

We simulate the motion of a 20 nm radius NP that is coated with a fixed number of polymers grafted to its surface, each ending with a dynein motor protein. The number of grafted polymers per NP, *N*_m_, is related to the particle radius *R* and the anchoring spacing, *ξ, via N*_m_ = 4*πR*^2^/ξ^2^. To accurately model the experiment, we distribute the polymers randomly over the NP surface. To reduce computation time, we do not consider the actual dynein structure. Instead, we treat the dynein as a simple cylindrical-like body, with a radius of 10 nm and a height of 60 nm. The height is an approximation that is built from the accumulated dimensions of the dynein components and the *αβ* importin complex (SI. Sec. 4). The reasoning behind the choice of 10 nm radius is explained below.

The MT surface, over which the motors step, is modeled as a cylinder with a 12.5 nm radius. The MT surface is covered with binding sites for the dynein MTBDs. To simulate the motion of the dynein pivot location, we used the MT binding site array for the MTBDs (as presented in previous work^39^) to create an artificial MT-dynein (i.e. dynein pivot) binding site grid. We did it by sampling all the possible configurations of an MTBD pair, and for each of the configurations, we calculated the corresponding location of the dynein pivot.

The simulation algorithm which describes a single NP trajectory, includes a sequence of events, beginning with the binding of a single motor to the MT, and ending when the last motor detaches (henceforth, we use the term “motor” to refer to the dynein-polymer complex). Each simulation iteration (MC-step) involves one of the following competing events that is allowed for each of the motors: (i) motor binding (for the unbound motors), (ii) unbinding (for the bound motors), or (iii) stepping (for the bound motors); the latter might lead to a jamming event as discussed below. The probability of each event *i* is proportional to its rate, *k*_i_. To avoid a non-vanishing, computation time consuming, probability for no occurrence of any of the three events, we choose all event probabilities, for each MC-step, to sum up to unity, such that the probability of an event *i* is given by 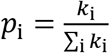. With this choice, the physical time corresponding to an MC-step is 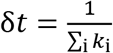, implying that this time increment (unlike common MC simulations) varies between different MC steps. For convenince, we separate the events to the above three groups such that the rate for an event class is 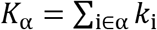, with α= (binding, unbinding, and stepping), such that the sum on i ∈ α runs over the different participating motors; the corresponding probability for each event class α is 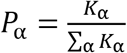. Thus, after an event group has been randomly selected with probability *P*_α_, a second random selection is performed for the actual event within the group with probability 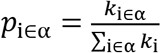. For instance, a random selection of a “binding” process (out of the three processes) is followed by the selection of the motor identity out of the unbound motors. The above procedure is entirely equivalent to a random selection of an event without the prior selection of one of the three classes, since *p*_i_ = *P*_α_ *p*_i∈α_. After each such MC-step, the NP is balanced mechanically by minimizing the free-energy, mimicking the relaxation processes mentioned above that are orders of magnitude faster than the MC physical timescale (see estimated time scales in SI Sec. 7).

In the following, we shall discuss the free-energy calculations, rate calculations, and the NP stepping algorithm.

#### Stepping Model for Mammalian Cytoplasmic Dynein

In a previous publication^39^, we presented a stochastic model for the 2D stepping of single cytoplasmic *yeast* dynein, which shows excellent agreement with experimental results. Here, we use a similar model for single *mammalian* dynein. Although these two motor proteins differ in several aspects, the conversion of our *yeast* dynein stepping model to the *mammalian* case can be achieved using a few assumptions and additional experimental data.

The main assumptions concerning the conversion of the (single) *yeast* dynein stepping model to *mammalian* dynein are: **(i)** Both *yeast* and *mammalian* dynein MTBDs bind to the same binding sites on the MT surface. **(ii)** All the required biological factors, such as bicaudal-D-homolog-2 (BICD2) and dynactin, are at optimal concentrations, allowing to neglect their binding-unbinding kinetics. **(iii)** The stepping vector probability distribution, P(L), for both dynein types, is written as a product of two functions:

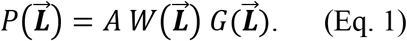

where 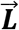 is the dynein step vector (with longitudinal and transverse components *L*_x_ and *L*_y_, respectively), and A is a normalization factor. 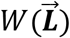 is a motor-specific function that accounts for the stepping model rules^39^ and is a symmetric function (i.e., invariant under the transformation 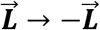); thus, its projection along *L*_y_, i.e., 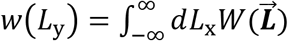, is also a symmetric function of its variable. 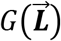 is a force exertion function that includes the internal (longitudinal) motor force, equal in magnitude to the so-called stalling force *F*_s_, thus breaking the symmetry between forward (minus-end directed) and backward (plus-end directed) steps. It also may include any external potential 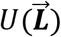 (with the definition *U* (0) = 0). It is given by

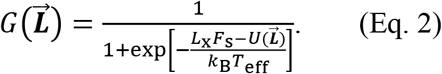

where *T*_eff_ is an effective temperature fit parameter. In particular, as considered in previous works^31,39^, for a longitudinal backward (plus-end directed) pulling force (defining it with F > 0) 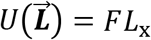 and

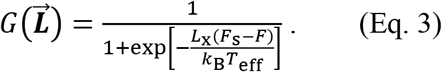

The above assumptions are not sufficient for the model adjustments, i.e., obtaining 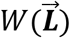, as for the *yeast* dynein case. For example, to the best of our knowledge, for the *mammalian* dynein case, no data is published on the inter-MTBD stretching distance distribution and/or its angular distribution. Therefore, to complete the model, we use here the very recent single-molecule results on *mammalian* dynein stepping statistics of Yildiz and co-workers^37,55^. To do so, we consider only the experimental data regarding the BICD2 adapter, as this was shown to bind mainly single dynein (~85 %)^37^ We use the experimental data for the longitudinal step-size distribution of the dynein pivot under stalling conditions, i.e. *F* = *F*_s_ in Eq. (3). Under such conditions, our model reads 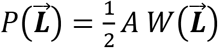. Yet, the provided data^37,55^ only entails the distribution of the longitudinal component *L*_x_. Thus, to obtain 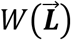, we build on the assumption (i) described in the previous paragraph, which implies that the one-to-one correspondence between longitudinal and transverse step components (*L*_x_ and *L*_y_, respectively) is identical for both *yeast* and *mammalian* dynein motors. Using this correspondence, we can infer the distribution of the *mammalian L*_y_ from its *L*_x_ distribution and, in turn, its 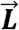 distribution. In addition, to artificially reach adequate data sampling of the experimental step distribution, we build on the symmetry discussed above for 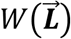, which implies that ideally the raw data^37,55^ should be composed of pairs of 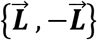 abundance. We observed that in the raw data some pairs are not completed (probably due to the statistical error associated with finite-size sampling of the side steps, and not due to experimental issues). Thus, we have forced the symmetry over the given data by completing the lacking pairs.

Next, we obtained the fit parameter *T*_eff_ by adjusting its value such that under no external force, *F* = 0, the plus-end directed and minus-end directed steps fractions are consistent with the measured values^37^: 23 % and 77 %, respectively, leading to *T*_eff_ = 1900 K. To validate the mentioned adjustments, we compared our predictions for the mean step-size with the data^55^ for different *F* values, see Table 6. Note the consistency of the experimental data with our single cytoplasmic *mammalian* dynein model predictions (within theoretical and experimental errors).

**Table 6.** Comparison between experimental and theoretical mean step-size of a single dynein (*mammalian-type*). Since the experimental results are affected by the FIONA method resolution of ±1 nm, the error in the experimental results is estimated as 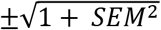. We believe that the small mismatch between model and experiment at *F* = 0 is likely due to an experimental error at vanishing laser trapping forces^2^.

| $F$ [pN] | Experimental mean step-size | Theoretical mean step-size |
| --- | --- | --- |
| 0 | $11.4 \pm 1.1$ nm | 9.1 nm |
| 1.5 | $5.4 \pm 1.5$ nm | 6.9 nm |
| 2.3 | $4.3 \pm 1.5$ nm | 5.2 nm |
| 2.9 | $2.5 \pm 1.5$ nm | 3.3 nm |
| 3.6 | $-0.5 \pm 1.5$ nm | 0.3 nm |

#### Application of the single-motor stepping model to the NP motion

The single *mammalian* dynein stepping model described above, which builds on our previous work on single *yeast* dynein stepping model and uses the experimental data of Yilditz and co-workers^37,55^, can be readily applied to any of the NP motors that are attached to the MT and attempt to step. Let 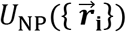 be the NP Helmholtz free-energy for a given configuration of MT bound motors 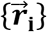 (denoting the bound motors locations), in which the NP is in mechanical equilibrium, 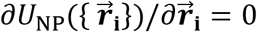 (for all bound motors i). The zero of energy is defined, for convenience, when there are no motors bound to the MT. To apply this free-energy to Eq. (2), we assume that after a step is performed, mechanical equilibrium is instantaneously restored, as described above, and the NP coordinates are updated accordingly. Hence, we identify the “external potential” in Eq. (2) as the difference in the NP Helmholtz free-energy upon stepping of a single motor, 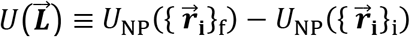, where 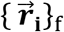 and 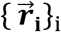 are the final and initial bound motor configurations.

##### Motor free-energy

We assume that the anchoring density is in the so-called “mushroom regime”^53^ (consistent with the experimental densities) such that *ξ* > *R*_g_. Furthermore, upon binding of a motor to the MT surface followed by stepping, the polymer becomes stretched such that the density of monomers in the transverse direction to the polymer end-to-end vector is reduced. Hence, polymer-polymer excluded volume interaction is not accounted for, and regarding the free-energies calculation, we can consider single polymer theory.

We choose a planar surface that is tangent to the spherical NP surface at the polymer anchoring position, see Fig. 11A. When the dynein at the polymer free-end (henceforth “dynein-polymer-end”) binds to a site on the MT surface, the polymer-end can be regarded as (temporarily) fixed in space, with position dictated by the dynein and importins dimensions (see Figs. 2C and 11C). Thus, dynein-polymer-end position is always residing on a virtual cylindrical surface whose cross-section radius (R_2_ in Fig. 11C) is the sum of the MT radius, the total dynein length, and the two importin protein sizes (SI Sec. 4).

**Figure 11.**
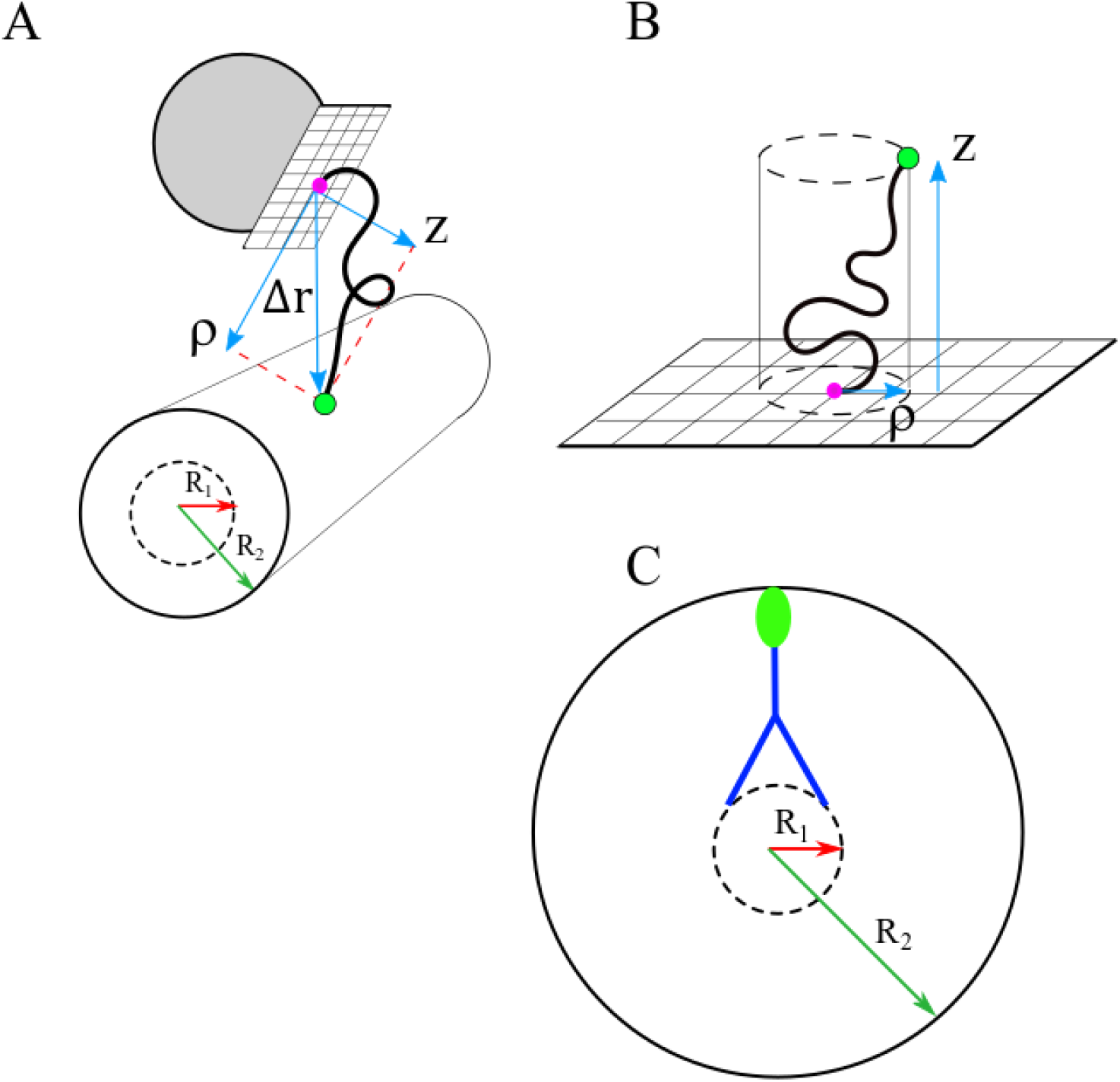
Illustration of the cylindrical coordinates (*ρ*, z). Note that the effective radius of the cylinder, *R*_2_ (green arrow) entails the MT radius, *R*_1_ (red arrow), the vertical length of the dynein (blue lines), and the vertical length of the *αβ* importin complex (green ellipse). Therefore, the value of *R*_2_ is taken as 60 nm (the length estimation is discussed in detail in SI Sec. 4). **(A)** Illustration of the coordinates for the case of a polymer that connects between a NP (gray circle) and the *αβ* importin complex (green dot) that in turn, is connected to the dynein tail domain. **(B)** Illustration of the coordinates for the general case. **(C)** Illustration of the cylindrical cross-sections of (A).

To describe the free-energy change upon a binding event, we use a function that accounts for the entropy loss of a polymer upon binding, and the dynein binding energy gain, *ϵ* > 0. Using cylindrical coordinates 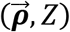 associated with this tangent surface (Fig. 11A,B), the polymer anchoring position is set at the origin (*0,0*), and the polymer other end, which is not anchored to the surface, is fixed at 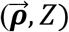.

This leads to the following equation for the free-energy *difference, U*_motor_, between a fixed and free-end polymer^78,79^:

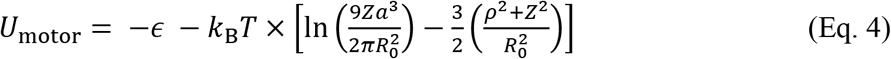

where a is the polymer Kuhn length (~0.76 nm for PEG^80,81^) and 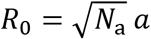 is the free-polymer end-to-end distance (*N*_a_ is the number of Kuhn segments) (see SI Sec. 2). We approximate *ϵ* to be 8 kBT^64^

Note that this free-energy approximately accounts for the excluded volume interaction between the whole polymer and the NP rigid surface (assuming *R*_g_ ≪ *R*). It does not account for the excluded-volume interaction between the polymer and the MT surface. However, since the contour length of the PEG is about 30 nm, it is evident that no monomer can ever reach the MT itself. The excluded-volume interaction between the polymer and the body of the dynein-*αβ* complex is neglected since – for similar considerations – the only domains where it could be relevant are the dynein tail and the two importins whose physical dimensions are relatively small.

As each polymer in anchored at a different position on the NP surface, its anchoring tangent plane is orientated differently with respect to the MT surface. Thus, we perform a coordinate transformation for each polymer to obtain its specific *U*_motor_. In Eq. (4), *ρ*^2^ + *Z*^2^ stands for the square of the (actual) end-to-end vector 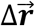 (Fig. 11A,B), which we calculate for each MT bound motor. This can be conveniently done by transforming to lab frame Cartesian coordinate system, redefining the origin at the NP center, and using 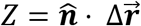 where 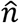 the unit vector normal to the NP surface at the anchoring position (pink dot, Fig. 11A,B).

##### Motor Stepping rate

For a single free motor, the mean stepping rate for any step-size 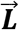, *k*_step_, can be estimated from *k*_step_ = 〈*ν_x_*〉/〈*L*_x_〉, where *ν*_x_ is the longitudinal velocity [nm/sec], and *L*_x_ is the longitudinal step-size [nm]. For *mammalian* cytoplasmic dynein, we set 〈*ν*_x_〉 to be the well-established value 800 nm/sec, and 〈*L*_x_〉 to be about 9 nm. This implies that the step-size dependent stepping rate is 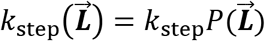, where 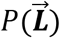 is the *normalized* step-size distribution defined in Eq. (1). Thus, consistently, 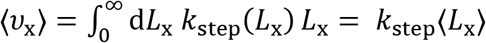. Note that *k*_step_ includes the dwell time.

##### Motor Binding/Unbinding rates

As described in Ref.^16^, we assume that the binding-unbinding rates of the different motors obey detailed balance relations. Therefore, the pair of rates, *k*_+_ – for binding of a single motor, and *k*_–_ – for the unbinding of the same motor, are assumed to obey the ratio 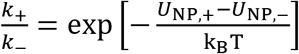, where 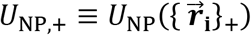 and 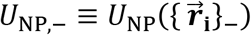 are the free-energies of the binding and unbinding states, respectively. In order to uniquely specify the two states, there is (as usual) a need for another condition, and (consistent with common choices in stochastic processes) we take *k*_+_ + *k*_–_ = *τ*^-1^, and adjust the free-parameter *τ*^-1^ such that when the NP is left with a single motor it unbinds with the experimentally known rate *k*_0_ = 1 Hz^31^. As show in^16^, this leads to the following rate expressions:

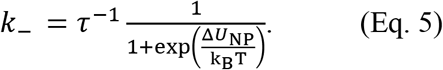

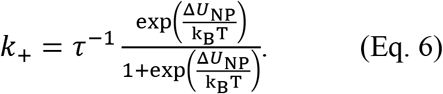

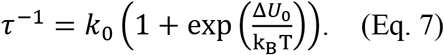

where Δ*U*_NP_ = *U*_NP,+_ – *U*_NP,–_ is NP free-energy difference between a (single motor) binding and unbinding states, and Δ*U*_0_ is associated with the special case where the NP entails only a single motor. Note again that Δ*U*_NP_ accounts for the free-energy change of the whole NP (upon a single binding event) of any one of the motors out of all the NP anchored motors, and when both states are assumed to be at mechanical equilibrium.

##### Motor-motor excluded volume interaction

When a motor attempts to step into a binding site, it may not be accessible due to the “excluded volume interaction” between the attempting motor and the neighboring bound motors and we term this as a “jamming event”. If a jamming event occurs, the attempting motor rests still, implying that the temporal NP longitudinal and angular velocities (ν_x_ and ω) both vanish. We determine the motor-motor excluded volume interaction using the following approach. First, the volume occupied by a single MTBD is described as that of a sphere of 4 nm radius^82^ (Fig. 12), which allows determining the excluded volume created when two MTBDs meet, i.e. the volume of a sphere of the MTBD diameter. Note that the known MTBD measured dimensions, 3.6 nm, are likely smaller than those we use; this practice ensures that we do not underestimate the volume occupied by the MTBD.

**Figure 12.**
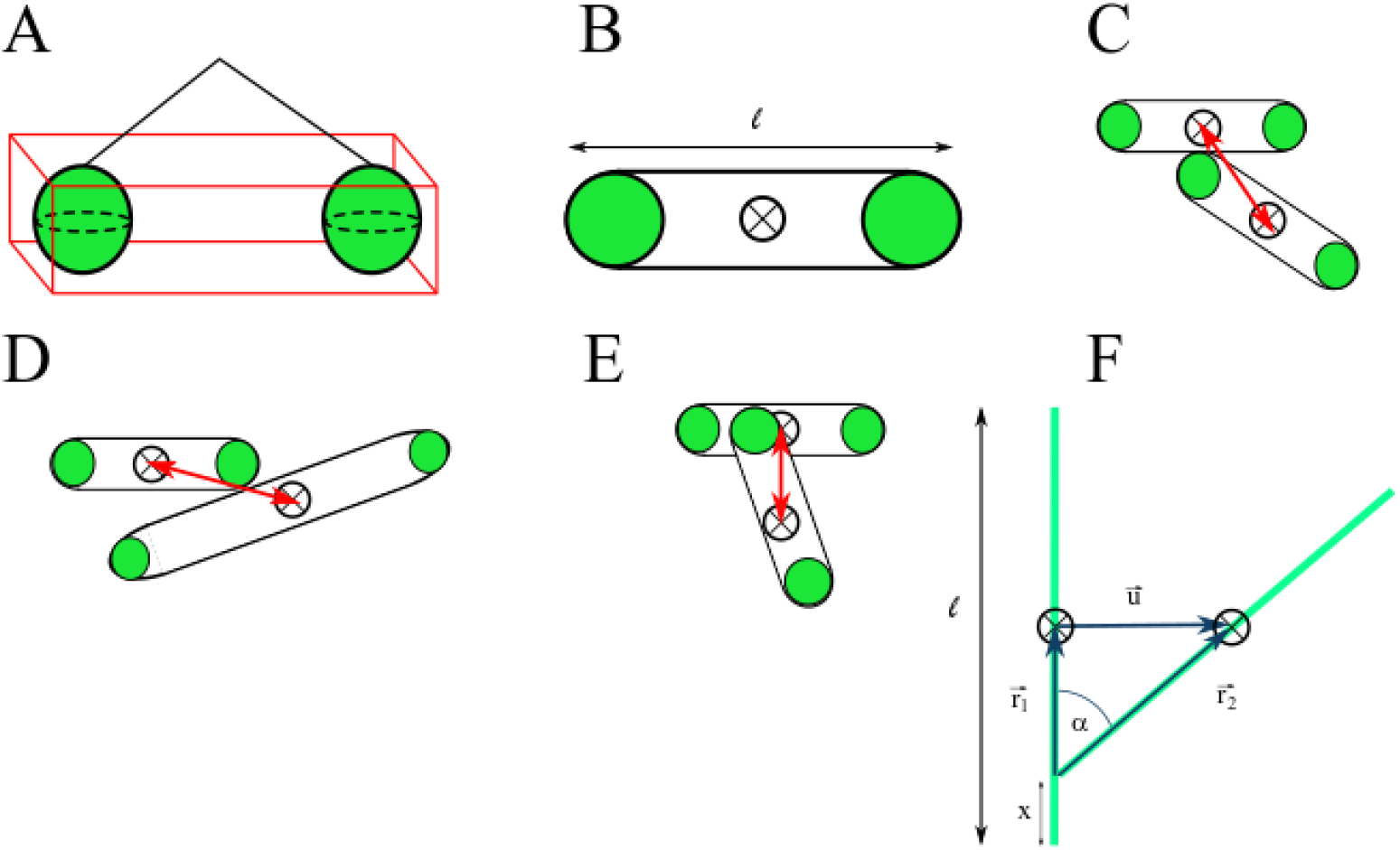
Illustration of the excluded volume and possible configurations of two MTBD pairs. MTBDs are represented by green spheres. The dynein pivots are represented by ⊗, and the distance between two dynein pivots (*d_jam_*) is represented by the red arrows. **(A)** Illustration of the volume occupied by a MTBD pair and the space in between the pair’s MTBDs (red square). **(B)** Illustration of the projection of (A). **(C-D)** Two examples of adjacent MTBD pairs. **(E)** Example of two MTBD pairs in “crossing” configuration. **(F)** Schematic illustration of a possible configuration of two MTBD pairs. MTBDs pairs are represented by green lines (of length *ℓ*), and the dynein pivots are represented by ⊗.

Second, let us now define *V*_pair,〈ij〉_ as the volume occupied by the 〈*ij*〉 MTBD *pair*, where the space between the MTBDs is also accounted (Fig. 12A). Thus, a jamming event occurs whenever the volumes of two pairs (corresponding to two motors), *V*_pair,〈ij〉_ and *V*_pair,〈km〉_ (〈ij〉 ≠ 〈km〉), overlap. However, since our model does not consider the actual dynein structure, we determined the average distance between two motors that leads to a jamming event, d_jam_ (Fig. 12C-E). We represent each pair as a projection on a 2D surface, or in other words as a ribbon of length *ℓ*, capped with two hemi-circles (Fig. 12B).

Next, we estimate 〈d_jam_〉 using the following calculation (Fig. 12C,D). We consider two such ribbons at contact (Fig. 12F), making an angle *α* between them. The vector positions of the two ribbon centers obey, 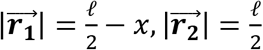. The vector connecting the two ribbon centers is 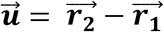. Therefore 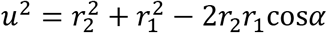, leading to 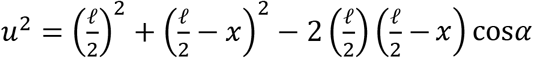. The configurational average of this distance is 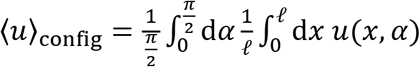, which can be evaluated numerically to give 〈*u*〉_config_ ≅ 0.543 *ℓ*. As the values of *ℓ* can vary, we use its mean, namely 〈*d*_jam_〉 = 0.543 〈*ℓ*〉 = 12 nm.

Note that this approximation does not account for crossing configurations (Fig. 12E). Thus, 〈*d*_jam_〉 = 12 nm can only be taken as an upper bound value estimate. To estimate a lower bound value, we need to account for crossing configurations. However, as far as we know, there is no data concerning dynein-dynein crossing configurations, so we only examine a simple crossing case in which the likelihood for steric interferences is relatively low; we consider the case where one MTBD resides between another pair of MTBDs belonging to a neighboring motor (Fig. 12E). We use the same method as above, but without accounting for the MTBDs dimension, resulting in shorter ribbon length, *ℓ*\* = *ℓ* – 2*d*_MTBD_ (where *d*_MTBD_ = 4 nm is the “MTBD sphere” radius) such that 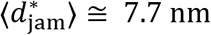. Since the real value of *d*_jam_ resides between the lower and the upper estimated bounds (i.e., between 7.7 and 12 nm), we use in our simulations 〈*d*_jam_〉 = 10 nm. Consequently, regarding the excluded volume, our motors are modeled as standing cylinders, with a cross-section diameter of 10 nm and vertical length as estimated in SI Sec. 4.

## Author Contributions

Anne Bernheim-Groswasser – development of experimental setup model, analysis of results, writing the article.

Rony Granek – development of the theoretical model, analysis of results, writing the article.

Itay Fayer – development of the theoretical model and computer code, analysis of results, writing the article.

Gal Halbi – development of experimental model and carry-out of experiments, analysis of results, writing the article.

Dina Aranovich – development of experimental set-up model and carry-out of experiments and analysis of results.

Shachar Gat – development of experimental set-up model and carry-out of experiments and analysis of results.

Shy Bar – development of experimental set-up model and carry-out of experiments.

Vitaly Erukhimovitch – analysis of results.

## Acknowledgments

This research was initially supported by the Focal Technological Area Program of the Israeli National Nanotechnology Initiative (INNI). We are highly grateful to Prof. Ahmet Yildiz for sharing with us the data associated with Ref^37,55^. We are grateful to Prof. Uri Raviv for providing purified tubulin.

## Supplementary Information

### SI. 1 NP Synthesis and Characterization

The NP synthesis process entails several consecutive steps, where at each step, a single component is added (see Fig. 1A, main text). Bare NPs saturated with Neutravidin were conjugated with Biotin-PEG-thiol spacers, then incubated for a short time in a solution containing SV40 T large antigen NLS peptides at variable concentrations; see Materials and Methods section. The resulting NPs were characterized using Cryo-TEM, dynamic light scattering (DLS), and zeta potential. In addition, adsorption isotherms from UVVis experiments yield the surface density and the mean number of grafted Biotin-PEG-thiol that are end-conjugated by NLS (PEG-NLS), 〈*N*〉. The latter is transformed into a mean anchoring distance between neighboring PEG-NLSs, *ξ*\* (Fig. 1B, main text). For the experiments we use bare NPs of different diameters. By using the same total surface area of bare NPs in the experiments, we assure that the mean anchoring distance is independent of the NPs diameter. Next, the NPs were incubated in (Hela) cell extract, allowing the recruitments of *α*- and *β*-importins, followed by *mammalian* dynein association; the recruitment of the importins and dynein is verified *via* Western Blot (Fig. 1C, main text). The particles are washed from the excess cell extract, incubated in an ATP solution, and then injected into a flow cell in which MTs are adsorbed and immobilized on a glass surface.

Several measurements were carried out to verify the NPs integrity and to study the various components effect on the NP characteristics, i.e., the complete integration of the NP and the different components. First, to study the effect of the grafted Biotin-PEG-thiol, we examined the distances between adjacent NPs using Cryo-TEM^1^. We compared cryo-TEM microscopy images of a control group without grafted Biotin-PEG-thiol (Fig. S1.1A) and NPs with grafted Biotin-PEG-thiol molecules (Fig. S1.1B). The images show an NPs in which inter-particle distances depend on the absence or presence of grafted Biotin-PEG-thiol molecules. Without Biotin-PEG-thiol, the mean spacing between the NPs is 10.5 ± 7 nm (mean ± STD), typically twice the diameter of a Neutravidin molecule^2^, and with a *M*_w_ = 5 kDa Biotin-PEG-thiol, it is 26 ± 5 nm mean ± STD), which is much larger than the theoretical value of the (free polymer) gyration radius: *R*_g_ = 2.27 nm (see SI.2 below). This is consistent with weak attractive entropic forces (e.g., depletion attraction) working against the entropic repulsion resulting from the anchored chains^3^. Note that the black dots on the particle’s surface are the Neutravidin molecules (Fig. S1.1).

Since Neutravidin diameter is about 5 nm^2^, and since the Neutravidins are closely packed on the NP surface, we conclude that the anchored Biotin-PEG-thiol molecules are effectively in the so-called “mushroom regime”^4^

**Figure S1.1.**
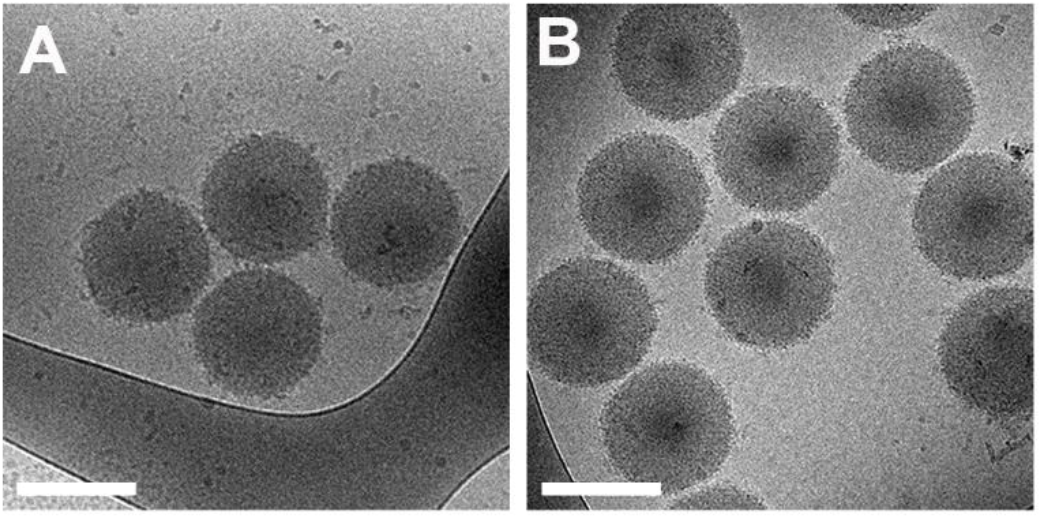
Cryo-TEM microscopy images of Neutravidin-coated NPs without **(A)** and with **(B)** grafted 5 kDa Biotin-PEG-thiol. Mean bare NPs diameter is 196 nm. Bars are 200 nm.

To follow the NP decoration during the various stages, we performed dynamic light scattering (DLS) experiments. The data show a significant increase in the NP hydrodynamic diameter (*D*_h_) during the first two decoration steps: 245 ± 10 nm (Stage I) and 410 ± 4 nm (Stage II) compared to the bare NPs, which have *D*_h_ = 198 ± 3 nm (all values correspond to mean ± STD). At the third step (NLS binding), a slight decrease is detected, confirming the Biotin-PEG-thiol binding to the NP.

Next, to conclude on the NP charge at the various decoration steps, we examined the zeta potential^5^ (see Materials and Methods section). The zeta potential is highly negative for the bare NPs and has a value of −41.4 ± 1 mV (mean ± STD) (i.e., the bare NPs are negatively charged), and it remains negative, but gradually decays (in absolute values) as NP decoration advances. Regarding stages I and III, this trend is consistent with the assumption that both the Neutravidin and NLS are positively charged; thus, a reduction of the (absolute value of) the zeta potential is expected after their binding. At the end of stage III (NLS binding), the zeta potential equals −9.6 ± 0.5 mV (mean ± STD).

Finally, to determine the mean number of bound PEG-NLS molecules per NP, 〈*N*〉, and the mean anchoring distance between adjacent PEG-NLS molecules, *ξ*\*, we carried out UVVis absorption experiments using fluorescently labeled NLS, TAMRA-NLS (see Materials and Methods section). We incubated Biotin-PEG-thiol-grafted NPs with increasing amounts of TAMRA-NLS and measured the adsorption of the remaining TAMRA-NLS molecules that *did not* absorb to the NPs (i.e., supernatant). From the absorbance, we deduced the number of TAMRA-NLS molecules that *did* absorb to the NPs,

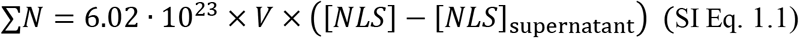

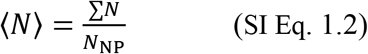

Where *V* is solution volume, [*NLS*] is the concentration of TAMRA-NLS in the incubation solution, [*NLS*]_supernatant_ is the concentration of TAMRA-NLS that remained in the bulk and did not absorb to the NPs, 6.02 · 10^23^ is the Avogadro number, ∑^*N*^ is the total number of TAMRA-NLS molecules that were absorbed to the NPs surface, *N*_NP_ is the total number of NPs – shown in Fig. S1.2 the results for bare NPs of 40 nm in diameter. Similar results are obtained with bare NPs of 196 nm (data not shown).

The dependence of 〈*N*〉 on the concentration of TAMRA-NLS (NLS in Fig. S1.2) follows a Langmuir-like isotherm. We used the parameters extracted from the fit to estimate the mean number of bound motors per NP for [*NLS*] of 0.05 and 0.025 μM, which are below the detection limit of our UVVis set-up (grey dot in Fig. S1.2). Our motility assays are performed with [*NLS*] varying between 0.025 and 0.3 μM, implying that 〈*N*〉 varies between 5 and 37.

**Figure S1.2.**
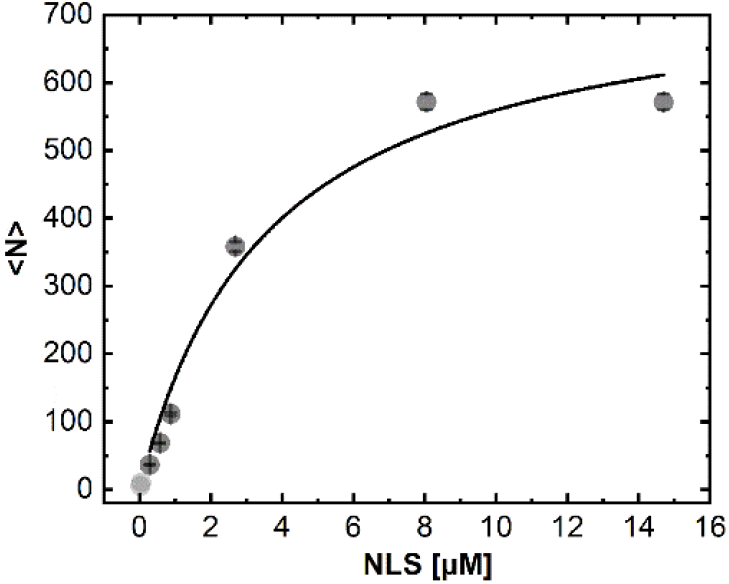
Dependence of the mean number of grafted PEG-NLS, 〈*N*〉, against the concentration of TAMRA-NLS (marked as NLS) follows a Langmuir-like isotherm (grey dots – experimental data, line - fit to the experimental data; *R*^2^ = 0.96). The two bright grey dots correspond to extrapolated values of 〈*N*〉 calculated from the fit, for [*NLS*] = 0.025 and 0.05 μM. Error bars indicate the standard deviations for 3 experiments. Bare NPs have a mean diameter of 40 nm.

Finally, we deduce the mean anchoring distance between adjacent PEG-NLS molecules, *ξ*\*, from 〈*N*〉 assuming that each PEG-NLS molecule occupies a square lattice *ξ*\*^2^,

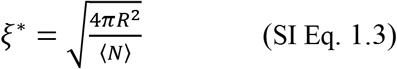

where *R* = 25 nm is the mean radius of the Neutravidin-coated NPs. The dependence of *ξ*\* on [*NLS*] is depicted in Figure 1B (see main text).

As a final step, we used Western-Blot (WB) to confirm the recruitment of importins and dyneins by the PEG-NLS coated NPs (Fig. 1C, main text). These results confirm that in the absence of NLS, the dynein machinery is not recruited to the NP surface.

### SI. 2 Estimation of the PEG contour length, *C*_L_, and Radius of Gyration, *R*_g_

In our motility experiments, we use biotin polyethylene glycol thiol (Biotin-PEG-thiol) with a molecular weight of 5 kDa. The molar mass of each ethylene glycol subunit, the building block of the Biotin-PEG-thiol polymer spacer (marked by the red rectangle, Fig. S2.1), is 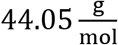. The number of repeating ethylene glycol subunits, n, for a 5 kDa Biotin-PEG-thiol, is 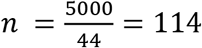.

**Figure S2.1.**
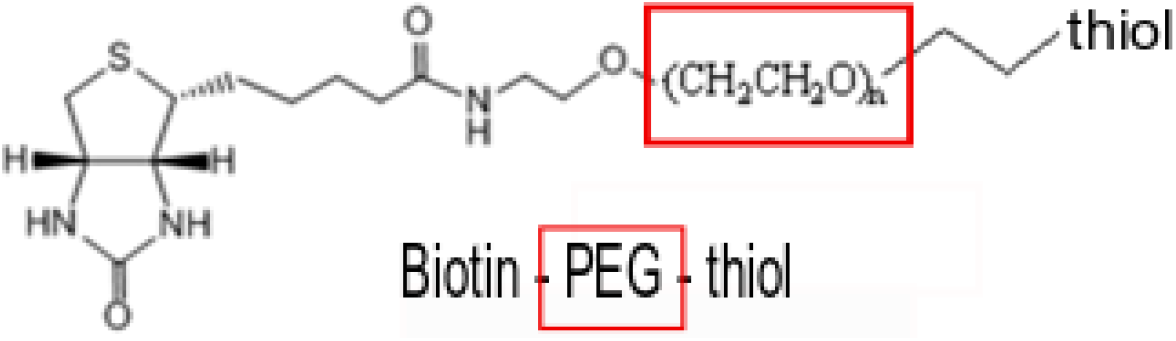
Molecular structure of the biotin polyethylene glycol thiol (Biotin-PEG-thiol) used. *n* refers to the number of ethylene glycol units per PEG polymer.

To estimate the Biotin-PEG-thiol contour length *C*_L_ and radius of gyration, *R*_g_, we use the length of an ethylene glycol subunit *d*_EG_^6^ and the number of ethylene glycol subunits, *n*, as follows^7–9^:

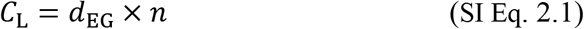

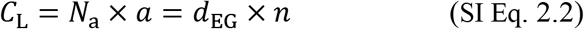

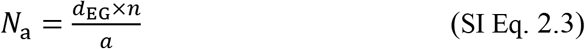

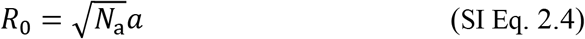

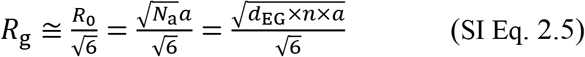

Where *R*_0_ is the polymer end-to-end distance (assuming that the PEG polymer behaves like a Gaussian chain^10,11^), *a* = 0.76 nm is the PEG Kuhn length^10,12^, *N*_a_ is the number of Kuhn length segments, and *d*_EG_ = 0.358 nm^6,10^. Using these parameters we find that *C*_L_ = 40.8 nm and *R*_g_ = 2.27 nm.

### SI. 3 Experimental Raw Data

**Table S3.1.** System V: NPs experimental mean velocities, run-times, and run-lengths (mean ± SEM). *ν* is the absolute velocity, *ν*_x_ is the longitudinal velocity, *ν*_y_ is the transverse velocity, *τ*_p_ is the processivity time, *λ*\* is the absolute and total distance that the NP covered regardless of the motion direction, and *λ* is the total longitudinal run length in the direction of the MT minus-end. The bare NPs mean diameter is 40 nm.

|  | V |
| --- | --- |
| $[NLS]$ [$\mu\text{M}$] | 0.3 |
| $\langle N \rangle$ | 36.6 |
| $[CE]$ [$\mu\text{M}$] | 6.8 |
| #steps | 536 |
| $\langle \vert v\vert \rangle$ [nm/sec] | $320 \pm 20$ |
| $\langle v_x \rangle$ [nm/sec] | $72 \pm 22$ |
| $\langle v_x \rangle \vert v_x > 0$ [nm/sec] | $253 \pm 20$ |
| $\langle v_x \rangle \vert v_x < 0$ [nm/sec] | $-319 \pm 43$ |
| $\langle v_y \rangle$ [nm/sec] | $39 \pm 10$ |
| $\langle v_y \rangle \vert v_y > 0$ [nm/sec] | $132 \pm 9$ |
| $\langle v_y \rangle \vert v_y < 0$ [nm/sec] | $-162 \pm 18$ |
| $\langle \tau_p \rangle$ [sec] | $23.3 \pm 4.1$ |
| $\langle \lambda^* \rangle$ [nm] | $7700 \pm 1100$ |
| $\langle \lambda \rangle$ [nm] | $1700 \pm 600$ |
| Fractions of forward & backward steps | $0.67 \text{ f} \vert 0.33 \text{ b}$ |
| Fraction of Right- & Left-handed steps | $0.67 \text{ r} \vert 0.33 \text{ l}$ |

**Table S3.2.** Standard deviations, **STDs, for the experimental velocity, run-time, and run-length values of systems I to V**. *ν* is the absolute velocity, *ν*_x_ is the longitudinal velocity, *ν*_y_ is the transverse velocity, *τ*_p_ is the processivity time, *λ*\* is the absolute accumulated distance that the NP covered regardless of its direction, and *λ* is the total longitudinal run length in the direction of the MT minus-end. The bare NPs mean diameter is 40 nm.

|  | I | II | III | IV | V |
| --- | --- | --- | --- | --- | --- |
| $[NLS]$ [$\mu\text{M}$] | 0.025 | 0.05 | 0.05 | 0.3 | 0.3 |
| $[CE]$ [$\mu\text{M}$] | 3.4 | 3.4 | 3.4 | 3.4 | 6.8 |
| #steps | 80 | 62 | 378 | 154 | 573 |
| $\langle\delta\vert v\vert \rangle$ [nm/sec] | 168 | 126 | 542 | 288 | 477 |
| $\langle\delta v_x\rangle$ [nm/sec] | 605 | 386 | 567 | 326 | 518 |
| $\langle\delta v_y\rangle$ [nm/sec] | 226 | 192 | 454 | 193 | 234 |
| $\langle\delta\tau_p\rangle$ [sec] | 1.61 | 2.13 | 2.82 | 6.95 | 16.24 |
| $\langle\delta\lambda^*\rangle$ [nm] | 1055 | 1436 | 4250 | 3610 | 4340 |
| $\langle\delta\lambda\rangle$ [nm] | 932 | 1277 | 3450 | 3750 | 2390 |

**Figure S3.1.**
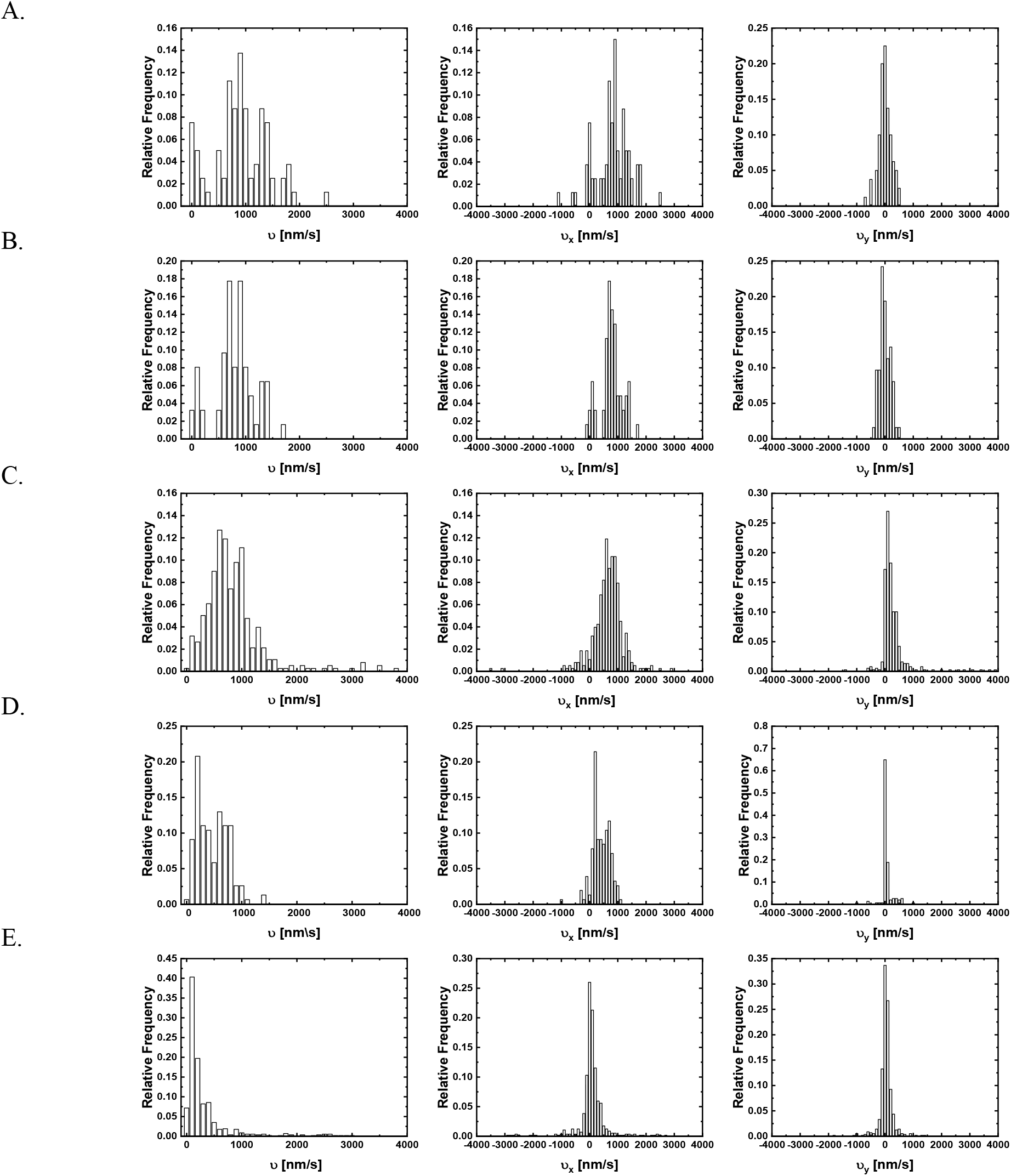
Histogram of the experimental velocities for systems I to V (rows A-E respectively), where *ν* is the absolute velocity, *ν*_x_ is the longitudinal velocity, and *ν*_y_ is the transverse velocity. The bare NPs mean diameter is 40 nm.

### SI. 4 Particle tracking algorithm for NP motion quantification

We use the IDL multi-particle tracking method^13^ implemented for MATLAB to automatically detect distinct NP trajectories. Using this method, we extract the 2D center-of-mass coordinates (*X*(*t*), *Y*(*t*)) per time point *t* (frame) of the individual NPs, from which we determine the individual NP trajectory over time, NP(*X, Y*)_t_. We carry out an initial (automatic) filtering where we select trajectories that **(i)** include at least 6 motion steps and **(ii)** have a travel distance of at least 6 pixels (pixel ≅ 230 nm). Next, we to perform a more delicate (manual) filtering where we exclude NPs, or NP trajectories, that fall under one or more of the following categories: (i) The NP shape is not symmetric or too large, which implies NPs aggregation, (ii) the NP is moving in a region where the density of MTs is too dense, and thus, it is impossible to discern between individual MT tracks, and (iii) the NP moves only for a short period and then get stuck along the MT track for a long period of time. Note that there are cases where the NPs are moving continuously between crossing MTs tracks. In that case, we extract the trajectory on each of the MT tracks separately. The overall run time and run length are calculated from the accumulated traveled time/distance. For NPs moving between crossing MTs, the overall run time and run length correspond to the accumulated traveled time/distance of all individual trajectories.

The NP velocity *ν* was calculated by taking the NP’s center-of-mass position at times *t*, (*X*(*t*), *Y*(*t*))_NP_ and *t* + Δ*t* (*X*(*t* + Δ*t*), *Y*(*t* + Δ*t*)), and dividing by Δ*t*, that is 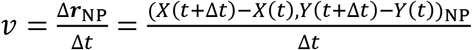, where *X* and *Y* refer to lab frame Cartesian coordinates. The longitudinal velocity, *ν*_x_, was calculated by projecting the NP velocity *ν* onto the MT direction (By definition, positive direction points towards the MT minus-end), i.e., the unit vector of the MT (along its long axis), 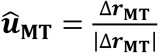, where 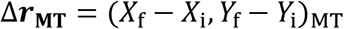. The indexes *i* and *f*, refer to the NP trajectory initial and final time points, and (*X*_i_, *Y*_i_)_MT_ and (*X*_f_, *Y*_f_)_MT_ are the corresponding MT center-of-mass coordinates. Thus, the longitudinal velocity 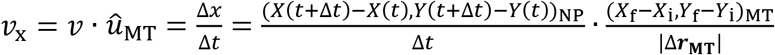, where 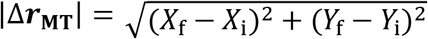. Positive *ν*_x_ corresponds to a NP step Δ*x* in the direction of the MT minus-end, whereas negative values of *ν*_x_ refers to a NP step towards the MT plus-end.

Similarly, the transverse motion, v_y_, was determined by projecting the NP velocity v onto the normal unit vector of the MT, 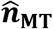. Thus, the transverse velocity 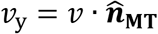 and the corresponding transverse step is given by Δ*y* = *ν*_y_ · Δ*t*. Note, that for each MT, or MT unit vectors 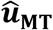, there are two opposite normal unit vectors that can be assigned, 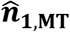 and 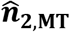, that account for positive and negative NP transverse motions, respectively (Fig. S4.1). In this paper, we have defined right-handed transverse motion as positive and left-handed transverse motion as negative (see Fig. 2C in the main text for the definition of spatial orientation).

**Figure S4.1.**
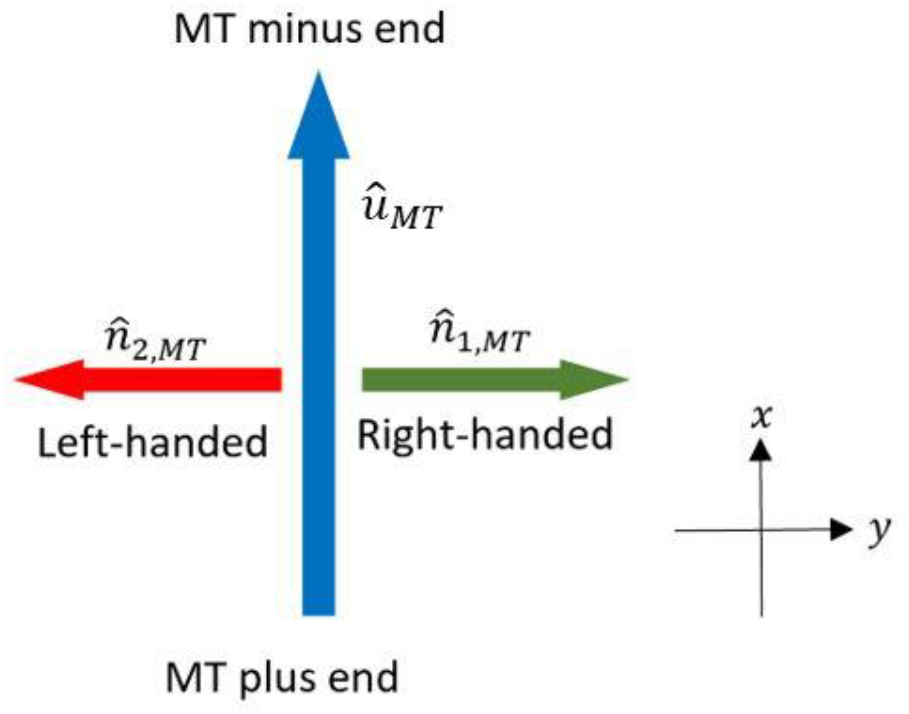
Illustration of the 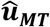 vector (blue arrows) and of the two 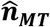 vectors (green and red arrows). The *x*-axis is along the MT cylindrical axis of symmetry, and *y*-axis is orthogonal to it.

Then, we filter unrealistic velocity values as follows: first, we filter out absolute velocity values, 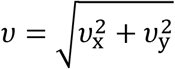, exceeding 4000 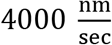, which represents to the very rare – and somewhat not plausible – case where the dynein steps 100 consecutive 40 nm sized steps. Second, before calculating the angular velocity and helical pitch, we filter out unrealistic transverse steps. For that, we set a maximum size of a transverse step, Δ*y* = 2 × *R*_orb_ = 2 × 153 nm (see below), which corresponds to the scenario where the NP moves half a circle along the MT perimeter, from left to right, or *vice versa*. A transverse step that is larger than the said limit is omitted.

We use the mean value of transverse steps, 〈Δ*y*〉, to extract the mean angular velocity, 〈*ω*〉, and its error, *ω*_e_, as follows:

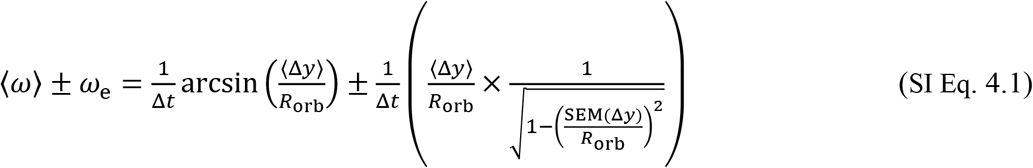

Where SEM(Δ*y*) is the standard error of the mean (SEM) value of Δ*y*, and *R*_orb_ is the approximated distance between the MT and NP center (Figs. 2C and 6A, main text) which reads:

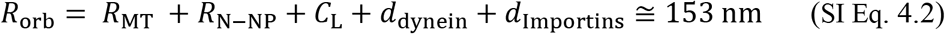

Where *R*_MT_ = 12.5 nm is the MT radius, *C*_L_ = 40.8 nm is the PEG polymer contour length (see section SI.2), *d*_dynein_ = 45nm*^3^ is the dynein characteristic dimension, and *d*_Importins_ = 15nm**^4^ is the *α* and *β* importin complex dimension. *R*_N-NP_ = 40 nm corresponds to the largest Neutravidin-coated NP radius in the sample. The NPs diameter has been extracted from cryo-TEM images (data not shown). This sets the upper limit of accessible *R*_orb_.

Next, we calculate the mean helical pitch, 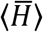, and its corresponding error, *H*_e_, by

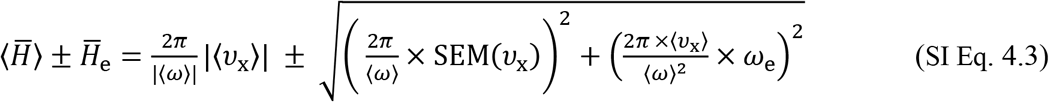

where SEM(*ν*_x_) is the standard error of the mean value of *ν*_x_. Similarly, we can estimate the mean angular velocities for right- (〈*ω*〉|*ω* > 0) and left- (〈*ω*〉|*ω* < 0) handed motions, and the resulting mean helical pitches for the anticipated right- 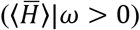 and left- 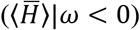 handed helices.

### SI. 5 Experimental mean angular velocity *ω* and helical pitch 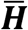

**Table S5.1.**
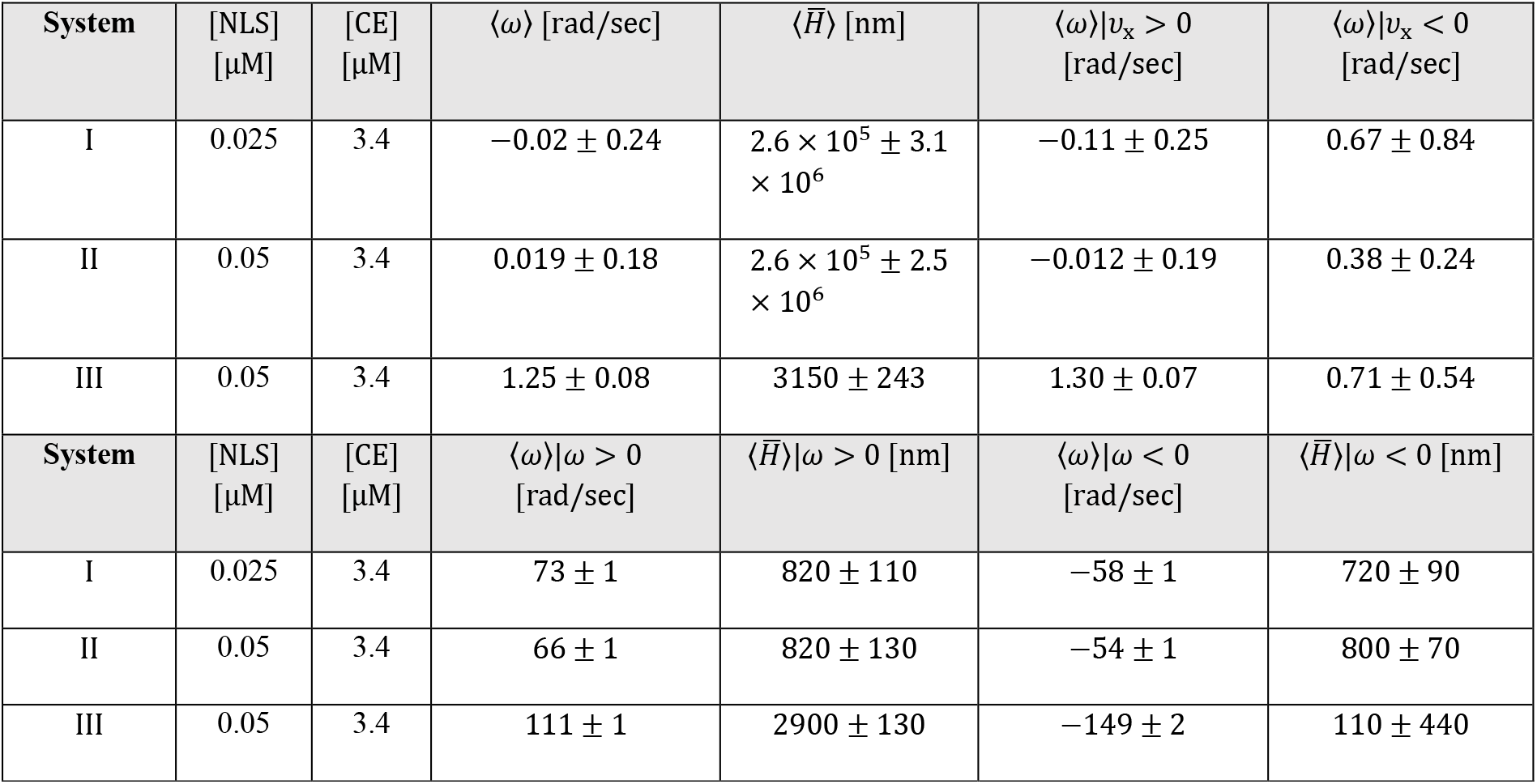
Raw data of Figure 5. Mean angular velocity, 〈*ω*〉, mean angular velocities for right- (〈*ω*〉|*ω* > 0) and left- (〈*ω*〉|*ω* < 0) handed motions, and estimated mean helical pitch size, 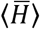, for systems I, II, and III (mean ± SEM). An identical cell extract was used in systems I and II and another extract was used in systems III. The bare NPs mean diameter is 40 nm.

| System | [NLS]<br>[μM] | [CE]<br>[μM] | $\langle\omega\rangle$ [rad/sec] | $\langle\bar{H}\rangle$ [nm] | $\langle\omega\rangle\vert v_x > 0$<br>[rad/sec] | $\langle\omega\rangle\vert v_x < 0$<br>[rad/sec] |
| --- | --- | --- | --- | --- | --- | --- |
| I | 0.025 | 3.4 | $-0.02 \pm 0.24$ | $2.6 \times 10^5 \pm 3.1 \times 10^6$ | $-0.11 \pm 0.25$ | $0.67 \pm 0.84$ |
| II | 0.05 | 3.4 | $0.019 \pm 0.18$ | $2.6 \times 10^5 \pm 2.5 \times 10^6$ | $-0.012 \pm 0.19$ | $0.38 \pm 0.24$ |
| III | 0.05 | 3.4 | $1.25 \pm 0.08$ | $3150 \pm 243$ | $1.30 \pm 0.07$ | $0.71 \pm 0.54$ |
| System | [NLS]<br>[μM] | [CE]<br>[μM] | $\langle\omega\rangle\vert \omega > 0$<br>[rad/sec] | $\langle\bar{H}\rangle\vert \omega > 0$ [nm] | $\langle\omega\rangle\vert \omega < 0$<br>[rad/sec] | $\langle\bar{H}\rangle\vert \omega < 0$ [nm] |
| I | 0.025 | 3.4 | $73 \pm 1$ | $820 \pm 110$ | $-58 \pm 1$ | $720 \pm 90$ |
| II | 0.05 | 3.4 | $66 \pm 1$ | $820 \pm 130$ | $-54 \pm 1$ | $800 \pm 70$ |
| III | 0.05 | 3.4 | $111 \pm 1$ | $2900 \pm 130$ | $-149 \pm 2$ | $110 \pm 440$ |

### SI. 6 Simulation and Algorithm

Computer Simulation Algorithm:

**Figure S6.1.**
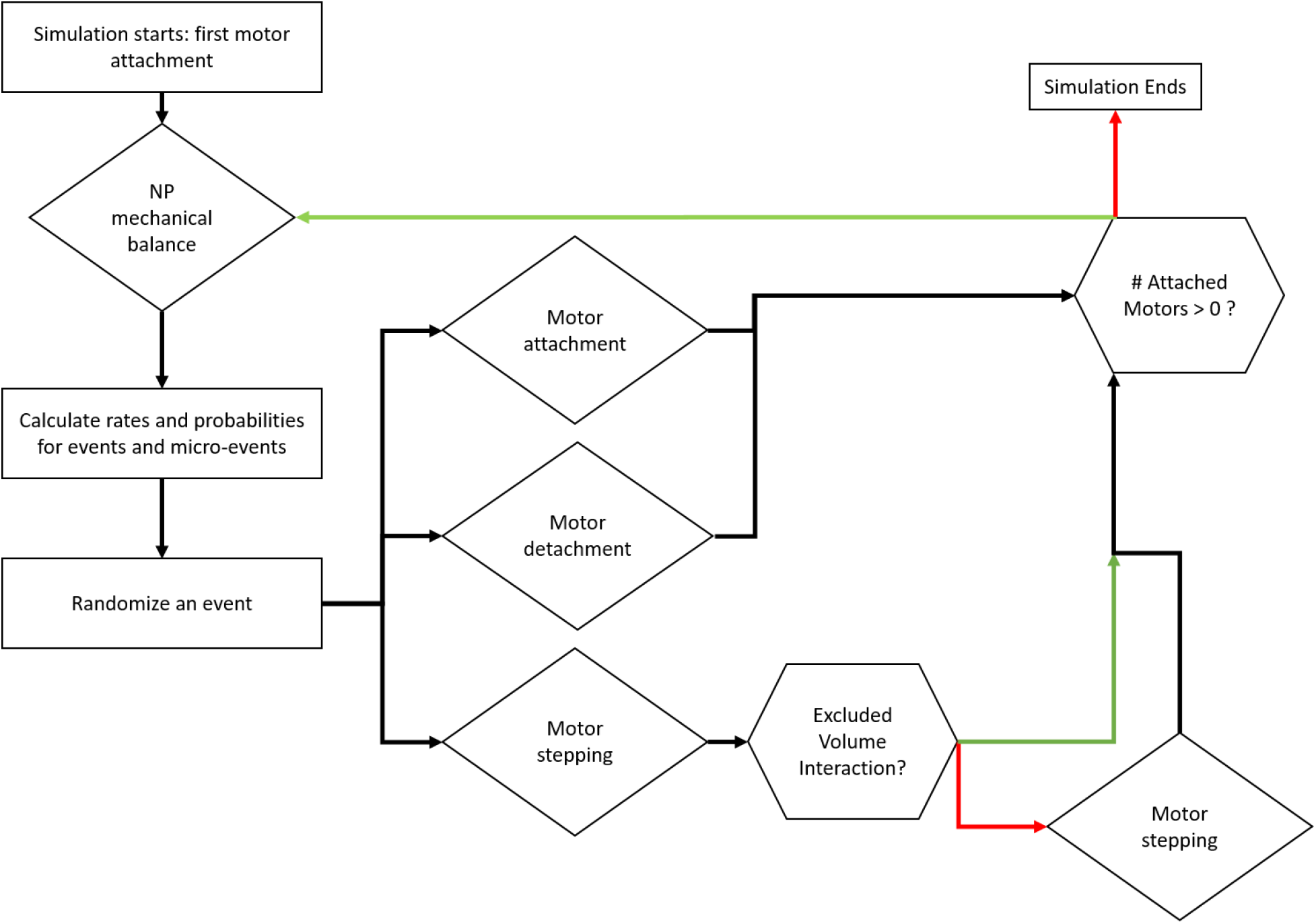
Scheme of the simulation algorithm. Upon a question presented (e.g., “Excluded Volume Interaction?”), green line represents positive answer, and red line represents negative answer.

### SI. 7 Time-scales estimate

There are a few microscopic dynamical processes associated with the NP dynamics that were not accounted for explicitly in our model. As shown below, these processes are much faster than both timescales for motor stepping and motor binding-unbinding kinetics. This separation of timescales allows us to use a few model assumptions mentioned in the fourth chapter. Consider first the Zimm relaxation time for the spacer polymer configurational fluctuations, 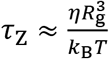, where η is the medium viscosity, and *R*_g_ is the gyration radius. More precisely, for a Gaussian chain^8^, 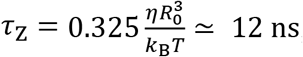, for the value of *R*_0_ ≃ 5.56 nm used in this work (see SI Sec. 2). This is about 5 orders of magnitude shorter than the shortest MC step time in our simulations (*τ*_MC_ ~ 1 ms). Hence, the polymer may be assumed to explore all its configurational phase space during any stepping, binding, or unbinding event (henceforth “MC-event”), which justifies the use of a polymer free-energy without accounting for the polymer dynamics explicitly.

Next, we consider the NP translational and rotational motion. They are involved in the mechanical re-equilibration of the mean NP center-of-mass position and orientation after a MC-event. They are also involved in the relaxation of thermal fluctuations around the mean position when no such events take place. For the translational re-equilibration motion, assume that, after a MC-event, the NP translates a distance r to reach translational mechanical equlibrium. This generates a (polymer) restoring force of magnitude *f* = *K*_p_*r*, where *K*_p_ is the entropic (free-polymer) spring constant, 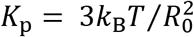. Note that - for large extensions (i.e. when *r* ≫ *R*_0_) – it is much higher, therefore this force can serve as an estimated lower bound. Using the NP translational Stokes drag coefficient, *γ*_t_ = 6*πηR*, we find an (upper bound) estimate for the translational relaxation time, 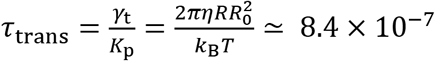 sec (taking the NP radius *R* ≃ 20 nm). This value is about three orders of magnitude shorter than the shortest MC step time in our simulations, *τ*_MC_ ~ 1 ms, thus allowing to assume that the translational equilibration occurs instantaneously after each MC-event.

Next, we wish to estimate the order of magnitude of the NP rotational equilibration time, avoiding exhaustive details associated with the geometry. Assume that, after an MC-event, the NP requires to rotate an angle *θ* to reach rotational mechanical equilibrium. The torque *T* exerted by a force f on the NP is roughly *T* ~ *fR* ≃ *K*_p_*rR*, and the angle *θ* is related to the associated polymer extension by roughly *r* ~ *R θ* leading to a (restoring) torque *T* ~ *K*_p_*R*^2^*θ*. Using the rotational Stokes drag coefficient of the NP, *γ_θ_* = 8*πηR*^3^, leads to the following estimate for the rotational relaxation time, 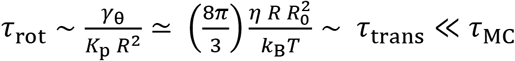, leading to the same conclusion. Linear response theory suggests that the same relaxation times also govern the relaxation of thermal fluctuations near equilibrium, again suggesting that it is adequate to consider the mean center-of-mass position and mean orientation of the NP before and after an MC-event.

### SI. 8 Theoretical raw data for Δ*t* = 0.27 sec

**Table S8.1.** Raw data for Figure 7A. Mean longitudinal velocity, 〈*ν*_x_〉 (mean ± SEM) and STD, 〈*δν*_x_〉, of the different (*R* = 20 nm, *N*_m_) configurations for time intervals equal to 0.27 sec.

| $N_m$ | 1 | 2 | 3 | 4 | 5 | 6 | 7 |
| --- | --- | --- | --- | --- | --- | --- | --- |
| $\langle v_x \rangle$ | $833 \pm 4$ | $769 \pm 3$ | $693 \pm 2$ | $640 \pm 2$ | $605 \pm 1$ | $584 \pm 1$ | $569 \pm 1$ |
| $\langle \delta v_x \rangle$ | 245 | 297 | 322 | 326 | 321 | 315 | 308 |
| $N_m$ | 8 | 9 | 10 | 11 | 12 | 13 | |
| $\langle v_x \rangle$ | $557 \pm 1$ | $547 \pm 1$ | $540 \pm 1$ | $533 \pm 1$ | $528 \pm 1$ | $523 \pm 1$ | |
| $\langle \delta v_x \rangle$ | 302 | 296 | 291 | 286 | 282 | 277 | |

**Table S8.2.**
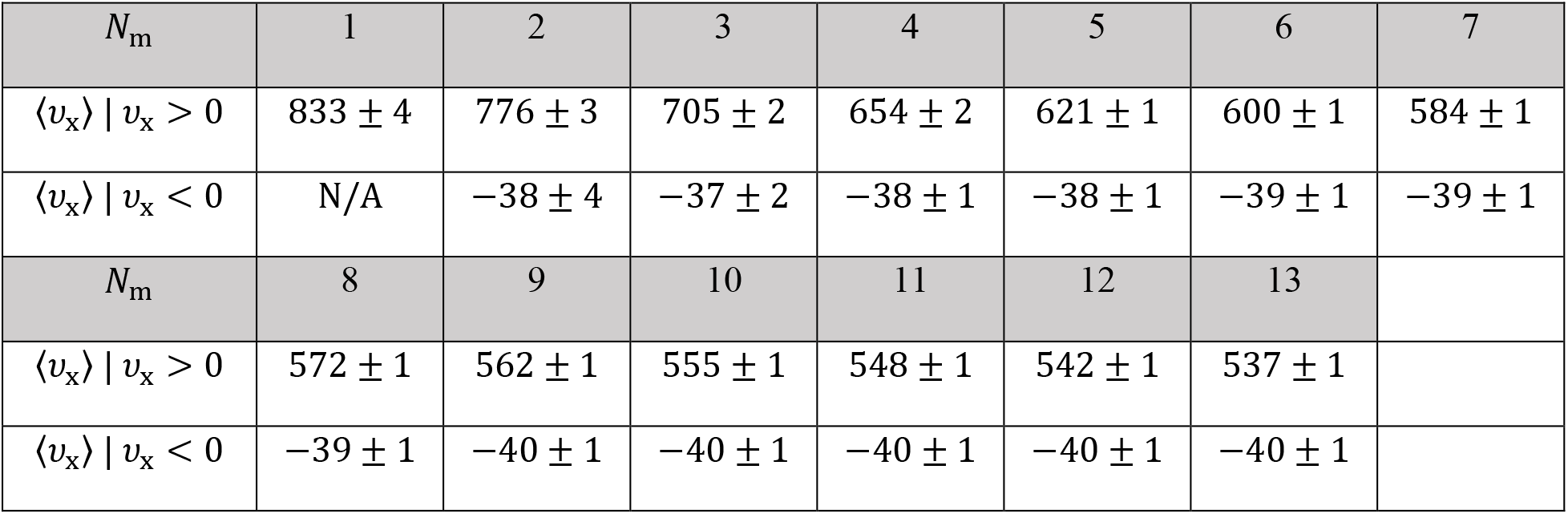
Raw data for Figure 7C. Mean longitudinal velocity 〈*ν*_x_〉 (mean ± SEM) separated for minusend directed, *ν*_x_ > 0, and plus-end directed, *ν*_x_ < 0, directions for the different (*R* = 20 nm, *N*_m_) configurations for time intervals of 0.27 sec.

**Table S8.3.** Raw data for Figure 9A. Mean angular velocity 〈*ω*〉 (mean ± SEM) and STD, 〈*δω*〉, of the different (*R* = 20 nm, *N*_m_) configurations for time intervals equal 0.27 sec.

| $N_m$ | 1 | 2 | 3 | 4 | 5 | 6 | 7 |
| --- | --- | --- | --- | --- | --- | --- | --- |
| $\langle\omega\rangle$ | $1.97 \pm 0.14$ | $1.79 \pm 0.09$ | $1.47 \pm 0.06$ | $1.28 \pm 0.04$ | $1.10 \pm 0.03$ | $0.97 \pm 0.02$ | $0.87 \pm 0.02$ |
| $\langle\delta\omega\rangle$ | 8.16 | 7.88 | 7.28 | 6.78 | 6.37 | 6.06 | 5.80 |
| $N_m$ | 8 | 9 | 10 | 11 | 12 | 13 | |
| $\langle\omega\rangle$ | $0.80 \pm 0.01$ | $0.74 \pm 0.01$ | $0.68 \pm 0.01$ | $0.63 \pm 0.01$ | $0.59 \pm 0.01$ | $0.56 \pm 0.01$ | |
| $\langle\delta\omega\rangle$ | 5.58 | 5.38 | 5.21 | 5.05 | 4.91 | 4.78 | |

**Figure S8.1.**
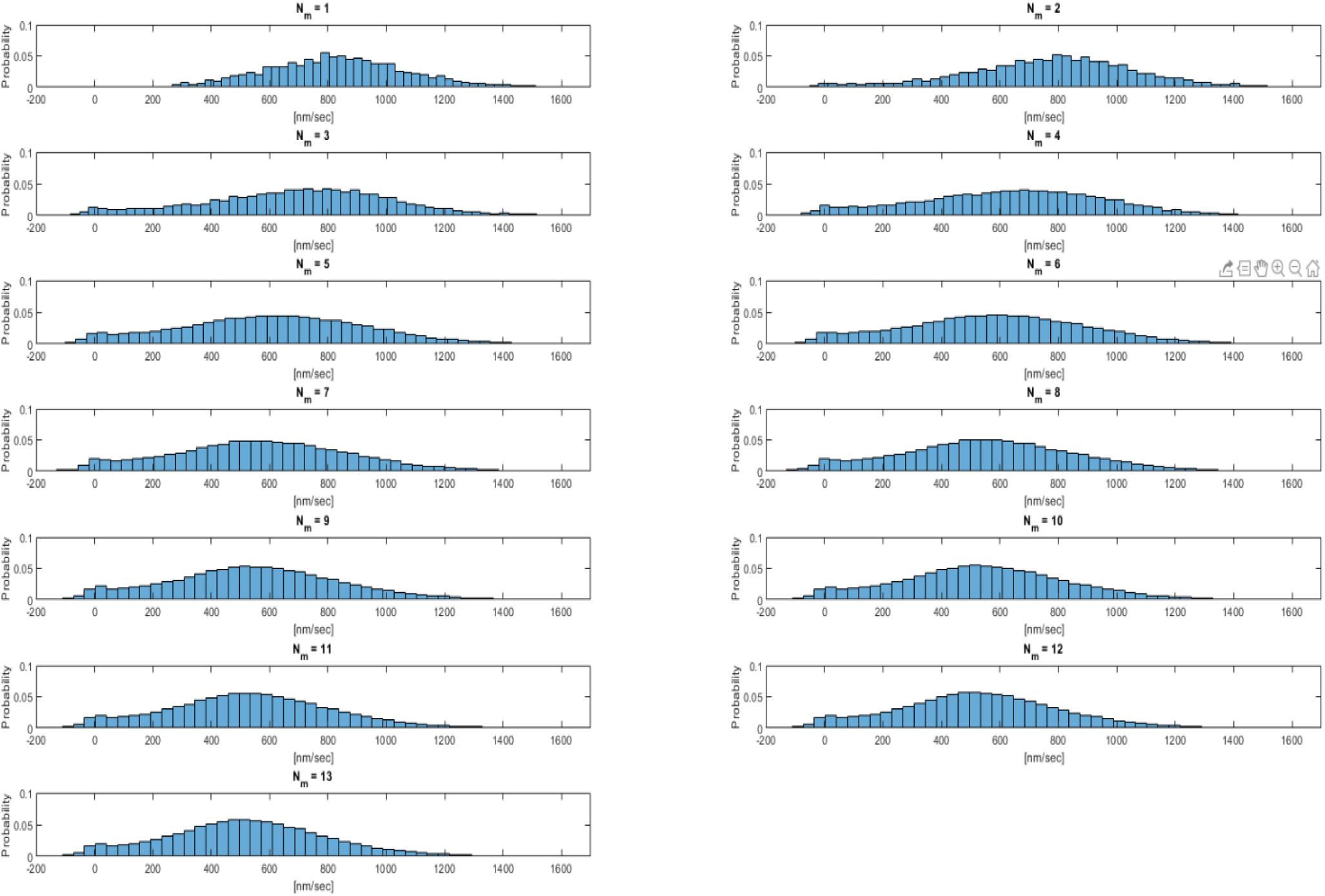
Distributions of the longitudinal velocity, *ν*_x_, for the different (*R* = 20 nm, *N*_m_) configurations for time interval of 0.27 sec.

**Table S8.4.** Raw data for Figure 9C. Mean angular velocity, 〈***ω***〉, separated for minus-end directed, ***ν***_x_ > **0**, and plus-end directed, ***ν***_x_ < **0**, motions for the different (***R*** = ***20 nm, N***_**m**_) configurations for time intervals of **0. 27 sec**. Values correspond to (**mean ± SEM**).

| $N_m$ | 1 | 2 | 3 | 4 | 5 | 6 | 7 |
| --- | --- | --- | --- | --- | --- | --- | --- |
| $\langle \omega \rangle \vert v_x > 0$ | 1.97<br>$\pm 0.14$ | 1.81<br>$\pm 0.09$ | 1.50<br>$\pm 0.06$ | 1.31<br>$\pm 0.04$ | 1.12<br>$\pm 0.03$ | 0.99<br>$\pm 0.02$ | 0.90<br>$\pm 0.02$ |
| $\langle \omega \rangle \vert v_x < 0$ | N/A | 0.05<br>$\pm 0.24$ | 0.05<br>$\pm 0.12$ | 0.05<br>$\pm 0.08$ | 0.04<br>$\pm 0.06$ | 0.06<br>$\pm 0.04$ | 0.07<br>$\pm 0.03$ |
| $N_m$ | 8 | 9 | 10 | 11 | 12 | 13 | |
| $\langle \omega \rangle \vert v_x > 0$ | 0.82<br>$\pm 0.01$ | 0.75<br>$\pm 0.01$ | 0.69<br>$\pm 0.01$ | 0.65<br>$\pm 0.01$ | 0.61<br>$\pm 0.01$ | 0.57<br>$\pm 0.01$ | |
| $\langle \omega \rangle \vert v_x < 0$ | 0.07<br>$\pm 0.03$ | 0.06<br>$\pm 0.02$ | 0.06<br>$\pm 0.02$ | 0.06<br>$\pm 0.02$ | 0.06<br>$\pm 0.02$ | 0.06<br>$\pm 0.01$ | |

### SI. 9 Theoretical raw data for Δ*t* = MC time step

**Figure S9.1.**
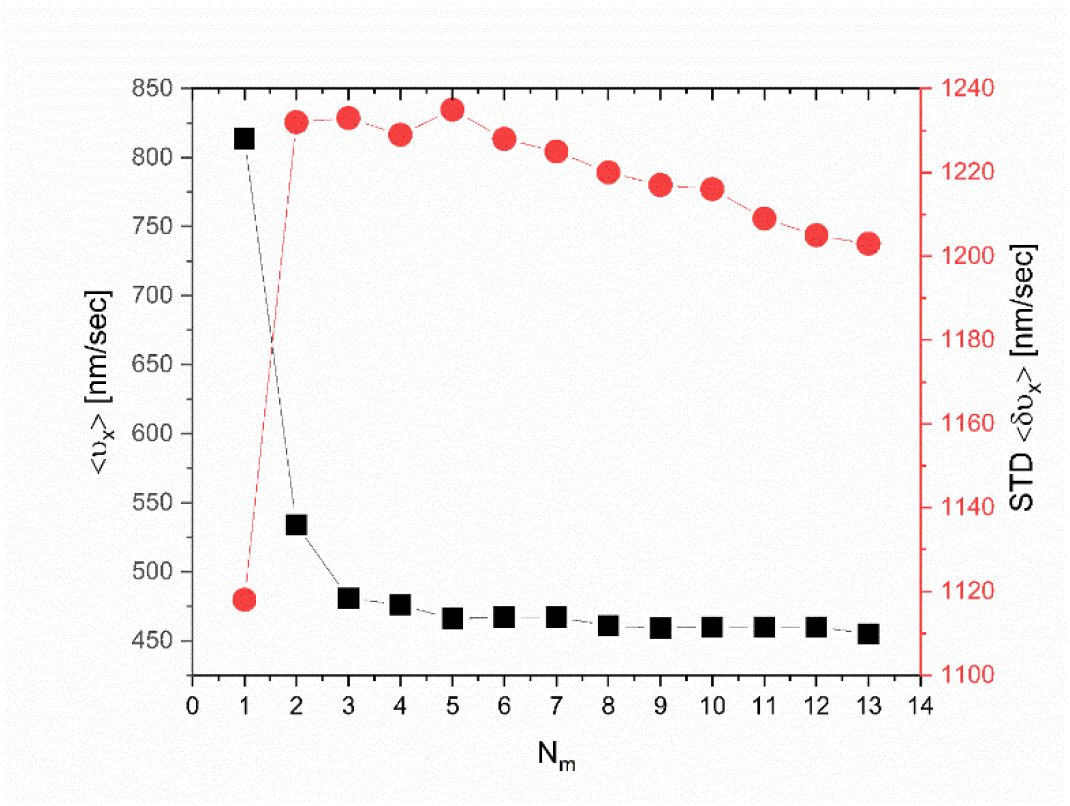
Mean longitudinal velocity, 〈*ν*_x_〉 (mean ± SEM), and STD, 〈*δν*_x_〉, of the different (*R* = 20 nm, *N*_m_) configurations for time intervals equal to MC step times.

**Table S9.1.** Raw data for Figure S9.1. Mean longitudinal velocity, 〈*ν*_x_〉 (mean ± SEM) and STD, 〈*δν*_x_〉, of the different (*R* = 20 nm, *N*_m_) configurations for time intervals equal to MC step times.

| $N_m$ | 1 | 2 | 3 | 4 | 5 | 6 | 7 |
| --- | --- | --- | --- | --- | --- | --- | --- |
| $\langle v_x \rangle$ | $814 \pm 4$ | $534 \pm 4$ | $481 \pm 2$ | $458 \pm 2$ | $466 \pm 2$ | $467 \pm 2$ | $467 \pm 2$ |
| $\langle \delta v_x \rangle$ | 1188 | 1232 | 1233 | 1227 | 1235 | 1228 | 1225 |
| $N_m$ | 8 | 9 | 10 | 11 | 12 | 13 | |
| $\langle v_x \rangle$ | $461 \pm 1$ | $459 \pm 1$ | $460 \pm 1$ | $454 \pm 2$ | $451 \pm 1$ | $452 \pm 1$ | |
| $\langle \delta v_x \rangle$ | 1220 | 1217 | 1216 | 1208 | 1203 | 1202 | |

**Figure S9.2.**
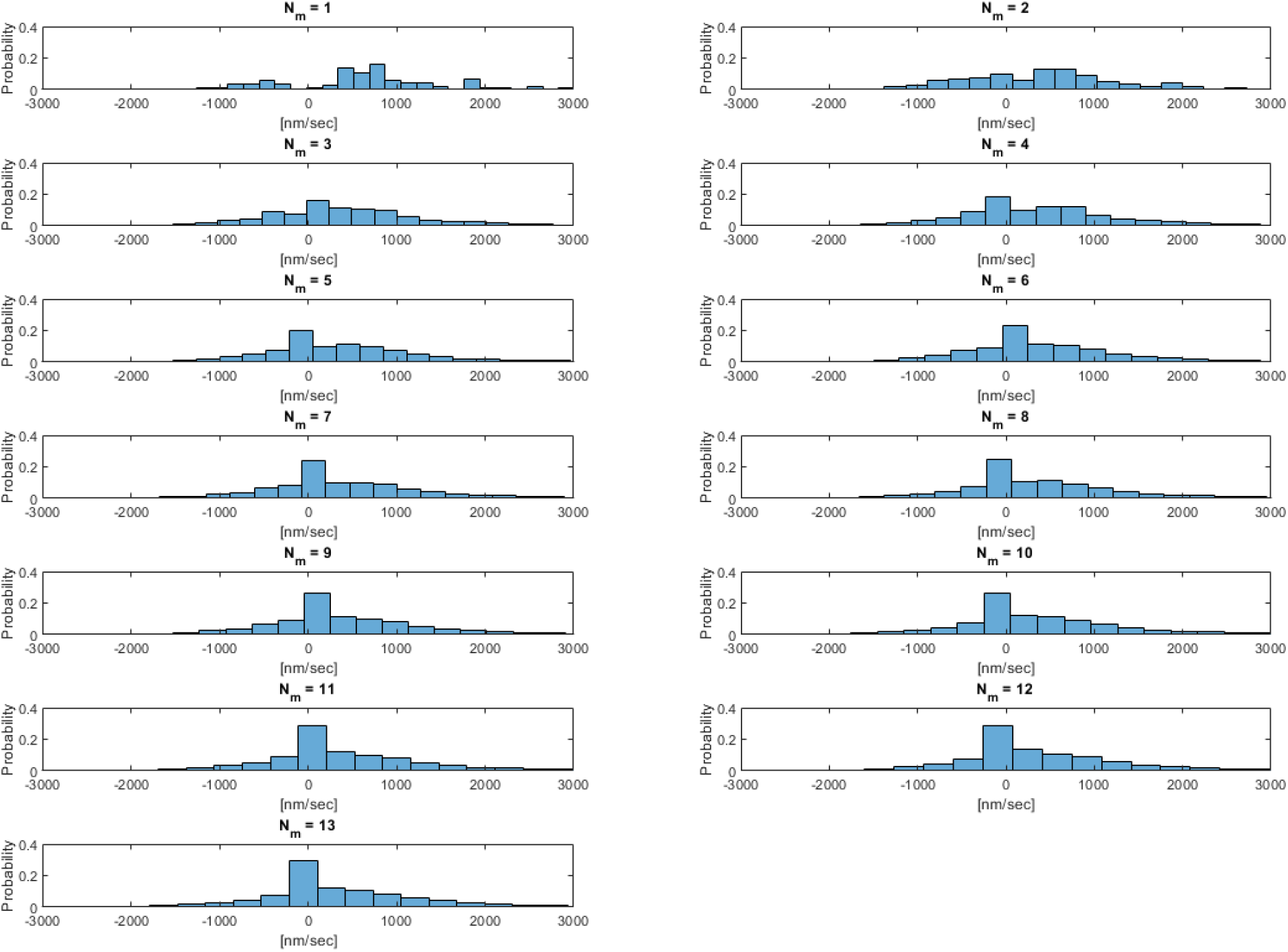
Distributions of the longitudinal velocity, *ν*_x_, for the different (*R* = 20 nm, *N*_m_) configurations for time intervals equal to MC step times.

**Table S9.2.**
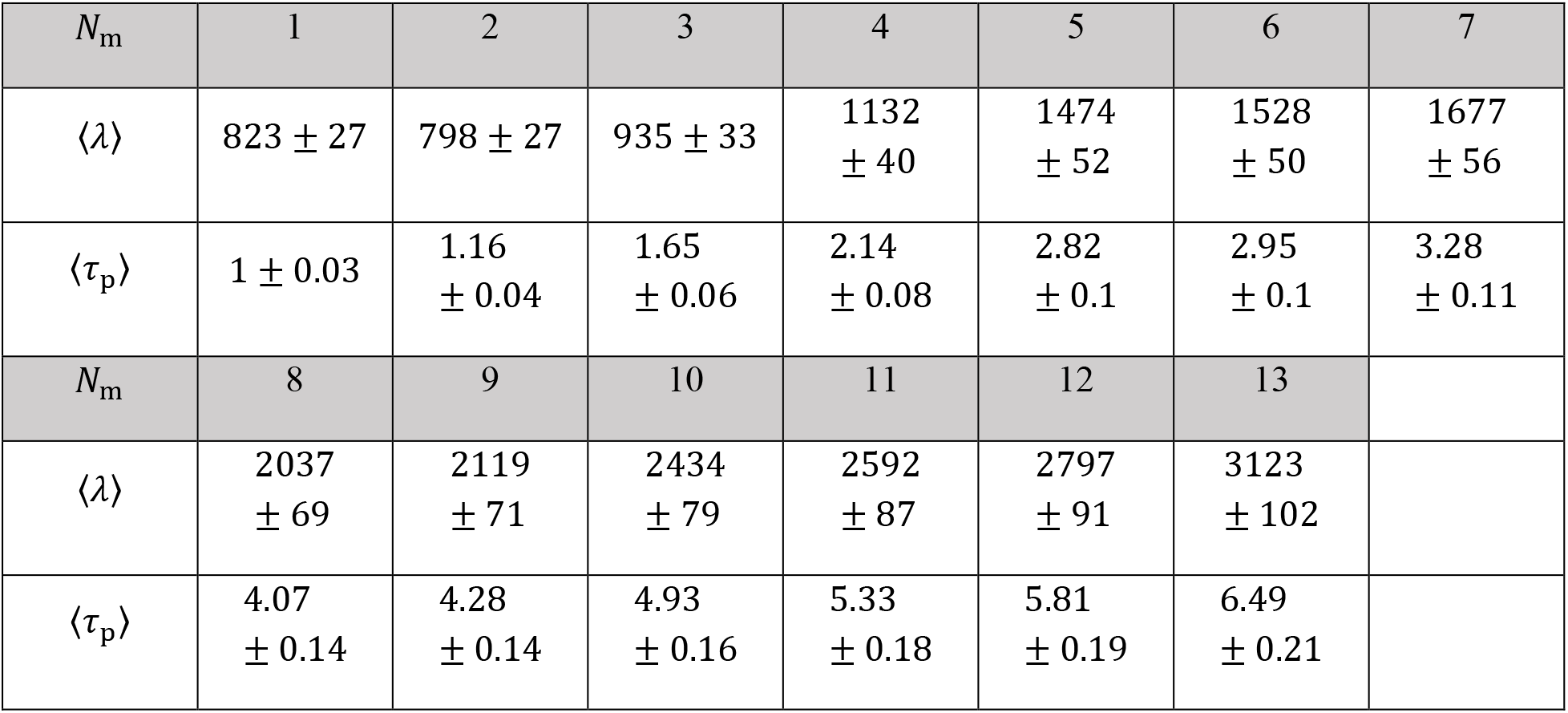
Raw data for Figure 7B. Mean longitudinal run-lengths (along the MT symmetry axis), 〈*λ*〉, and processivity times, 〈*τ*_p_〉, for the different (*R* = 20 nm, *N*_m_) configurations for time intervals equal to MC step times. Values correspond to (mean ± SEM).

| $N_m$ | 1 | 2 | 3 | 4 | 5 | 6 | 7 |
| --- | --- | --- | --- | --- | --- | --- | --- |
| $\langle\lambda\rangle$ | $823 \pm 27$ | $798 \pm 27$ | $935 \pm 33$ | $1132 \pm 40$ | $1474 \pm 52$ | $1528 \pm 50$ | $1677 \pm 56$ |
| $\langle\tau_p\rangle$ | $1 \pm 0.03$ | $1.16 \pm 0.04$ | $1.65 \pm 0.06$ | $2.14 \pm 0.08$ | $2.82 \pm 0.1$ | $2.95 \pm 0.1$ | $3.28 \pm 0.11$ |
| $N_m$ | 8 | 9 | 10 | 11 | 12 | 13 | |
| $\langle\lambda\rangle$ | $2037 \pm 69$ | $2119 \pm 71$ | $2434 \pm 79$ | $2592 \pm 87$ | $2797 \pm 91$ | $3123 \pm 102$ | |
| $\langle\tau_p\rangle$ | $4.07 \pm 0.14$ | $4.28 \pm 0.14$ | $4.93 \pm 0.16$ | $5.33 \pm 0.18$ | $5.81 \pm 0.19$ | $6.49 \pm 0.21$ | |

**Table S9.3.**
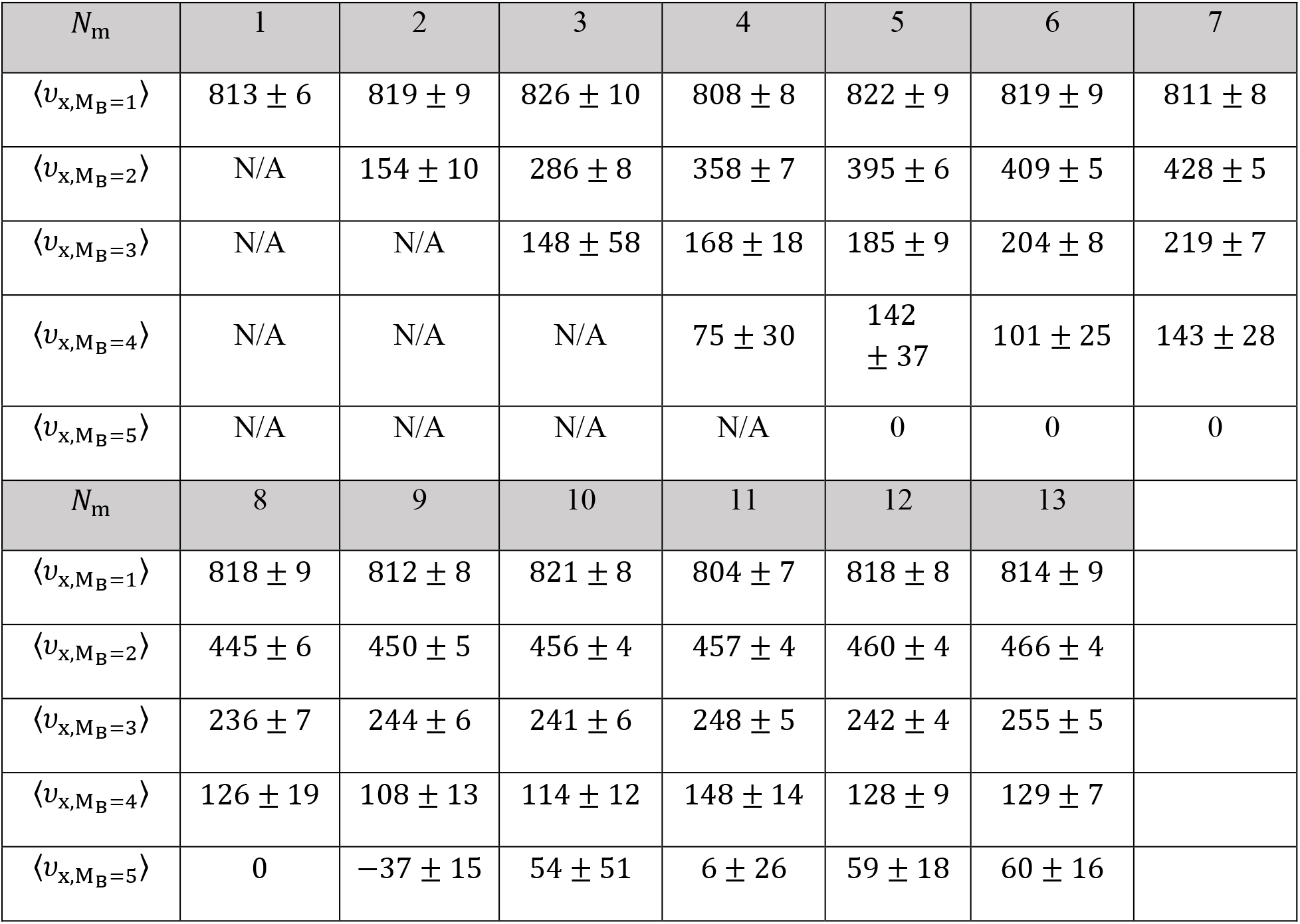
Raw data for Figure 8C. Mean longitudinal velocity of the different NP states, where a state is defined according to the number of MT bound motors, *M*_B_, for the different (*R* = 20 nm, *N*_m_) configurations for time intervals equal to MC step times (mean ± SEM).

**Table S9.4.** Longitudinal velocity STD of the different NP states. A state is defined according to the number of MT bound motors, MB, for the different (*R* = 20 nm, *N*_m_) configurations for time intervals equal to MC step times (see Fig. 8C in main text).

| $N_M$ | 1 | 2 | 3 | 4 | 5 | 6 | 7 |
| --- | --- | --- | --- | --- | --- | --- | --- |
| $\langle \delta v_{x, M_B=1} \rangle$ | 0 | 296 | 321 | 251 | 284 | 252 | 256 |
| $\langle \delta v_{x, M_B=2} \rangle$ | N/A | 163 | 227 | 205 | 188 | 169 | 158 |
| $\langle \delta v_{x, M_B=3} \rangle$ | N/A | N/A | 487 | 239 | 206 | 211 | 201 |
| $\langle \delta v_{x, M_B=4} \rangle$ | N/A | N/A | N/A | 60 | 150 | 157 | 255 |
| $\langle \delta v_{x, M_B=5} \rangle$ | N/A | N/A | N/A | N/A | N/A | N/A | N/A |

| $N_M$ | 8 | 9 | 10 | 11 | 12 | 13 |
| --- | --- | --- | --- | --- | --- | --- |
| $\langle \delta v_{x, M_B=1} \rangle$ | 285 | 247 | 254 | 234 | 256 | 293 |
| $\langle \delta v_{x, M_B=2} \rangle$ | 178 | 144 | 121 | 123 | 120 | 121 |
| $\langle \delta v_{x, M_B=3} \rangle$ | 214 | 179 | 188 | 145 | 123 | 157 |
| $\langle \delta v_{x, M_B=4} \rangle$ | 217 | 198 | 226 | 189 | 177 | 169 |
| $\langle \delta v_{x, M_B=5} \rangle$ | N/A | N/A | 141 | 58 | 80 | 96 |

**Figure S9.3.**
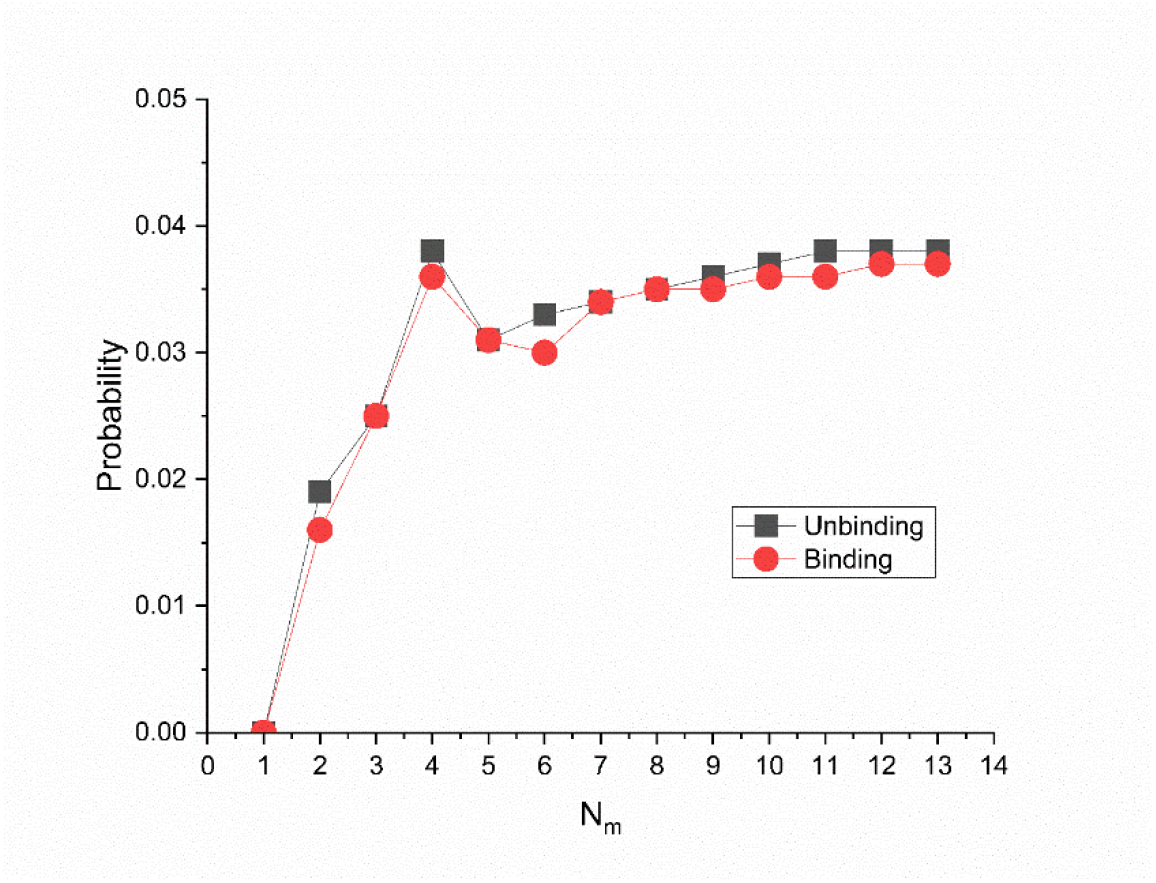
Fraction of binding and unbinding events out of the total possible events for the different (*R* = 20 nm, *N*_m_) configurations for time intervals equal to MC step times.

**Table S9.5.** Raw data for Figure S9.3. Fraction of binding and unbinding events out of the total possible events for the different (*R* = 20 nm, *N*_m_) configurations for time intervals equal to MC step times.

| $N_m$ | 1 | 2 | 3 | 4 | 5 | 6 | 7 |
| --- | --- | --- | --- | --- | --- | --- | --- |
| $F_{\text{unbinding}}$ | <0.01 | 0.019 | 0.025 | 0.032 | 0.031 | 0.033 | 0.034 |
| $F_{\text{binding}}$ | <0.01 | 0.016 | 0.025 | 0.029 | 0.031 | 0.03 | 0.034 |
| $N_m$ | 8 | 9 | 10 | 11 | 12 | 13 | |
| $F_{\text{unbinding}}$ | 0.035 | 0.036 | 0.037 | 0.038 | 0.038 | 0.038 | |
| $F_{\text{binding}}$ | 0.035 | 0.035 | 0.036 | 0.036 | 0.037 | 0.037 | |

**Figure S9.4.**
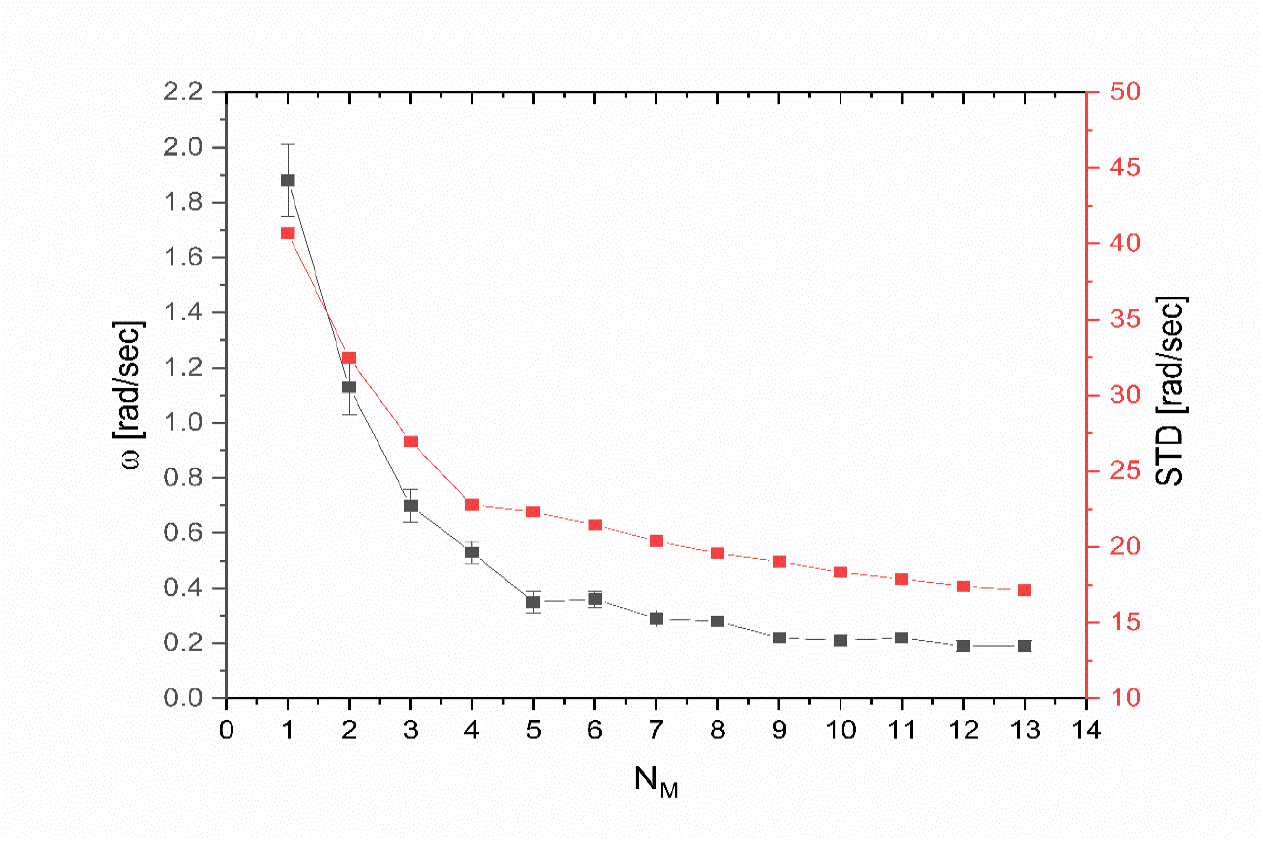
Mean angular velocity, 〈*ω*〉 (mean ± SEM), and STD, 〈*δω*〉, of the different (*R* = 20 nm, *N*_m_) configurations for time intervals equal to MC step times.

**Table S9.6.**
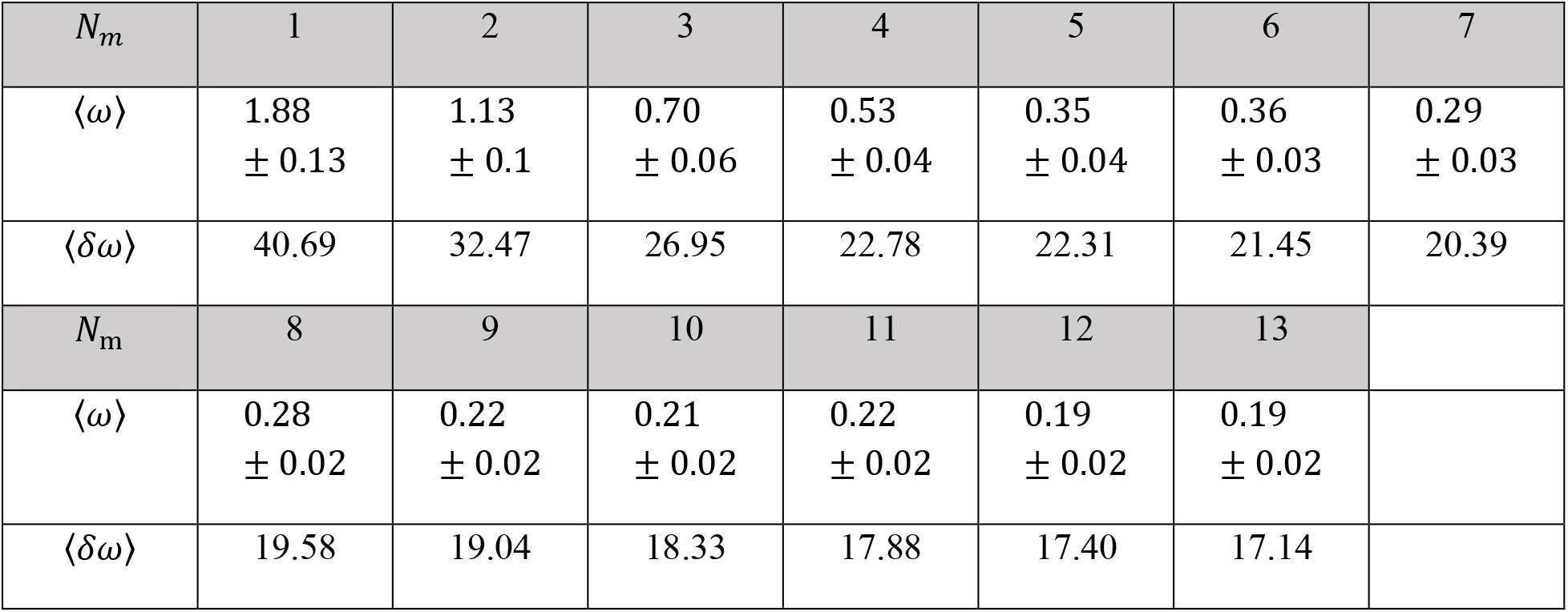
Raw data for Figure S9.4. Mean angular velocity, 〈*ω*〉 (mean ± SEM), and STD, 〈*δω*〉, of the different (*R* = 20 nm, *N*_m_) configurations for time intervals equal to MC step times.

| $N_m$ | 1 | 2 | 3 | 4 | 5 | 6 | 7 |
| --- | --- | --- | --- | --- | --- | --- | --- |
| $\langle \omega \rangle$ | 1.88<br>$\pm 0.13$ | 1.13<br>$\pm 0.1$ | 0.70<br>$\pm 0.06$ | 0.53<br>$\pm 0.04$ | 0.35<br>$\pm 0.04$ | 0.36<br>$\pm 0.03$ | 0.29<br>$\pm 0.03$ |
| $\langle \delta \omega \rangle$ | 40.69 | 32.47 | 26.95 | 22.78 | 22.31 | 21.45 | 20.39 |
| $N_m$ | 8 | 9 | 10 | 11 | 12 | 13 | |
| $\langle \omega \rangle$ | 0.28<br>$\pm 0.02$ | 0.22<br>$\pm 0.02$ | 0.21<br>$\pm 0.02$ | 0.22<br>$\pm 0.02$ | 0.19<br>$\pm 0.02$ | 0.19<br>$\pm 0.02$ | |
| $\langle \delta \omega \rangle$ | 19.58 | 19.04 | 18.33 | 17.88 | 17.40 | 17.14 | |

**Figure S9.5.**
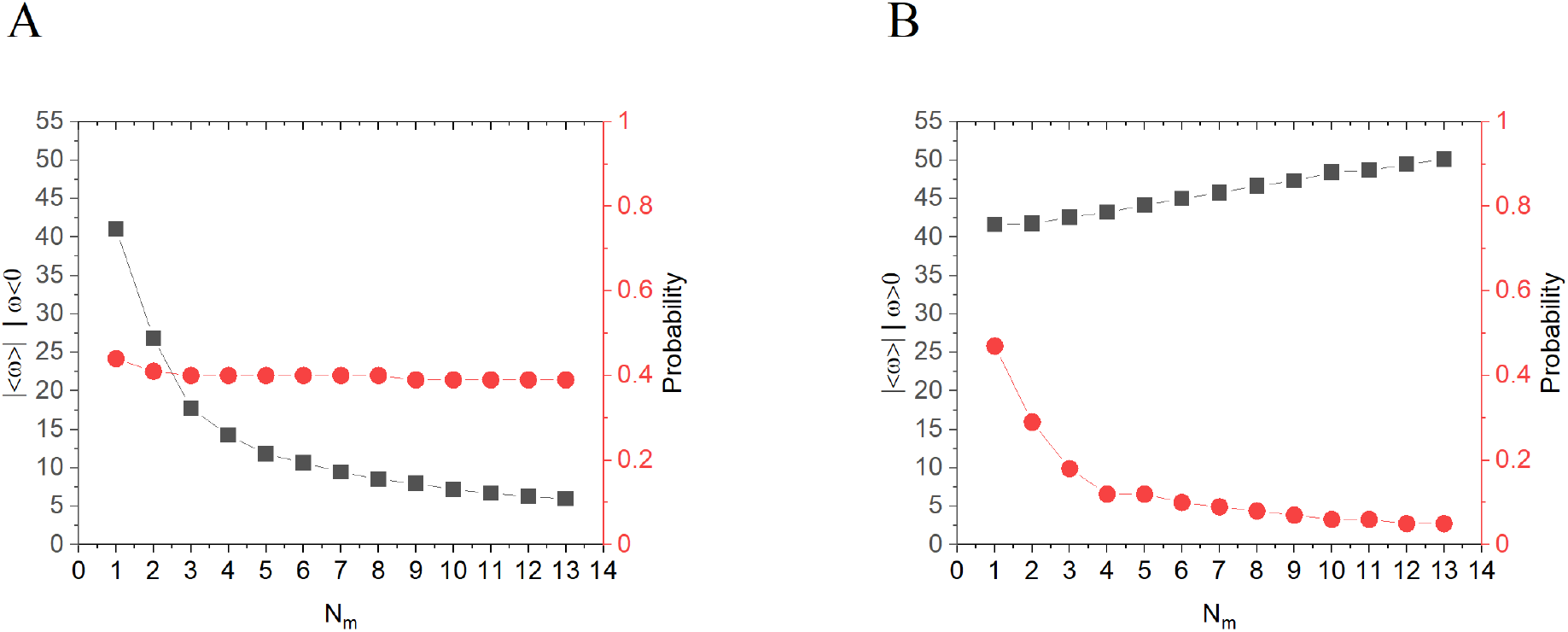
Mean absolute angular velocity segregated to left-handed and right-handed components of the distin*ct* (*R* = 20 nm, *N*_m_) configurations for time intervals equal to MC step times. **(A)** Mean absolute left-handed angular velocity, |〈*ω*_L_〉| = |〈*ω*〉| | *ω* < 0, (black) and probability for left-handed motion (red). **(B**) Mean absolute right-handed angular velocity, |〈*ω*_R_〉| = ļ〈*ω*〉| | *ω* > 0, (black) and probability for right-handed motion (red).

**Table S9.7.** Raw data for Figure S9.5. Mean angular velocity separated to left- (〈*ω*_L_〉) and right- (〈*ω*_R_〉) handed components of the distinct (*R* = 20 nm, *N*_m_) configurations for time intervals equal to MC step times. Values correspond to (mean ± SEM).

| $N_m$ | 1 | 2 | 3 | 4 | 5 | 6 | 7 |
| --- | --- | --- | --- | --- | --- | --- | --- |
| $\langle\omega_L\rangle$ | $-41.07 \pm 0.05$ | $-26.86 \pm 0.1$ | $-17.74 \pm 0.07$ | $-14.24 \pm 0.06$ | $-11.83 \pm 0.05$ | $-10.67 \pm 0.05$ | $-9.46 \pm 0.04$ |
| $\langle\omega_R\rangle$ | $41.66 \pm 0.05$ | $41.76 \pm 0.08$ | $42.59 \pm 0.07$ | $43.25 \pm 0.07$ | $44.18 \pm 0.07$ | $45.02 \pm 0.08$ | $45.79 \pm 0.08$ |
| $N_m$ | 8 | 9 | 10 | 11 | 12 | 13 | |
| $\langle\omega_L\rangle$ | $-8.52 \pm 0.04$ | $-7.97 \pm 0.04$ | $-7.18 \pm 0.03$ | $-6.71 \pm 0.03$ | $-6.29 \pm 0.03$ | $-5.99 \pm 0.03$ | |
| $\langle\omega_R\rangle$ | $46.7 \pm 0.08$ | $47.34 \pm 0.09$ | $48.44 \pm 0.09$ | $48.70 \pm 0.09$ | $49.50 \pm 0.09$ | $50.15 \pm 0.09$ | |

**Table S9.8.**
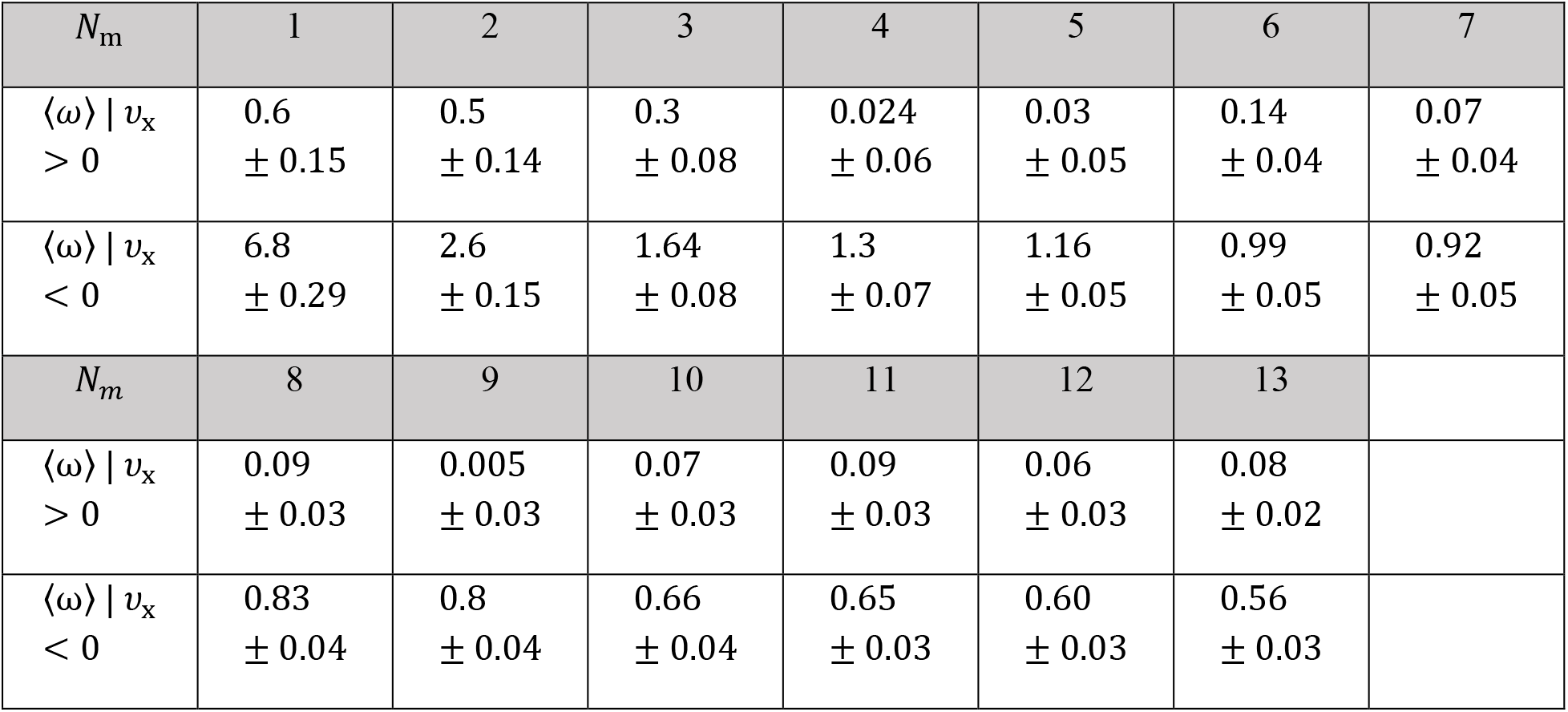
Raw data for Figure 9B. Absolute mean angular velocity segregated to minus-end directed, 〈*ω*〉 | *ν*_x_ > 0, and plus-end directed, 〈*ω*〉 | *ν*_x_ < 0, motions for the distinct (*R* = 20 nm, *N*_m_) configurations for time intervals equal to MC step times (mean ± SEM).

**Table S9.9.** Raw data for Figure 10A. Analysis of the actual NP helical motion from observed (simulated) trajectories separated to left- and right-handed motions (mean ± SEM). 〈*H*_L_〉 is the mean helical pitch size for left-handed helices (i.e., *ϕ* = −2*π*), and 〈*H_R_*〉 is the mean helical pitch size for right-handed helices (i.e., *ϕ* = +2*π*).

| $N_m$ | 1 | 2 | 3 | 4 | 5 | 6 | 7 |
| --- | --- | --- | --- | --- | --- | --- | --- |
| $\langle H_L \rangle$ | 993 ± 97 | 857 ± 79 | 1201<br>± 106 | 1185<br>± 72 | 1383<br>± 96 | 1794<br>± 133 | 1746<br>± 173 |
| $\langle H_R \rangle$ | 789 ± 28 | 824 ± 44 | 992 ± 42 | 1173<br>± 55 | 1542<br>± 65 | 1565<br>± 70 | 1671<br>± 71 |
| $N_m$ | 8 | 9 | 10 | 11 | 12 | 13 | |
| $\langle H_L \rangle$ | 2173<br>± 169 | 2034<br>± 183 | 2383<br>± 239 | 2541<br>± 191 | 2825<br>± 237 | 3169<br>± 245 | |
| $\langle H_R \rangle$ | 2026<br>± 92 | 2151<br>± 89 | 2575<br>± 120 | 2602<br>± 119 | 2780<br>± 133 | 3068<br>± 134 | |

**Table S9.10.**
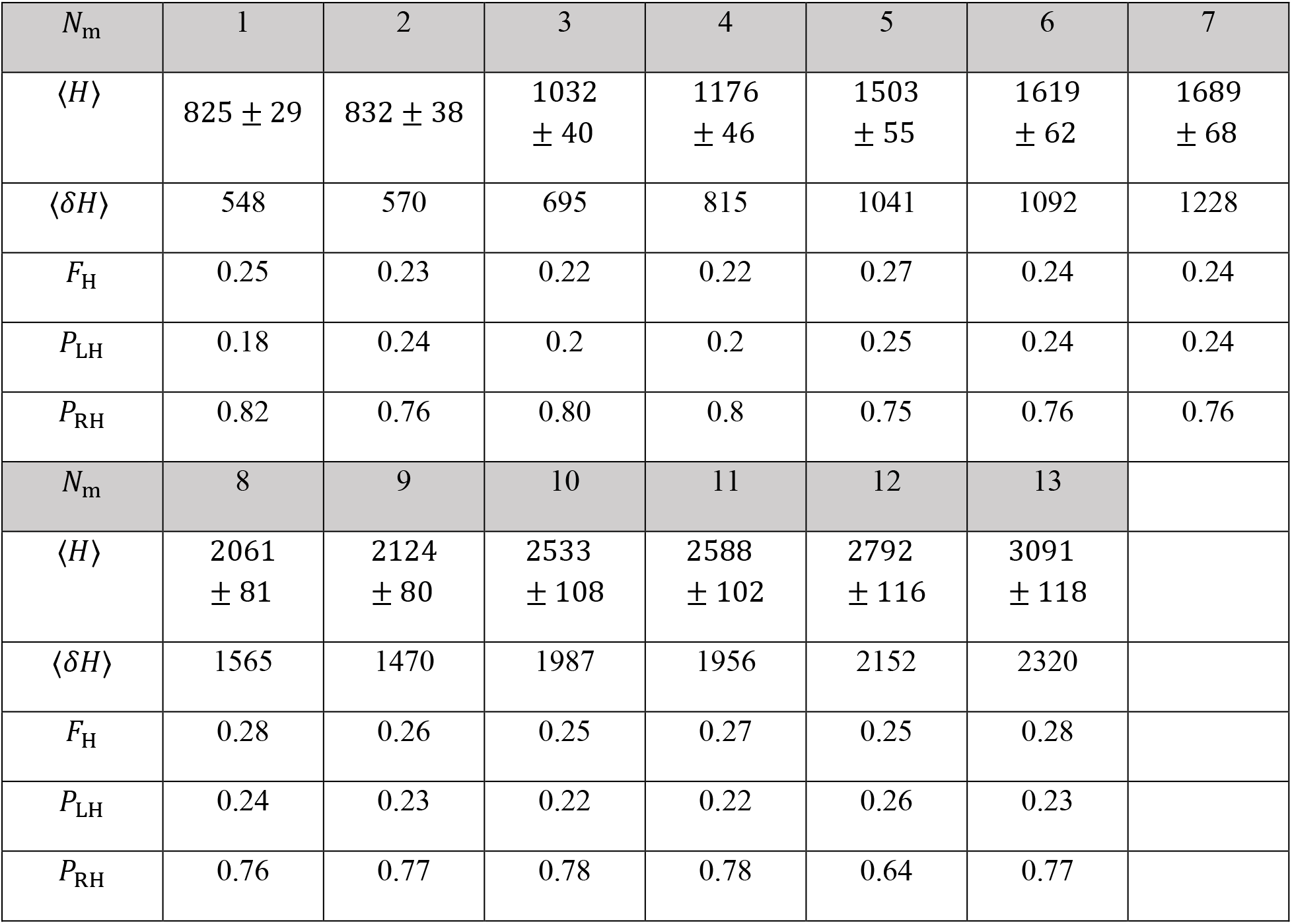
Raw data Figure 10B. Analysis of the actual NP helical motion from observed (simulated) trajectories, where 〈*H*〉 is the mean helical pitch size (mean ± SEM), 〈*δH*〉 is the helical pitch STD, *F*_H_ is the fraction of NP trajectories that completed at least one wind around the MT symmetry axis out of the total number of trajectories, *P*_LH_ is the probability for a left-handed helical motion, and *P*_RH_ is the probability for a right-handed helical motion. The mean pitch size is calculated by averaging over all the helical pitches of the same (*R, N*_m_) configuration, regardless of their direction (i.e., *ϕ* = +2*π* or i.e., *φ* = −2*π*). A helical pitch size equals to the NP accumulated longitudinal motion – towards the MT minus-end – up to the point when a wind around the MT is completed (i.e., *ϕ* = ±2*π*). Note that the motion does not have to be consistent towards either left-handed or right-handed direction, e.g., a right-handed helix can contain left-handed motion (Δ*ϕ* < 0) and vice versa, or in other words, the large transverse fluctuations do not reset the *ϕ* = ∑Δ*ϕ* count towards a wind completion (±2*π*).

**Table S9.11.**
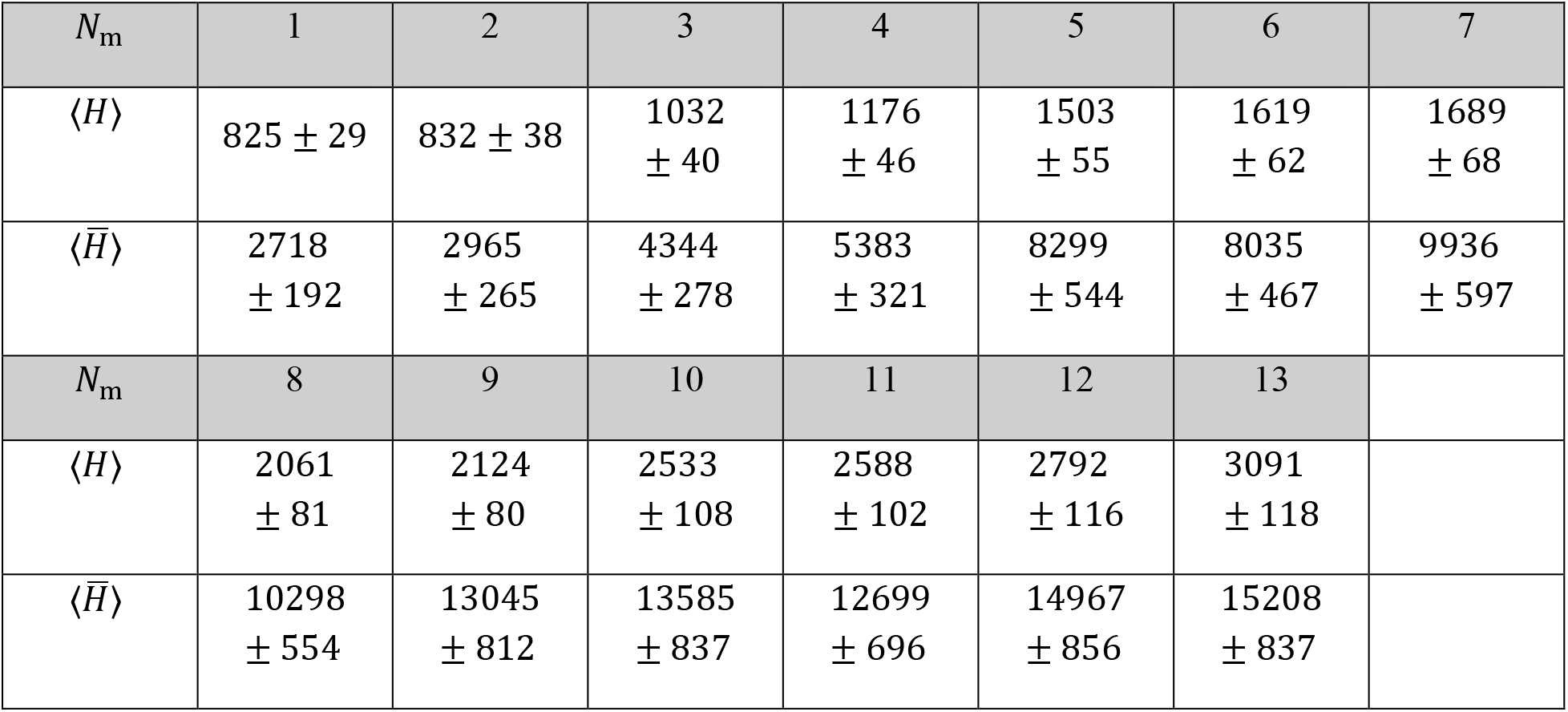
Raw data for Figure 10C. Helical pitch size estimated from the angular and longitudinal mean velocity values, 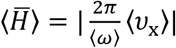. Values correspond to (mean ± SEM).

### SI. 10 Binomial Distribution of NP Bound Motors

The number of motors *N*_m_ that are bound to the NP (not to be confused with the MT-bound motors) is a random variable, as in any adsorption process to a finite size surface, suggesting that this number fluctuates between different NPs. The mean number of motors 〈*N*_m_〉 is related to the (mean) PEG-NLS-*αβ*-dynein anchoring distance *ξ* via 〈*N*_m_〉 = 4*πR*^2^/ξ^2^ or 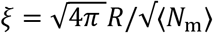. Likewise, we may define a mean surface relative coverage 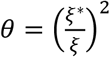 describing the fraction of PEG-NLSs that end with a bound motor. In order to account for the fluctuations, we assume independent site (i.e. PEG-NLS) adsorption with probability θ for adsorption per site, suggesting that the random bound motor number obeys the well-known binomial distribution

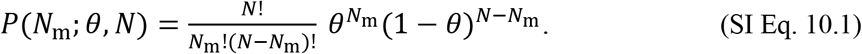

where *N* is the number of PEG-NLSs, consistent with the mean number of motors obeying 〈*N*_m_〉 = *Nθ*.

However, since only NPs with *N*_m_ ≥ 1 can bind to the MT, by assumption, we require the conditional binomial distribution *P*^*^(*N*_m_; 0, *N*) ≡ (*P*(*N*_m_; 0, *N*| *N*_m_ ≥ 1), which is given by

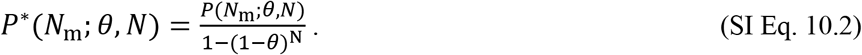

Considering the mean number of motors among those NPs with *N*_m_ ≥ 1 we find

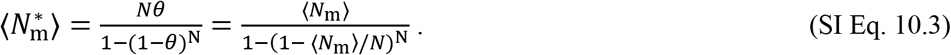

Given a value of *θ*, we can compute any mean of motility variable, *f*(*N*_m_), of the NP ensemble associated with this value of *θ*, e.g., velocity, run-time, and so on, as

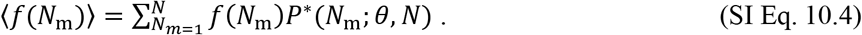

To examine the effect of the conditional binomial distribution on the observed motility, we first consider a characteristic value of 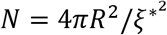 that corresponds to the estimated value of *ξ*^*^ of systems II-III, *N* ≃ 5 (using *ξ*^*^ from Table 1 in the main text). Using this value of *N*, we depict in Fig. S10.1 the theoretical ensemble average motility variables with 0 ranging from 0.1 to 1, which determines 〈*N*_m_〉 = *Nθ*. We do that for results obtained for both MC-step-time and experimental time interval (0.27 sec). We also show (red squares), for comparison, the theoretical (simulation results) for the case of a deterministic *N*_m_. While deviations between the two calculations (obviously) do appear, they are not significant for most cases. The most pronounced differences appear in the MC-step-time longitudinal mean velocity for *N*_m_ = 2,3 (due to the contribution to the mean of the *N*_m_ = 1 population, whose velocity is high), which is strongly reduced for results obtained on 0.27 sec time interval. To check the sensitivity to the value of *N*, we also examine in Fig. S10.2 the case of N = 10 (even though it does not correspond to any of the experimental systems IIV). Similar to Fig. S10.1, pronounced deviations appear in the MC-step-time longitudinal mean velocity for *N*_m_ = 2,3, and 4, but again the deviations are strongly reduced for results obtained for 0.27 sec time interval. We conclude that while accurate comparison between experimental and theoretical results do require the (conditional) binomial distribution averaging, for approximate comparison it is sufficient to consider deterministic values of *N*_m_.

**Figure S10.1.**
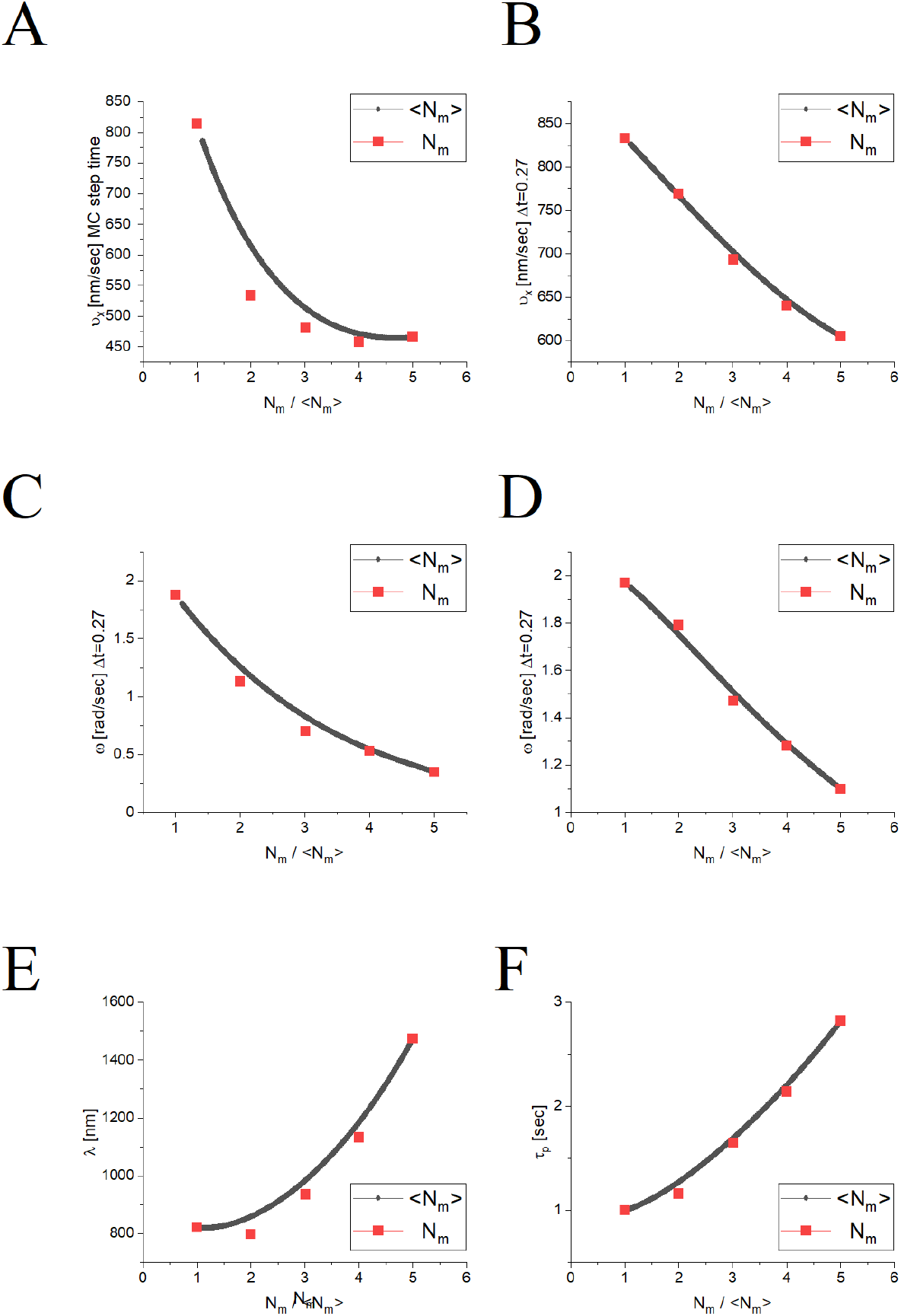
Theoretical comparison between motility variables associated with deterministic values of *N*_m_, and those averaged over the conditional binomial distribution, SI Eq. 10.2, using *N* = 5 (〈*N*_m_〉). Red squares depict the motility variables against a deterministic number of NP-bound motors, *N*_m_, i.e. when all the particles have exactly the same *N*_m_. Black lines depict the motility variables against the mean number of NP-bound motors, 〈*N*_m_〉, i.e. when the value of *N*_m_ can vary between NPs – hence yielding also non-integer numbers 〈*N*_m_〉. We show the following motility variables: **(A)** Longitudinal velocity, *ν*_x_, for MC step time. (**B**) Longitudinal velocity, *ν*_x_, for 0.27 sec time interval. **(C)** Angular velocity, *ω*, for MC step time (**D**) Angular velocity, *ω*, for 0.27 sec time interval. (**E**) Longitudinal run-length, A. (**F**) Runtime, *τ*_p_.

**Figure S10.2.**
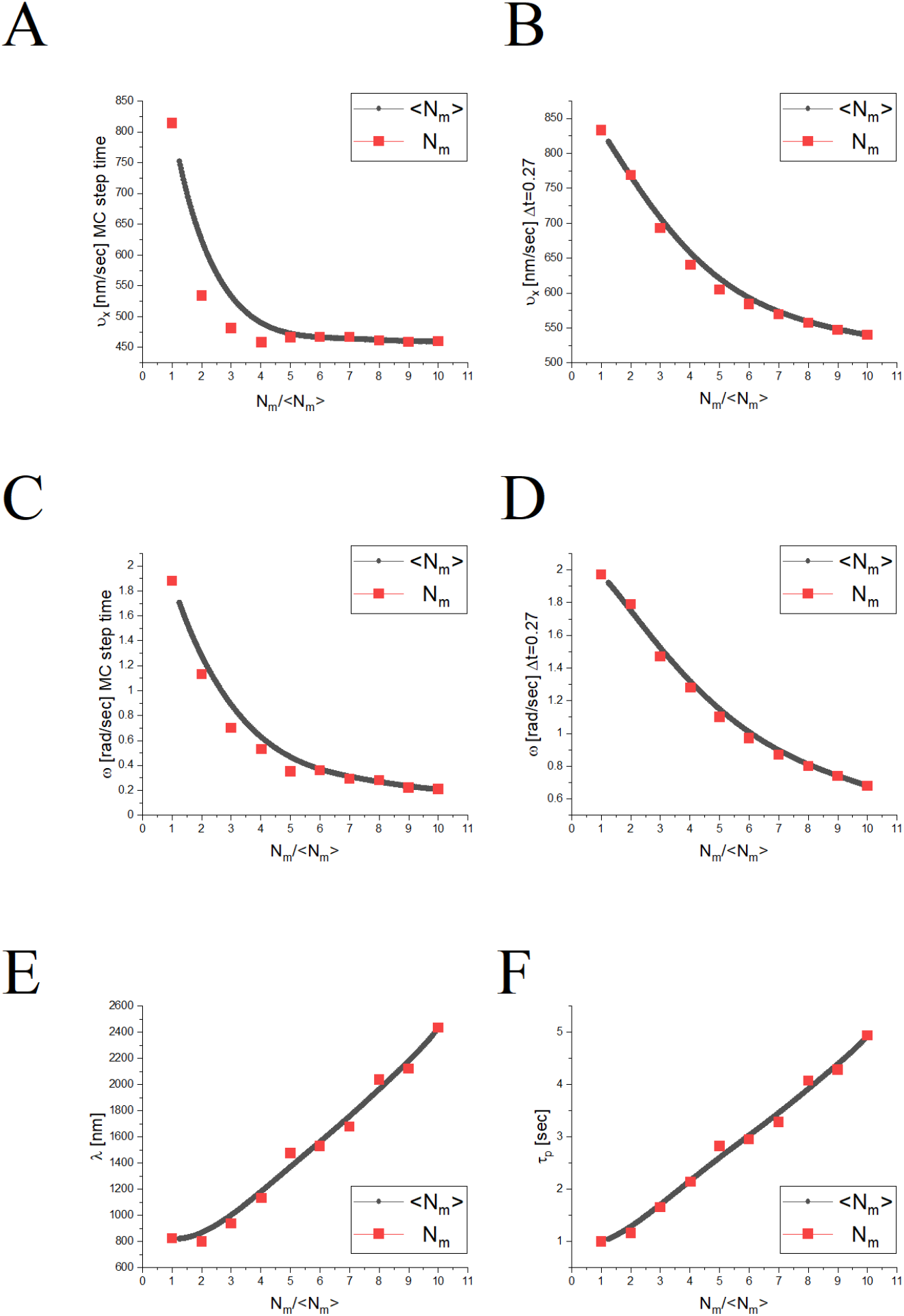
Same as Fig. S10.1 but for *N* = 10.

### Movies

**Movie 1** depicts the motion of NPs (green) on MTs (red) (system III). (Bar is 5 μm).

**Movie 2** depicts the motion of a NP (green) hopping between crossing MT tracks (red) (system III). (Bar is 5 μm).

**Movie 3** depicts the motion of a NP with a single NP-anchored motor (N_m_ = 1).

**Movie 4** depicts the motion of a NP with three NP-anchored motor (N_m_ = 3).

**Movie 5** depicts the motion of a NP with seven NP-anchored motor (N_m_ = 7).

**Movie 6** depicts the motion of a NP with 13 NP-anchored motor (N_m_ = 13).

## Footnotes

1 Using the NP translational Stokes drag coefficient, *γ_t_* = 6*πηR*, and velocity of 4 μm/s, leads to a drag force *f* = *γv*~10^-2^ pN, for *η* = 10 mPa s as an upper bound for the CE viscosity, that is much smaller with respect to the typical motor force 4 pN.

2 In any case, it should have only a relatively small effect on the motion of the NP. First, during the periods with simultaneous binding of more than one motor, it is very rare to find a motor at zero load. Second, during the periods of single motor binding, the only effect of the above mismatch is on the temporal velocity, which we adjust to the known value by fitting the step time to 0.0108 sec.

3 *d*_dynein_ is estimated as follows: The dynein structure, without the tail domain, can be said to consist of three major domains: MTBD, stalk, and AAA+ ring, with the approximated dimensions of 4,15, and 10 nm, respectively^14,15^. The length of the dynein (*d*_dynein_), more or less, equals to the length of the dynein heavy chain. It is known^16^ that two-thirds of the chain (C-terminal) spans from the MTBD to the AAA+ ring, and the remaining portion is the “tail” domain backbone (N-terminal). Thus, a simple estimation of *d*_dynein_ leads to a total length of about 45 nm.

4 *d*_Importins_ is taken as 15nm. However, it is a rough estimation obtained by using Chimera that lies between two conformations: dense and relaxed^17,18^.

## References

1. Hirokawa, N. Kinesin and dynein superfamily proteins and the mechanism of organelle transport. Science 279, 519–526 (1998).

2. Vale, R.D. The molecular motor toolbox for intracellular transport. Cell 112, 467–480 (2003).

3. Vaisberg, E.A., Koonce, M.P. & McIntosh, J.R. Cytoplasmic dynein plays a role in mammalian mitotic spindle formation. The Journal of cell biology 123, 849–858 (1993).

4. Gerson-Gurwitz, A. et al. Directionality of individual kinesin-5 Cin8 motors is modulated by loop 8, ionic strength and microtubule geometry. The EMBO journal 30, 4942–4954 (2011).

5. Hirokawa, N., Noda, Y., Tanaka, Y. & Niwa, S. Kinesin superfamily motor proteins and intracellular transport. Nature reviews Molecular cell biology 10, 682 (2009).

6. Hirose, K. Handbook of dynein, (Pan Stanford, 2012).

7. Salman, H. et al. Nuclear localization signal peptides induce molecular delivery along microtubules. Biophysical journal 89, 2134–2145 (2005).

8. Pilling, A.D., Horiuchi, D., Lively, C.M. & Saxton, W.M. Kinesin-1 and Dynein are the primary motors for fast transport of mitochondria in Drosophila motor axons. Molecular biology of the cell 17, 2057–2068 (2006).

9. Campbell, E.M. & Hope, T.J. HIV-1 capsid: the multifaceted key player in HIV-1 infection. Nature Reviews Microbiology 13, 471 (2015).

10. Jayappa, K.D., Ao, Z. & Yao, X. The HIV-1 passage from cytoplasm to nucleus: the process involving a complex exchange between the components of HIV-1 and cellular machinery to access nucleus and successful integration. International journal of biochemistry and molecular biology 3, 70 (2012).

11. Carnes, S.K. & Aiken, C. Host proteins involved in microtubule-dependent HIV-1 intracellular transport and uncoating. Future Virology 14, 361–374 (2019).

12. Dharan, A. & Campbell, E.M. Role of microtubules and microtubule-associated proteins in HIV-1 infection. Journal of virology 92, e00085–18 (2018).

13. McDonald, D. et al. Visualization of the intracellular behavior of HIV in living cells. The Journal of cell biology 159, 441–452 (2002).

14. Scherer, J. & Vallee, R.B. Adenovirus recruits dynein by an evolutionary novel mechanism involving direct binding to pH-primed hexon. Viruses 3, 1417–1431 (2011).

15. Kelkar, S.A., Pfister, K.K., Crystal, R.G. & Leopold, P.L. Cytoplasmic dynein mediates adenovirus binding to microtubules. Journal of virology 78, 10122–10132 (2004).

16. Cohen, O. & Granek, R. Nucleus-targeted drug delivery: theoretical optimization of nanoparticles decoration for enhanced intracellular active transport. Nano letters 14, 2515–2521 (2014).

17. Gennerich, A., Carter, A.P., Reck-Peterson, S.L. & Vale, R.D. Force-induced bidirectional stepping of cytoplasmic dynein. Cell 131, 952–965 (2007).

18. Can, S., Lacey, S., Gur, M., Carter, A.P. & Yildiz, A. Directionality of dynein is controlled by the angle and length of its stalk. Nature 566, 407 (2019).

19. Qiu, W. et al. Dynein achieves processive motion using both stochastic and coordinated stepping. Nature structural & molecular biology 19, 193 (2012).

20. Reck-Peterson, S.L. et al. Single-molecule analysis of dynein processivity and stepping behavior. Cell 126, 335–348 (2006).

21. Ori-McKenney, K.M., Xu, J., Gross, S.P. & Vallee, R.B. A cytoplasmic dynein tail mutation impairs motor processivity. Nature cell biology 12, 1228 (2010).

22. Cianfrocco, M.A., DeSantis, M.E., Leschziner, A.E. & Reck-Peterson, S.L. Mechanism and regulation of cytoplasmic dynein. Annual review of cell and developmental biology 31, 83–108 (2015).

23. Reck-Peterson, S.L., Redwine, W.B., Vale, R.D. & Carter, A.P. The cytoplasmic dynein transport machinery and its many cargoes. Nature Reviews Molecular Cell Biology 19, 382–398 (2018).

24. Gross, S.P., Vershinin, M. & Shubeita, G.T. Cargo transport: two motors are sometimes better than one. Current biology 17, R478–R486 (2007).

25. Vershinin, M., Carter, B.C., Razafsky, D.S., King, S.J. & Gross, S.P. Multiple-motor based transport and its regulation by Tau. Proceedings of the National Academy of Sciences 104, 87–92 (2007).

26. Hendricks, A.G., Holzbaur, E.L. & Goldman, Y.E. Force measurements on cargoes in living cells reveal collective dynamics of microtubule motors. Proceedings of the National Academy of Sciences 109, 18447–18452 (2012).

27. Grotjahn, D.A. et al. Cryo-electron tomography reveals that dynactin recruits a team of dyneins for processive motility. Nature structural & molecular biology 25, 203–207 (2018).

28. Torisawa, T. et al. Autoinhibition and cooperative activation mechanisms of cytoplasmic dynein. Nature cell biology 16, 1118–1124 (2014).

29. Mesika, A., Kiss, V., Brumfeld, V., Ghosh, G. & Reich, Z. Enhanced intracellular mobility and nuclear accumulation of DNA plasmids associated with a karyophilic protein. Human gene therapy 16, 200–208 (2005).

30. Rossi, L., Hohn, B. & Tinland, B. Integration of complete transferred DNA units is dependent on the activity of virulence E2 protein of Agrobacterium tumefaciens. Proceedings of the National Academy of Sciences 93, 126–130 (1996).

31. Peker, I. & Granek, R. Multimotor Driven Cargos: From Single Motor under Load to the Role of Motor-Motor Coupling. The Journal of Physical Chemistry B 120, 6319–6326 (2016).

32. Müller, M.J., Klumpp, S. & Lipowsky, R. Tug-of-war as a cooperative mechanism for bidirectional cargo transport by molecular motors. Proceedings of the National Academy of Sciences 105, 4609–4614 (2008).

33. Derr, N.D. et al. Tug-of-war in motor protein ensembles revealed with a programmable DNA origami scaffold. Science 338, 662–665 (2012).

34. Hancock, W.O. Bidirectional cargo transport: moving beyond tug of war. Nature reviews Molecular cell biology 15, 615 (2014).

35. Müller, M.J., Klumpp, S. & Lipowsky, R. Bidirectional transport by molecular motors: enhanced processivity and response to external forces. Biophysical journal 98, 2610–2618 (2010).

36. Can, S., Dewitt, M.A. & Yildiz, A. Bidirectional helical motility of cytoplasmic dynein around microtubules. Elife 3, e03205 (2014).

37. Elshenawy, M.M. et al. Cargo adaptors regulate stepping and force generation of mammalian dynein-dynactin. Nature chemical biology, 1–9 (2019).

38. DeWitt, M.A., Chang, A.Y., Combs, P.A. & Yildiz, A. Cytoplasmic dynein moves through uncoordinated stepping of the AAA+ ring domains. Science 335, 221–225 (2012).

39. Fayer, I. & Granek, R. Directional stepping model for yeast dynein: Longitudinal-and side-step distributions. Biophysical journal (2019).

40. TyleráMcLaughlin, R. Collective dynamics of processive cytoskeletal motors. Soft Matter 12, 14–21 (2016).

41. Berger, F., Keller, C., Lipowsky, R. & Klumpp, S. Elastic coupling effects in cooperative transport by a pair of molecular motors. Cellular and Molecular Bioengineering 6, 48–64 (2013).

42. Khataee, H. & Howard, J. Force generated by two kinesin motors depends on the load direction and intermolecular coupling. Physical review letters 122, 188101 (2019).

43. Miles, C.E. The University of Utah (2018).

44. Wang, Q. & Kolomeisky, A.B. Theoretical Analysis of Run Length Distributions for Coupled Motor Proteins. The Journal of Physical Chemistry B (2019).

45. Guérin, T., Prost, J., Martin, P. & Joanny, J.-F. Coordination and collective properties of molecular motors: theory. Current opinion in cell biology 22, 14–20 (2010).

46. Kolomeisky, A.B. & Fisher, M.E. Molecular motors: a theorist’s perspective. Annu. Rev. Phys. Chem. 58, 675–695 (2007).

47. Ando, J. et al. Small stepping motion of processive dynein revealed by load-free high-speed single-particle tracking. Scientific reports 10, 1–11 (2020).

48. Kunwar, A. & Mogilner, A. Robust transport by multiple motors with nonlinear force–velocity relations and stochastic load sharing. Physical biology 7, 016012 (2010).

49. Rai, A.K., Rai, A., Ramaiya, A.J., Jha, R. & Mallik, R. Molecular adaptations allow dynein to generate large collective forces inside cells. Cell 152, 172–182 (2013).

50. Belyy, V., Hendel, N.L., Chien, A. & Yildiz, A. Cytoplasmic dynein transports cargos via load-sharing between the heads. Nature communications 5, 1–9 (2014).

51. Ferro, L.S., Can, S., Turner, M.A., ElShenawy, M.M. & Yildiz, A. Kinesin and dynein use distinct mechanisms to bypass obstacles. Elife 8, e48629 (2019).

52. Rosano, C., Arosio, P. & Bolognesi, M. The X-ray three-dimensional structure of avidin. Biomolecular Engineering 16, 5–12 (1999).

53. De Gennes, P. Model polymers at interfaces. in Physical basis of cell-cell adhesion 39–60 (CRC Press Boca Raton, FL, 1988).

54. Klumpp, S., Keller, C., Berger, F. & Lipowsky, R. Molecular motors: Cooperative phenomena of multiple molecular motors. in Multiscale Modeling in Biomechanics and Mechanobiology 27–61 (Springer, 2015).

55. Yildiz, A. & Canty, J.T. Private Communication. (2019).

56. Crocker, J.C. & Grier, D.G. Methods of digital video microscopy for colloidal studies. Journal of colloid and interface science 179, 298–310 (1996).

57. Savin, T. & Doyle, P.S. Statistical and sampling issues when using multiple particle tracking. Physical Review E 76, 021501 (2007).

58. Ananthanarayanan, V. Harvard Medical School, Boston (1986).

59. Bieling, P., Telley, I.A., Piehler, J. & Surrey, T. Processive kinesins require loose mechanical coupling for efficient collective motility. EMBO reports 9, 1121–1127 (2008).

60. Böhm, K., Stracke, R. & Unger, E. Speeding up kinesin-driven microtubule gliding in vitro by variation of cofactor composition and physicochemical parameters. Cell biology international 24, 335–341 (2000).

61. Lippert, L.G. et al. Angular measurements of the dynein ring reveal a stepping mechanism dependent on a flexible stalk. Proceedings of the National Academy of Sciences 114, E4564–E4573 (2017).

62. Li, L., Alper, J. & Alexov, E. Cytoplasmic dynein binding, run length, and velocity are guided by long-range electrostatic interactions. Scientific reports 6, 31523 (2016).

63. Belyy, V. et al. The mammalian dynein–dynactin complex is a strong opponent to kinesin in a tug-of-war competition. Nature cell biology 18, 1018 (2016).

64. Bameta, T., Padinhateeri, R. & Inamdar, M.M. Force generation and step-size fluctuations in a dynein motor. Journal of Statistical Mechanics: Theory and Experiment 2013, P02030 (2013).

65. Roberts, A.J. et al. AAA+ Ring and linker swing mechanism in the dynein motor. Cell 136, 485–495 (2009).

66. Elshenawy, M.M. et al. Cargo adaptors regulate stepping and force generation of mammalian dynein-dynactin. Nature chemical biology 15, 1093–1101 (2019).

67. Fayer, I. & Granek, R. Directional stepping model for yeast dynein: Longitudinal-and side-step distributions. Biophysical Journal 117, 1892–1899 (2019).

68. Dohner, K. et al. Function of dynein and dynactin in herpes simplex virus capsid transport. Molecular biology of the cell 13, 2795–2809 (2002).

69. Mesika, A., Grigoreva, I., Zohar, M. & Reich, Z. A regulated, NFκB-assisted import of plasmid DNA into mammalian cell nuclei. Molecular Therapy 3, 653–657 (2001).

70. Ding, H.m. & Ma, Y.q. Theoretical and computational investigations of nanoparticle–biomembrane interactions in cellular delivery. Small 11, 1055–1071 (2015).

71. Vankayala, R., Kuo, C.L., Nuthalapati, K., Chiang, C.S. & Hwang, K.C. Nucleus-targeting gold nanoclusters for simultaneous in vivo fluorescence imaging, gene delivery, and NIR-light activated photodynamic therapy. Advanced Functional Materials 25, 5934–5945 (2015).

72. Fu, M.-m. & Holzbaur, E.L. JIP1 regulates the directionality of APP axonal transport by coordinating kinesin and dynein motors. Journal of Cell Biology 202, 495–508 (2013).

73. Kalderon, D., Roberts, B.L., Richardson, W.D. & Smith, A.E. A short amino acid sequence able to specify nuclear location. Cell 39, 499–509 (1984).

74. Danino, D., Bernheim-Groswasser, A. & Talmon, Y. Digital cryogenic transmission electron microscopy: an advanced tool for direct imaging of complex fluids. Colloids and Surfaces A: Physicochemical and Engineering Aspects 183, 113–122 (2001).

75. Delgado, Á.V., González-Caballero, F., Hunter, R., Koopal, L. & Lyklema, J. Measurement and interpretation of electrokinetic phenomena. Journal of colloid and interface science 309, 194–224 (2007).

76. Shemesh, A., Ginsburg, A., Levi-Kalisman, Y., Ringel, I. & Raviv, U. Structure, assembly, and disassembly of tubulin single rings. Biochemistry 57, 6153–6165 (2018).

77. Gell, C. et al. Microtubule dynamics reconstituted in vitro and imaged by single-molecule fluorescence microscopy. in Methods in cell biology, Vol. 95 221–245 (Elsevier, 2010).

78. Phelps, K. & Walker, R. NEM tubulin inhibits microtubule minus end assembly by a reversible capping mechanism. Biochemistry 39, 3877–3885 (2000).

79. Hyman, A. Preparation of marked microtubules for the assay of the polarity of microtubule-based motors by fluorescence. J Cell Sci 1991, 125–127 (1991).

80. Bickel, T., Jeppesen, C. & Marques, C. Local entropic effects of polymers grafted to soft interfaces. The European Physical Journal E 4, 33–43 (2001).

81. DiMarzio, E.A. Proper accounting of conformations of a polymer near a surface. The Journal of Chemical Physics 42, 2101–2106 (1965).

82. Lee, H., Venable, R.M., MacKerell Jr, A.D. & Pastor, R.W. Molecular dynamics studies of polyethylene oxide and polyethylene glycol: hydrodynamic radius and shape anisotropy. Biophysical journal 95, 1590–1599 (2008).

83. Kienberger, F. et al. Static and dynamical properties of single poly (ethylene glycol) molecules investigated by force spectroscopy. Single Molecules 1, 123–128 (2000).

84. Lacey, S.E., He, S., Scheres, S.H. & Carter, A.P. Cryo-EM of dynein microtubule-binding domains shows how an axonemal dynein distorts the microtubule. Elife 8(2019).

85. Israelachvili, J.N. Intermolecular and surface forces, (Academic press, 2011).

86. Oesterhelt, F., Rief, M. & Gaub, H. Single molecule force spectroscopy by AFM indicates helical structure of poly (ethylene-glycol) in water. New Journal of Physics 1, 6 (1999).

87. De Gennes, P.-G. & Gennes, P.-G. Scaling concepts in polymer physics, (Cornell university press, 1979).

88. Doi, M. & Edwards, S.F. The theory of polymer dynamics, (oxford university press, 1988).

89. Rubinstein, M. & Colby, R.H. Polymer physics, (Oxford university press New York, 2003).

90. Dobrinski, H., Wilkens, M., Benecke, W. & Binder, J. Flexible microfluidic-device-stamp-system with integrated electrical sensor for real time DNA detection. in 1st Annual International IEEE-EMBS Special Topic Conference on Microtechnologies in Medicine and Biology. Proceedings (Cat. No. 00EX451) 33–35 (IEEE, 2000).

91. Kon, T. et al. The 2.8 A crystal structure of the dynein motor domain. Nature 484, 345–50 (2012).

92. Kon, T., Mogami, T., Ohkura, R., Nishiura, M. & Sutoh, K. ATP hydrolysis cycle-dependent tail motions in cytoplasmic dynein. Nature structural & molecular biology 12, 513 (2005).

93. Biagioni, A. et al. Delivery systems of CRISPR/Cas9-based cancer gene therapy. Journal of Biological Engineering 12, 1–9 (2018).

94. Cingolani, G., Bednenko, J., Gillespie, M.T. & Gerace, L. Molecular basis for the recognition of a nonclassical nuclear localization signal by importin beta. Mol Cell 10, 1345–53 (2002).

95. Fayer, I. et al., to be published.

## SI References

1. Danino, D., Bernheim-Groswasser, A. & Talmon, Y. Digital cryogenic transmission electron microscopy: an advanced tool for direct imaging of complex fluids. Colloids and Surfaces A: Physicochemical and Engineering Aspects 183, 113–122 (2001).

2. Rosano, C., Arosio, P. & Bolognesi, M. The X-ray three-dimensional structure of avidin. Biomolecular Engineering 16, 5–12 (1999).

3. Israelachvili, J.N. Intermolecular and surface forces, (Academic press, 2011).

4. De Gennes, P. Model polymers at interfaces. in Physical basis of cell-cell adhesion 39–60 (CRC Press Boca Raton, FL, 1988).

5. Delgado, Á.V., González-Caballero, F., Hunter, R., Koopal, L. & Lyklema, J. Measurement and interpretation of electrokinetic phenomena. Journal of colloid and interface science 309, 194–224 (2007).

6. Oesterhelt, F., Rief, M. & Gaub, H. Single molecule force spectroscopy by AFM indicates helical structure of poly (ethylene-glycol) in water. New Journal of Physics 1, 6 (1999).

7. De Gennes, P.-G. & Gennes, P.-G. Scaling concepts in polymer physics, (Cornell university press, 1979).

8. Doi, M. & Edwards, S.F. The theory of polymer dynamics, (oxford university press, 1988).

9. Rubinstein, M. & Colby, R.H. Polymer physics, (Oxford university press New York, 2003).

10. Lee, H., Venable, R.M., MacKerell Jr, A.D. & Pastor, R.W. Molecular dynamics studies of polyethylene oxide and polyethylene glycol: hydrodynamic radius and shape anisotropy. Biophysical journal 95, 1590–1599 (2008).

11. Kienberger, F. et al. Static and dynamical properties of single poly (ethylene glycol) molecules investigated by force spectroscopy. Single Molecules 1, 123–128 (2000).

12. Dobrinski, H., Wilkens, M., Benecke, W. & Binder, J. Flexible microfluidic-device-stamp-system with integrated electrical sensor for real time DNA detection. in 1st Annual International IEEE-EMBS Special Topic Conference on Microtechnologies in Medicine and Biology. Proceedings (Cat. No. 00EX451) 33–35 (IEEE, 2000).

13. Crocker, J.C. & Grier, D.G. Methods of digital video microscopy for colloidal studies. Journal of colloid and interface science 179, 298–310 (1996).

14. Lacey, S.E., He, S., Scheres, S.H. & Carter, A.P. Cryo-EM of dynein microtubule-binding domains shows how an axonemal dynein distorts the microtubule. Elife 8 (2019).

15. Kon, T. et al. The 2.8 A crystal structure of the dynein motor domain. Nature 484, 345–50 (2012).

16. Kon, T., Mogami, T., Ohkura, R., Nishiura, M. & Sutoh, K. ATP hydrolysis cycle–dependent tail motions in cytoplasmic dynein. Nature structural & molecular biology 12, 513 (2005).

17. Biagioni, A. et al. Delivery systems of CRISPR/Cas9-based cancer gene therapy. Journal of Biological Engineering 12, 1–9 (2018).

18. Cingolani, G., Bednenko, J., Gillespie, M.T. & Gerace, L. Molecular basis for the recognition of a nonclassical nuclear localization signal by importin beta. Mol Cell 10, 1345–53 (2002).

